# Dynamic holocentric genomes facilitate divergent evolutionary paths through chromosomal rearrangements, hybrid dysfunction, and recombination suppression

**DOI:** 10.64898/2026.07.21.739116

**Authors:** Rogelio Sánchez-Villegas, Ashwini V. Mohan, Inés Gómez-Ramos, Modesto Luceño, Mónica Míguez, Enrique Maguilla, José Ignacio Márquez-Corro, Diego Herrero-Doblado, Nafiseh Sargheini, André Marques, Kay Lucek, Marcial Escudero, Santiago Martín-Bravo

## Abstract

Theory predicts chromosomal rearrangements (CRs) to promote reproductive isolation and local adaptation by disrupting meiosis and altering recombination landscapes. These processes are especially important in holocentric organisms, whose diffuse centromeres facilitate CRs. Here, we investigated the genomic origins and evolutionary consequences of CRs in the holocentric sedge *Carex laevigata*, a species with extreme intraspecific chromosome-number variation (2n = 69-84). Establishing chromosome-scale genome assemblies, experimental crosses involving more than one thousand living plants throughout three generations and eight years, linkage mapping, and Quantitative Trait Loci (QTL) analyses, we identified extensive CRs among karyotypically distinct populations. Breakpoint regions of CRs were enriched in GC-rich and repetitive sequences, particularly LTR-Gypsy elements, suggesting recurrent genomic regions prone to structural instability. Inter-cytotype hybrids formed complex meiotic configurations and showed reduced germination success, consistent with hybrid dysfunction associated with increasing chromosomal divergence. Rearranged chromosomes exhibited strong recombination suppression and segregation distortion near breakpoint regions and within inverted segments. QTL analyses further identified fitness-related loci associated with both rearranged and collinear chromosomes. Together, our results corroborate theoretical predictions providing novel empirical evidence that CRs arise preferentially in structurally fragile genomic regions and contribute to genomic divergence through hybrid dysfunction and recombination suppression.

## Introduction

The eukaryotic genome is far from being a static blueprint; but rather a dynamic landscape constantly reshaped by structural upheavals. Chromosomal rearrangements (CRs) have emerged as key contributors to this diversity across the Tree of Life. Since the discovery of the first rearrangements including Robertsonian translocations in grasshoppers (Robertson, 1916) and inversions in *Drosophila* (Sturtevant, 1921), CRs have been recognized not merely as structural anomalies, but as fundamental drivers of evolution that can dictate the pace and direction of diversification (Berdan et al., 2023; Lucek et al., 2023; Benítez-Benítez et al., 2025).

Fusions, fissions, translocations and inversions are CRs that do not entail changes in genome size, but may or not result in changes in chromosome number (Benítez-Benítez et al., 2025). They originate primarily from DNA double-strand breaks (DSBs) that are imperfectly repaired, leading to changes in the linear order and orientation of chromosomal segments (Carvalho and Lupski, 2016; Burssed et al., 2022). Such breaks are not distributed randomly; instead, they are often concentrated in ‘fragile’ regions governed by the local genomic environment (Luceño, 1994; Wright and Schaeffer, 2022; Escudero et al., 2024). Key determinants of such instability include nucleotide composition, specifically GC-rich regions, which are prone to higher mutation rates and recombination initiation (Fullerton et al., 2001), and the presence of repetitive elements such as transposable elements (TEs), which can facilitate ectopic recombination (Robberecht et al., 2013; Farré et al., 2015).

A major, long-standing challenge lies in understanding how diversity in CRs translates into reproductive isolation and ultimately speciation. Two key mechanisms have been proposed through which CRs can drive these processes. First, by causing hybrid dysfunction: hybrids between individuals with different karyotypes often exhibit reduced fitness leading to strong selection against heterokaryotypes and reinforcing reproductive isolation (Coyne and Orr, 2004). However, CRs fixation and role in speciation can be paradoxical: strongly underdominant rearrangements are rarely fixed in populations, whereas weakly underdominant ones are more easily to become fixed but contribute little to reproductive isolation and, consequently, to speciation (Faria and Navarro, 2010). Secondly, CRs can profoundly influence the recombination landscape across the genome. Hybrids between different karyotypes often exhibit reduced recombination in rearranged chromosomal regions, which can act as barriers to gene flow between populations (Rieseberg, 2001). When rearranged regions harbor locally adaptive allelic combinations, they can act as so called supergenes, maintaining co-adapted gene complexes and thereby promote differentiation among karyotypes through divergent selection (Ayala and Coluzzi, 2005). Beyond these mechanisms, other factors such as alterations in three-dimensional genome architecture or epigenetic modifications induced by CRs may also contribute to differentiation and speciation among karyotypes (Spielmann et al., 2018; Vara et al., 2021; Mohan et al., 2024). Holocentric organisms display several peculiarities that can influence how CRs behave during cell division, in stark contrast with monocentric lineages. Holocentric chromosomes possess kinetochore activity distributed along their entire length rather than being restricted to a single centromere (Hofstatter et al., 2022a). Whereas acentric fragments or dicentric chromosomes produced by non-Robertsonian fissions or fusions would normally be lost or missegregated in monocentric species, such rearrangements can still segregate properly in holocentric species, as their kinetochoric activity remains intact (Wahl, 1940; Luceño, 1993; Zedek and Bureš, 2018; Márquez-Corro et al., 2019). Moreover, several holocentric taxa exhibit inverted meiosis, in which sister chromatids segregate during the first meiotic division and homologous chromosomes during the second one (Faulkner, 1972; Márquez-Corro et al., 2019, Marques and Dinnenberg, 2025). This mechanism especially facilitates the proper pairing and segregation of multivalent configurations (Lukhtanov et al., 2018). Together, these features reduce the degree of underdominance associated with CRs, at least in heterozygotes between closely related karyotypes within species (Lucek et al., 2022). This may account for the great variation in chromosome number displayed by some holocentric lineages, such as Lepidoptera (butterflies and moths), the largest holocentric group (2n = 10-446; Lukhtanov, 2015), and the plant family Cyperaceae (2n = 4-226; Márquez-Corro et al., 2021; Shafir et al., 2023). Holocentricity therefore plays a key role in generating and maintaining biodiversity by chromosomal evolution (de Vos et al., 2020; Márquez-Corro et al., 2021; Tribble et al., 2025), although large scale comparative studies remain still limited (Márquez-Corro et al., 2018, Lucek et al. 2023).

Holocentricity also shapes recombination dynamics as the number of crossovers per chromosome is more limited in holocentric organisms, being typically one and maximal two (Nokkala et al., 2004; Zedek et al., 2026). Genome-wide recombination rate during meiosis is therefore directly proportional to only chromosome number in holocentric organisms, whereas in monocentric ones it also depends on the average number of chiasmata per chromosome (Zedek et al., 2026). In monocentric organisms, recombination typically decreases toward the centromere and increases near the telomeres (Johnston, 2024). In contrast, holocentric species show more diverse patterns, where the nematode *Caenorhabditis elegans* resemble monocentric patterns due to local clustering of holocentromeres (Rockman and Kruglyak, 2009), whereas others, like Lepidoptera and several holocentric plants, display more uniform or variable distributions compared to the typical monocentric pattern (Haenel et al., 2018; Hofstatter et al., 2022b; Castellani et al., 2024).

Holocentricity has evolved at least 20 times in eukaryotes (Melters et al., 2012; Escudero et al., 2016a), representing 15-20% of the extant eukaryotic species diversity (Márquez-Corro et al., 2018). Among these lineages, the family Cyperaceae is the most species rich holocentric plant group, including more than 5,500 species (POWO, 2025). *Carex* is by far the largest genus of the family, with more than 2,000 species (Martín-Bravo et al., 2019; POWO, 2025). *Carex* exhibits extraordinary karyotype variability, not only among species (2n = 10-132; Roalson et al., 2008; Márquez-Corro et al., 2021), but also among populations of the same species with ∼40% of all studied species showing chromosome number variability (e.g. *C. scoparia* with 2n = 56-70, Escudero et al., 2013a; or *C. helodes* with 2n = 68-75, Escudero et al., 2008; 2023). Specifically, CRs entailing changes in chromosome number but not in genome size (dysploidy; Ehrendorfer, 1964; Dyer, 1979; Escudero et al., 2014), have played a major role in the macroevolutionary diversification of the genus (Márquez-Corro et al., 2021; Tribble et al., 2025). Despite the capacity of holocentric chromosomes to tolerate CRs during pairing and segregation, hybrid dysfunction has been experimentally documented between different chromosomal races of a few *Carex* species (Escudero et al., 2016b;Whitkus, 1988). The consequences of CRs for recombination remain less explored in *Carex*, although recent work in *C. scoparia* has shown that fissions and fusions can lead to suppressed recombination (Escudero et al., 2023). However, whether recombination suppression occurs in genomic regions underlying local adaptation remains unexplored.

Here, we present an integrative analysis of *Carex laevigata*, investigating both the genomic origins and the evolutionary consequences of CRs within a holocentric species. Characterized by some of the highest dysploid variation within the genus *Carex* (2n=69 to 84; Fig. 1b), this species has emerged as a model system for intra-specific chromosomal evolution over 30 years of research (Luceño & Castroviejo, 1991; Escudero et al., 2013b; Escudero et al., 2015; Márquez-Corro et al., 2024). Building on the hypothesis that CRs preferentially arise within architecturally ‘fragile’ regions (Farré et al., 2015), we first characterized the genomic landscape of rearrangement breakpoints. Specifically, we evaluated the association between these breakpoints and various genomic features, including GC content, repetitive element composition, and the spatial distribution of coding sequences. We then assessed the evolutionary impact of these rearrangements. Through controlled crosses between individuals from distinct karyotypic populations, we generated parental selfed lines, F hybrids (from interpopulation crosses), and F offspring (from selfed F individuals). Germination success (as the proportion of germinated seeds out of the total) allowed us to quantify the fitness consequences of structural heterozygosity (i.e. hybrid dysfunction) for both F s and F s. Using genomic data from F s, we constructed a genetic linkage map to test whether and to which degree CRs suppress recombination. Finally, we evaluated fitness-related morphological traits in F individuals and performed QTL mapping to determine whether rearranged regions coincide with loci likely associated with local adaptation. Together, this integrative framework links the mechanistic basis of chromosomal fragility with its consequences for recombination, fitness, and population divergence.

**Figure 1.**
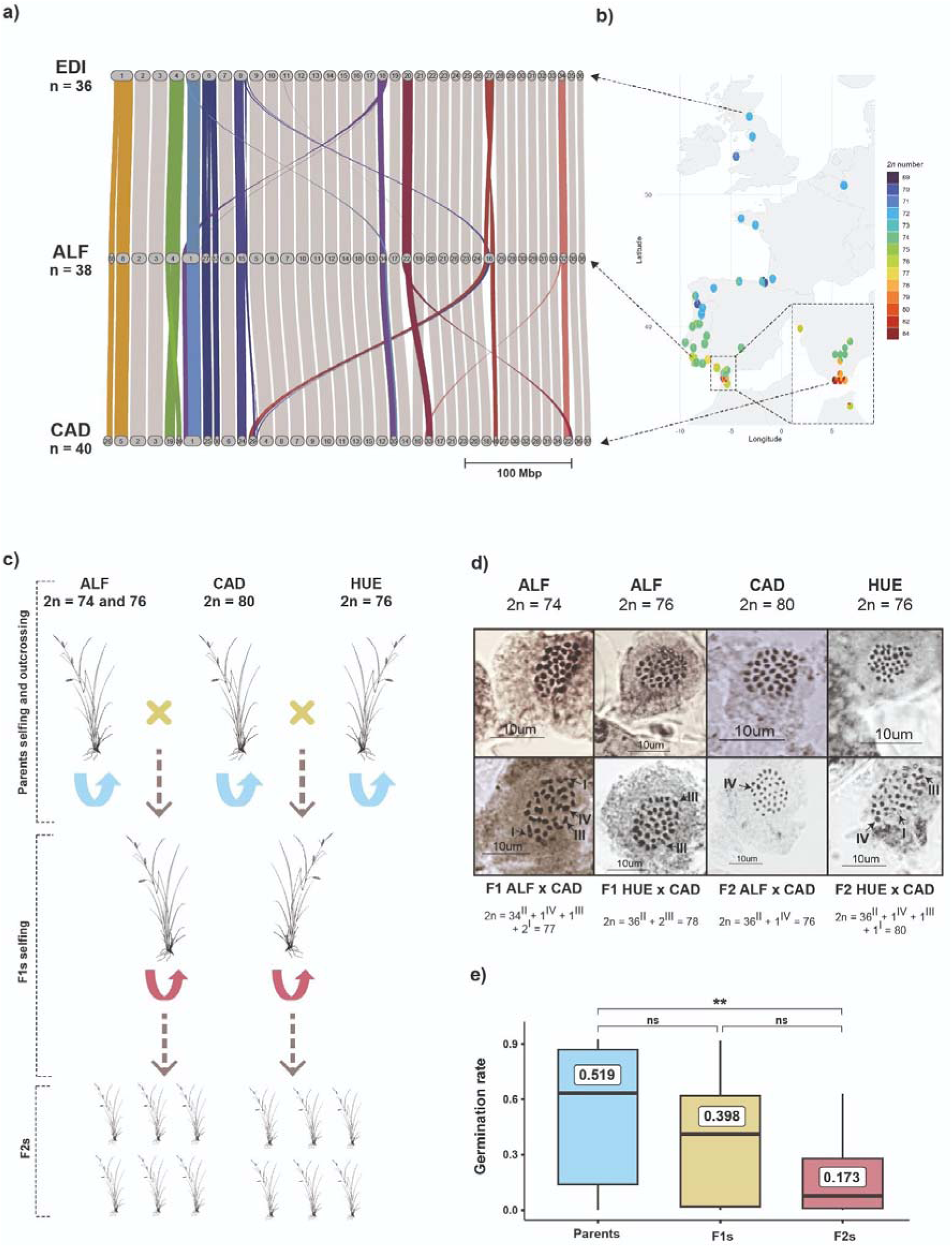
Karyotypic variation, chromosomal rearrangements and hybrid fitness. **a)** Synteny plot of the EDI genome and the parental genomes ALF and CAD. Each genome is shown as a single horizontal line divided into chromosomes (numbered blocks). Syntenic regions are depicted as braids connecting homologous regions across genomes. Colored braids highlight chromosomes that show chromosomal rearrangements (CRs) relative to the other genomes, while grey braids indicate non-rearranged chromosomes. **b)** The distribution of *C. laevigata* karyotypes is shown across populations. Chromosome number, represented by colours, were obtained from counts by Davies (1956), Löve (1972), Luceño and Castroviejo (1991), Escudero et al. (2013) and Márquez-Corro et al. (2024). Different chromosome numbers found in a population are represented as pie chart proportions. Dashed arrows indicate the populations from which the genomes were obtained. **c)** Schematic representation of the crosses performed in the greenhouse to obtain the F populations. **d)** Karyotypes of the parental lines used in the crosses and representative F and F_2_ hybrids. Arrows indicate non-bivalent configurations observed during meiosis, including univalents (^I^), trivalents (^III^), and tetravalents (^IV^). **e)** Boxplots showing germination success resulting from the different cross types (Parents, F s and F_2_s). The line and value within each box indicate the mean value. Pairwise statistical differences among germination success for each cross type are indicated above boxes (Kruskal–Wallis test followed by pairwise Wilcoxon tests with Benjamini–Hochberg correction). Significance codes: *p* < 0.01 (**), *ns* = not significant.

## Material and Methods

### Study species

*C. laevigata* (*Carex* clade Spirostachyae, Cyperaceae) is distributed predominantly across the Palearctic Atlantic region, with populations extending from Western Europe to northern Morocco, typically occurring in wet meadows and riparian woodlands at low elevations (0 to 1,200 m) (Luceño and Escudero, 2008; Luceño et al., 2023). The species shows a latitudinal gradient in chromosome number (Luceño and Castroviejo, 1991; Escudero et al., 2013; Fig. 1b). Chromosome counts of 2n = 72 are most frequent in northern Spain, northern Portugal, and elsewhere in Europe; 2n = 74 predominates in central Portugal; 2n = 76 in southern Portugal; and 2n = 78-84 are reported from southern Spain. Notably, Escudero et al. (2013) found that climatic variables were strong predictors of chromosome number variation, beyond geographic location or phylogeographic structure, suggesting the potential influence of local adaptation. Landscape genomics analyses further showed a correlation between variation in chromosome number and several bioclimatic variables (temperature seasonality, minimum temperature of the coldest month, precipitation of wettest quarter and precipitation of driest quarter), supporting the hypothesis that distinct *C. laevigata* karyotypes are locally adapted (Márquez-Corro et al., 2024).

### Sampling, experimental crosses and germination success estimation

In early 2019, individuals of *C. laevigata* were collected from three populations from South Iberian Peninsula with different chromosome number and established under greenhouse conditions at the Pablo de Olavide University. These individuals were subsequently used to perform controlled experimental crosses among populations (Fig. 1a). Two inter-population crosses were performed: one between a parent from Los Alcornocales, Cádiz, Spain (2n = 80; hereafter CAD) and a parent from Alferce, Portugal (2n = 74-76; hereafter ALF), and another between the same CAD parent and a parent from Doñana National Park, Huelva, Spain (2n = 76; hereafter HUE) to generate a set of F individuals. By late 2019, seeds from both crosses were harvested and germinated under sterile conditions in a POL-EKO ST1 germinator following the same procedure as for *C. helodes* in Narbona et al. (2013). Germinated seedlings were then transplanted into pots and grown under greenhouse conditions. In spring 2021, once the F plants reached reproductive maturity, we carried out self-fertilizations of the F s as well as the original parental individuals to obtain a set of selfed parental lines and a set of selfed F_1_s (from now on referred to as F_2_). As in the previous generation, seeds were germinated under sterile conditions and seedlings were cultivated in the greenhouse until spring 2023. To assess whether differences in chromosome number among parentals lead to hybrid dysfunction, we used seed germination success as a proxy for fitness. Germination success was calculated as the proportion of germinated seeds over the total number of collected seeds, and compared between seeds from self-fertilized parental lines, from inter-population crosses to generate F s and from selfed F s to generate the F_2_s. Fresh leaf tissue from the parents, the F individuals, and all available F progeny was collected and preserved in silica gel prior to DNA extraction. In addition, voucher specimens of all individuals involved in the crosses were prepared and deposited in the UPOS herbarium.

### Cytogenetic study

The parents and some of the F and F_2_ individuals from both crosses were karyotyped. When they started flowering, anthers were fixed and stained following Luceño (1988), and chromosomes were observed in metaphase I (MI) of meiosis using a Nikon Eclipse E400 microscope equipped with a digital camera Nikon DXM1200F.

### Genome sequencing and assembly

The genomes of the two karyotypically most different parents (CAD 2n = 80 and ALF 2n = 74) were sequenced and assembled. Unfortunately, the HUE parent died before its genome could be sequenced. To overcome this limitation, we sequenced and assembled the genome of one of the resulting F hybrids from the CAD × HUE cross and generated the haploid phased assemblies (hap) in order to reconstruct both the CAD and the missing HUE parental genomes. All genomes were sequenced using Pacific Biosciences single-molecule HiFi long reads combined with short-read Hi-C chromatin-interaction mapping data. Long-read sequencing was performed on a PacBio Revio (Pacific Biosciences, Menlo Park, CA, USA) by BGI Genomics (Hong Kong, China) from fresh leaves, flash frozen with liquid nitrogen and kept at -80°C until DNA extraction. For the Hi-C, nuclei extraction was carried out using the CelLytic™ PN Isolation/Extraction Kit from Merck (Merck KGaA, Germany) using fresh leaf tissue. DNA crosslinking, ligation, amplification, and library preparation was carried out using the Arima Hi-C high-throughput kit and a modified protocol for plant tissue (Escuer et al. submitted). The prepared Hi-C libraries were sequenced on an Illumina Novaseq X with 150 bps paired-end reads.

HiFi data quality was assessed using *NanoPlot* v1.20.0 (De Coster and Rademakers, 2023). K-mer histograms were generated with *Jellyfish* v2.3.1 (Marcais and Kingsford, 2011) and analyzed with *GenomeScope* v1.0 (Vurtre et al., 2017). Hi-C reads were cleaned following the Arima Hi-C mapping pipeline (Juric et al., 2019).

HiFi data was assembled using *HifiASM* v0.25.0 (Cheng et al., 2021), producing contig level assemblies. Duplicate purging was carried out through an initial purge within *HifiASM* followed by an additional round of haplotig removal using *purge_dups* v1.2.6 (Guan et al., 2020). Then, Hi-C data was mapped on the assembly using the Arima Hi-C mapping pipeline (Juric et al., 2019) implementing *bwa* 0.7.15 (Li and Durbin, 2009), *fastp* 0.23.5 (Chen et al., 2018) and *Picard* 1.141 (http://broadinstitute.github.io/picard). Scaffolding was then performed with *YaHS* v1.2a.2 (Zhou et al., 2023) followed by manual curation in *JuiceBox* 2.15 (Durand et al., 2016) correcting misalignments between scaffolds.

Assembly quality was evaluated by measuring contiguity with *gfastats* 1.3.11 (Formenti et al., 2022) and completeness with *CompleASM* v0.2.27 (Huang et al., 2023), using the Poales_odb12 BUSCO database. Haplotype-specific accuracy was assessed through k-mer based analysis with *Merqury* v1.3 (Rhie et al., 2020). Contaminant scaffolds, including organellar sequences, microbial fragments, and short low-coverage scaffolds, were identified and removed using *Tiara* v1.0.3 (Karlicki et al., 2022). Telomeric repeats were subsequently identified and visualized to determine the number of telomere-to-telomere scaffolds and to guide the removal of small residual scaffolds using TIDK (Brown et al., 2025).

Repetitive elements were annotated using the Earl Grey v7.0.2 pipeline (Baril et al., 2024). The resulting repeat library was then employed to soft-mask the genome assembly. Following masking, gene annotation was performed using BRAKER3 (Bruna et al., 2021) using the OrthDB poales_odb10 protein database (Kriventseva et al., 2018).

We also downloaded the publicly available *C. laevigata* genome from an individual collected near Edinburgh, Scotland with a karyotype of 2n = 72 from GenBank (Bioproject PRJEB78975, Darwin Tree of Life Project Consortium, 2022, 2024; further referred to as EDI) for the comparative genomic analyses. Organellar scaffolds were removed from this assembly, and the genome was annotated following the same workflow used for our newly generated assemblies.

### Synteny analysis

Comparative synteny analysis was conducted using GENESPACE v1.3.1 (Lovell et al., 2022) for the four *C. laevigata* genomes available, using the previously published genome (EDI) as a reference and the others ordered by increasing chromosome number which also matches the previously reported north to south gradient (ALF, CAD × HUE hap 1 and 2, and CAD). Protein-coding gene annotations and genome assemblies were parsed using the *parse_annotations* function to generate standardized FASTA and BED files with matching gene identifiers. Pairwise and multiple-genome collinearity blocks were then identified with the *run_genespace* function. In cases involving reciprocal translocations, chromosome fragments were oriented by prioritizing the largest collinear segments to ensure consistent alignment across genomes. Following reorientation, genomes were re-annotated using BRAKER3 and reanalyzed with GENESPACE to generate the final riparian plots.

### Genome features visualization and enrichment analysis

Genomic features were analyzed in non-overlapping 50-kb windows across all chromosomes of the EDI, ALF, and CAD assemblies using *BEDtools* v2.31.1 (Quinlan and Hall, 2010). The haplotypes from F CAD × HUE assemblies were not included in this analysis, as they correspond to a haploid phased reconstruction of a hybrid genome, which does not represent a true haploid chromosome set from either parental lineage. For each window, the proportion of coding sequence (CDS), total repeats and GC were quantified. For each genomic feature, we generated genome-wide profiles using scatterplots of window values and LOESS smoothing (span = 0.3) with *ggplot2* (Wickham, 2016) in *R v4.4.2* (R Core Team, 2021). To evaluate relationships among genomic features, pairwise correlation analyses were conducted using the *R* packages *GGally* (Schloerke et al., 2025). Associations between variables were assessed using Spearman’s rank correlation coefficient.

An enrichment analysis was performed with *regioneR* v1.4.0 (Gel et al., 2016) to test whether breakpoint regions were enriched or depleted for specific genomic features. Breakpoint intervals were obtained from the GENESPACE output and were considered as genomic regions marking transitions in syntenic collinearity between chromosomes of different genomes, including regions associated with CRs. Extracted breakpoints were visually verified with the HiC maps of the assembled parental genomes (Supplementary Fig. 2). The genomes are highly contiguous with 54 contigs in CAD and 59 contigs ALF and therefore, breakpoints are not artefacts of genome assembly. For each genome (EDI, ALF, and CAD), four primary feature datasets were compiled: coding sequences (CDS), GC-content (50-kb), total repeat elements, and individual repeat families identified by *Earl Grey*. GC values were treated as a continuous variable and represented as a *GRanges* object. Breakpoint intervals for each genome were intersected with each feature dataset using permutation-based tests implemented in *regioneR*. Two complementary analyses were performed: one including chromosomal ends when they were involved in CRs, and another excluding them to minimize potential telomeric effects on the statistical power of the analysis. Enrichment of discrete genomic features (CDS, total repeats, and each repeat family) was evaluated using the *numOverlaps* function, whereas GC content was tested using the *meanInRegions* function. In all cases, significance was assessed using 1,000 random permutations of breakpoint regions across the genome with *randomizeRegions*, maintaining the width of each breakpoint interval and using chromosome size as the genomic background. For each test, *regioneR* computed the observed statistic (number of overlaps or mean value), the null distribution from permutations, the empirical *p*-value, and the associated *z*-score. This framework allowed us to assess not only genome-wide patterns (e.g., CDS or GC enrichment) but also whether specific repeat superfamilies were associated with breakpoint regions.

In addition to the global enrichment tests, a fine-grained analysis was performed to evaluate the genomic environment of each individual breakpoint independently. For this purpose, each breakpoint interval was treated as a separate query region and compared against the genomic background using the same permutation framework. This approach allowed us to verify whether global genomic trends were consistently represented at the local scale and to identify specific breakpoints significantly associated with unique genomic features. Furthermore, this resolved if breakpoint regions of specific CRs shared common genomic signatures.

### DNA extraction, RAD-seq library preparation, data processing, and SNP filtering

Codominant markers from the parents, F_1_s and F_2_s involved in both crosses (CAD × ALF, CAD × HUE) were genotyped using RAD-seq. DNA was extracted with the *DNeasy Plant Mini Kit* (Qiagen, Valencia, CA, USA) following the manufacturer’s instructions, except that the incubation step at 65 °C was extended to two hours to increase DNA yield. RAD-seq libraries were prepared following Baird et al. (2008), using the restriction enzyme *PstI*, by Floragenex Inc. (Beaverton, OR, USA). Libraries were sequenced as single-end 150 bp reads on an Illumina Hiseq 2000 platform.

The quality of the RAD-seq data was first assessed with *FastQC* v0.12.1 (Andrews, 2010). After confirming that the data met per base and per sequence quality standards, reads from both crosses were clustered and analyzed using *Stacks* v2.65 (Rochette et al., 2019). Libraries were initially demultiplexed using *process_radtags*. The resulting reads were then aligned against the reference parental genomes with the *mem* algorithm of *BWA* v0.7.18 (Li and Durbin, 2009). Reference genomes were initially indexed using *faidx* in *SAMtools* v1.21 (Danecek et al., 2021). For CAD × ALF, in which both parental genomes were sequenced, reads were aligned to each parental genome. In contrast, CAD × HUE F_2_ reads were aligned only against the CAD parental genome. A catalog of SNPs was then built with *gstacks*, and for each of the three datasets the *populations* program in *Stacks* was run to generate SNP matrices. In this final step, only markers present in at least 90% of the F individuals were retained, specifying the F map type and output in VCF format.

The three VCF files were then filtered with *VCFtools* v0.1.17 (Danecek et al., 2011) and *BCFtools* v1.22 (Danecek et al., 2021). Only reciprocally homozygous biallelic loci present in both parents were retained, and loci located within repetitive regions were removed.

### Genotype validation, quality control and visualization

The pedigree of both experimental crosses was validated through genotyping, using polymorphic microsatellite markers originally developed for the closely related *C. helodes* (Arroyo et al., 2016) and successfully adopted for *C. laevigata*. Four loci showing polymorphism between the parents were selected (WE1, 468, FG7, and XAF), and the results were visualized using *Peak Scanner* v1.0.

To assess the reliability of the filtered RAD-seq SNP datasets and to detect potential genotyping inconsistencies, the three datasets were analyzed with the R package *diemR* (Baird et al., 2023). This package allows the detection of genotype polarization errors, the estimation of individual heterozygosity, and the visualization of genotype clustering across individuals. Filtered VCF files were analysed using the R package *vcfR* (Knaus & Grünwald, 2017). Polarized genotypes were then imported with *importPolarized*, and individual heterozygosity (HI) was computed as the proportion of heterozygous loci per individual using the function *pHetErrOnStateCount*. Visualization of the results was carried out through the *diemR* plotting functions *plotPolarized* and *plotDeFinetti*, implemented within the R package RColorBrewer (Neuwirth, 2022).

### Linkage mapping and recombination rate estimation

Linkage mapping and recombination analyses were performed with the *R* package *OneMap* (v3.2.2; Margarido et al., 2007). Chromosome files were generated for each cross and reference genome. Each chromosome file was imported with the function *read_mapmaker*. A segregation distortion test was applied to all markers using *test_segregation*, and loci showing significant deviation from Mendelian expectations (p < 0.05) were flagged. To reduce redundancy among tightly linked markers, SNPs located within 50 kb of each other were removed. Pairwise recombination fractions were estimated for each chromosome using *rf_2pts*, and linkage groups were defined with the *group* function using a LOD threshold of 3 and a maximum recombination fraction of 0.5. To improve the accuracy and biological coherence of the genetic maps, linkage groups corresponding to readjusted chromosomes were further evaluated using pairwise recombination fraction matrices. Chromosomes exhibiting unusually large genetic distances or heterogeneous linkage patterns were inspected for discontinuities in recombination fraction values, indicative of weak or absent linkage between marker blocks. When clear recombination fraction discontinuities were detected (i.e., tightly linked blocks of markers showing low recombination fractions with respect to adjacent regions in readjusted chromosomes), the corresponding chromosomes were subdivided into independent segregation units. These units represent analytical mapping entities rather than definitive physical chromosomes, and their delimitation aims to prevent artificial map expansion caused by the inclusion of poorly linked marker regions. Marker order within each linkage group followed the physical order of the reference genome assembly and was implemented using the *map* function in *OneMap*. Linkage maps were manually curated, removing outlier markers exhibiting strong segregation distortion or disproportionately large genetic distances that inflated overall map length to improve map stability and interpretability. The resulting linkage maps were visualized using LinkageMapView (Ouellette et al., 2018).

Recombination landscapes of all the chromosomes from the three datasets were visualized using *ggplot2*. Genome-wide recombination rates were estimated for each dataset by dividing the total genetic map length in centimorgan (cM) by the corresponding physical genome size (Mb). The same approach was applied at the chromosome level to obtain chromosome-specific recombination rates.

Chromosomes were classified as rearranged or non-rearranged based on the GENESPACE analyses. To determine whether CRs were associated with alterations in recombination landscapes, we evaluated recombination patterns at two complementary scales. First, we tested for localized shifts in recombination along chromosomes at the marker level. Second, we assessed whether rearranged chromosomes exhibited broader chromosome-wide changes in average recombination rates and variance structure relative to non-rearranged chromosomes. At the marker level, a Linear Mixed Model (LMM) was implemented using the formula *Rate ∼ Status + (1|Chromosome)* to test for localized differences while accounting for the nested data structure and intra-chromosomal autocorrelation. At the chromosomal scale, a Generalized Additive Model for Location, Scale and Shape (GAMLSS) was implemented. This framework allowed us to model both the mean (μ) and the dispersion (σ) parameters simultaneously as a function of chromosome status (rearranged vs. non-rearranged) while explicitly incorporating physical chromosome size (Mb) as a covariate in both formulas (μ *∼ Status* + *Size*; log(σ) *∼ Status* + *Size*). This unifies the control for the biological constraints of chromosome size on mean recombination rates and directly tests for chromosome-wide desynchronization or inflation of variance associated CRs. All models were implemented in R using the packages lme4 v2.0 (Bates et al., 2015) and lmerTest v3.2 (Kuznetsova et al., 2017) for LMMs, and gamlss v5.9 (Rigby and Stasinopoulos, 2005) for chromosomal-level modeling. Residual diagnostics for GAMLSS were verified using randomized quantile residuals to ensure normal behavior and model assumptions were met.

To assess whether CRs were associated with segregation distortion, markers were classified into three distortion categories based on their segregation test *p*-values: non-distorted (*p* ≥ 0.05), moderately distorted (0.001 ≤ *p* < 0.05), and severely distorted (*p* < 0.001). For each dataset, the proportion of markers falling into each distortion category was calculated separately for rearranged and non-rearranged chromosomes. These proportions were then compared to evaluate whether rearranged chromosomes exhibited an enrichment of segregation distortion relative to structurally collinear chromosomes.

### Quantitative Trait Loci (QTLs) Analysis

A total of 15 morphological traits were measured for 155 F_2_ individuals for the QTL analysis. These included reproductive traits— number of male spikes, number of female spikes, number of androgynous spikes, total dry weight of the female portion of all spikes, total dry weight of the male portion of all spikes, number of mature utricles, total dry weight of mature utricles, number of floral stems, total floral stem weight excluding spikes, length of the longest floral stem and total reproductive biomass—and dry biomass and vegetative traits, including aboveground biomass excluding stems, maximum leaf length, maximum leaf width, and rhizome weight. Length measurements were recorded using a ruler to the nearest mm, while weight measurements were obtained using a laboratory balance to the nearest mg (Ohaus Adventurer AR3130). To explore the relationships between the 15 measured morphological traits, a pairwise correlation analysis was performed using Spearman’s rank correlation coefficient (*r*) with the R package *corrplot* (Wei and Simko, 2024).

QTL analyses were performed using the *R* package *R/qtl* (Browman et al., 2003). Quantitative traits related to plant vigor (e.g., leaf and rhizome weight) were log-transformed log_10_(x+1) to meet normality assumptions. The number of androgynous spikes and utricles were converted into binary variables (presence/absence) and were analyzed with the binary model. Reproductive traits characterized by a high frequency of zeros (zero-inflation) were analyzed using a separate two-part model.

Genotype probabilities were calculated at 1 cM intervals with an assumed genotyping error rate of 0.01 using the *calc.genoprob* function. Main QTL effects were detected via a single-QTL genome scan (*scanone*). The Expectation-Maximization (EM) algorithm was applied for binary and two-part models, while Haley-Knott (HK) regressions were utilized for normal models to optimize computational efficiency.

Significance thresholds for Logarithm of Odds (LOD) scores were determined through 1,000 genome-wide permutations to maintain a family-wise error rate of α = 0.05. For significant QTLs, the 95% Bayesian credible intervals were calculated using the *bayesint* function to define the most probable genomic location. Gene content within these intervals was subsequently examined using genome annotations generated with BRAKER. The total percentage of phenotypic variance explained (*R^2^*) by each identified QTL was estimated by fitting a multiple QTL model using the *fitqtl* function. To ensure accuracy in variance estimation, the imputation method was employed via the *sim.geno* function with 64 draws. Finally, the phenotypic effects of the identified loci were visualized by plotting phenotype distributions across the underlying genotypic classes—AA (homozygous for the maternal genome), AB (heterozygous), and BB (homozygous for the paternal genome)—at the lead marker of each significant peak.

## Results

### Experimental crosses, germination success and cytogenetics

The experimental crosses revealed a significant decline in germination success as breeding progressed from parental lines to F hybrids. A total of 83 crosses were performed between 2019 and 2023: 17 selfed parent crosses, 9 hybrid crosses between different populations to generate F s, and 57 selfing of F hybrids to generate F s, producing 1,265, 393 and 4,174 achenes, respectively. Of these, 573, 154, and 567 seeds germinated for the selfed parents, hybrid crosses, and selfed F_1_s crosses, respectively. The mean germination success were 0.519 for selfed parent crosses (0.476 for ALF, 0.708 for CAD, and 0.310 for HUE), 0.398 for hybrid crosses (0.243 for CAD × ALF and 0.593 for CAD × HUE), and 0.173 for selfed F_1_s individuals (0.116 for CAD × ALF and 0.192 for CAD × HUE) (Supplementary Fig. 1; Supplementary Table1: Fig. 1c). Subsequently 78 F s developed individuals from the CAD × ALF cross and 148 F s from the CAD × HUE cross were studied for QTLs and genotyped with RADseq.

Cytogenetic study confirmed that while parents maintained stable bivalent formations, F and F hybrids exhibited complex and more variable meiotic configurations consistent with intermediate chromosome numbers. As expected, karyotyping revealed chromosome numbers of 2n = 76 for the HUE parent, and 2n = 80 for the CAD parent, all of which displayed exclusively bivalent formations. Surprisingly, the ALF parent exhibited two distinct karyotypes, 2n = 74 and 2n = 76 (Fig. 1b), with 2n = 74 being much more frequent. The studied F s from the CAD × ALF cross exhibited 34 bivalents, one tetravalent, one trivalent, and two univalents, resulting in 2n = 34^II^+1^IV^+1^III^+2^I^= 77, consistent with an intermediate karyotype between the parental chromosome numbers, 2n = 74 and 2n = 80 (Fig. 1b). In contrast, the F generation showed a different configuration, consisting of 36 bivalents and one tetravalent (2n = 36^II^ + 1^IV^ = 76). The F s from the CAD × HUE cross typically showed 36 bivalents and two trivalents, yielding 2n = 36^II^+2^III^= 78 also consistent with an intermediate karyotype between the parental chromosome numbers, 2n = 76 and 2n = 80. However, the F progeny displayed a distinct configuration, with 36 bivalents, one tetravalent, one trivalent, and one univalent (2n = 36^II^ + 1^IV^ + 1^III^ + 1^I^ = 80).

### Genome assemblies and annotations

We produced three chromosome-scale, annotated genomes from our sampled *C. laevigata* populations in the Iberian Peninsula. The median PacBio read length was 12,187 bp for the ALF assembly, 12,437 bp for the CAD genome, and 12,739 bp for the CAD × HUE F genome. Mean read quality scores were 33.4 (ALF), 33.8 (CAD), and 37.4 (CAD × HUE F), with estimated sequencing coverages of 12.9× and 40.4×, respectively.

The final assemblies consisted of 38 and 40 scaffolds for the ALF and CAD genomes, respectively, and 37 and 36 scaffolds for the CAD × HUE F haplotype 1 and 2, respectively (Supplementary Fig. 2). Notably, the number of scaffolds in the ALF assembly did not match the most common karyotyped chromosome number (2n=37^II^=74) but matched the alternative karyotype found in the germ line (2n=38^II^=76; Fig. 1b). The total assembly lengths were 377.47 Mb for ALF, 385.23 Mb for CAD, 384.13 Mb for CAD × HUE F haplotype 1, and 383.51 Mb for CAD × HUE F haplotype 2. Scaffold continuity resulted in scaffold N50 values of 10.5 Mb for ALF, 9.40 Mb for CAD, 10.67 Mb for the CAD × HUE F haplotype 1 and 10.65 Mb for CAD × HUE F haplotype 2. Although the CAD × HUE F assembly was generated in a phased mode, it should not be interpreted as a complete or fully accurate reconstruction of the HUE and CAD parental genomes. Due to selfing (Hipp et al., 2009; Escudero et al., 2013a), genomes in *C. laevigata* are highly homozygous, limiting the ability to distinguish haplotypes and increasing the likelihood of allelic collapse. Consequently, the structure of the F assembly may deviate from the expected parental chromosome number, including the absence of the expected n=38 karyotype, and likely reflects a combination of true haplotypic signal and assembly artefacts rather than a fully resolved representation of segregating HUE and CAD parental chromosomes.

BUSCO indicated high completeness across all assemblies, recovering 95.22% complete single-copy genes in ALF (2.13% duplicated, 0.84% fragmented, 1.8% missing) and 96.05% in CAD (2.18% duplicated, 0.81% fragmented, 0.96% missing). For the CAD × HUE F hybrid, haplotype 1 recovered 96.05% complete genes (2.15% duplicated, 0.83% fragmented, 0.97% missing), and haplotype 2 showed comparable results with 96.04% completeness (2.16% duplicated, 0.80% fragmented, 1.00% missing). Repetitive elements were the predominant genomic feature across all assemblies, accounting for 59.3% (29.2% unclassified) in ALF, 60.3% (24.9%) in CAD, 59.6% (25.8%) in EDI, and 60.4% (25.1%) and 58.67% (26.67%) in CAD × HUE F haplotypes 1 and 2, respectively. Among the annotated repeats, long terminal repeat (LTR) elements were the most abundant class, comprising 14.6%, 15.9%, 14.3%, 17.9%, and 15.6% of the ALF, CAD, EDI, and the two F haplotypes, respectively. Gene prediction using the BRAKER3 pipeline identified 26,941, 27,121, 29,085, 28,289 and 29,601 protein-coding genes in the ALF, CAD, EDI, and CAD × HUE F haplotypes 1 and 2, respectively.

### Synteny analysis

The synteny analysis revealed substantial chromosomal restructuring among the analyzed *C. laevigata* genomes (Fig. 1a). Although broad-scale synteny was conserved across most chromosomes, multiple types of CRs, including chromosome fusions, fissions, translocations, and inversions were detected. The EDI genome represents the lineage with the lowest haploid chromosome number (n = 36), and exhibits most chromosomal fusions, alongside translocations and inversions relative to the other assemblies. Comparisons between the ALF and CAD genomes revealed several structural differences, including a fusion/fission involving ALF chromosome 4 and CAD chromosomes 19 and 39 (Fig. 1a), a reciprocal translocation with inverted segments between ALF chromosomes 22 and 32 and CAD chromosomes 22 and 33, and a complex rearrangement involving ALF chromosomes 15 and 16 and CAD chromosomes 24, 29, and 40 (Fig. 1a). CAD × HUE F haplotypes recapitulate the same rearranged regions observed in the CAD and ALF genomes, with highly disrupted synteny between haplotypes (Supplementary Fig. 3).

### Genomic features and enrichment analysis

The mean proportion of coding sequences (CDS) in each genome was 11.89% in EDI, 7.96% in ALF, and 7.81% in CAD. GC content was broadly similar across genomes, with mean values of 33.93% (EDI), 34% (ALF), and 34.03% (CAD). Overall, coding regions and repeat elements were comparatively evenly distributed along the genomes (Supplementary Fig. 4 to 6); however, as expected, repeat elements and GC proportion showed a pronounced enrichment toward telomeric regions in all three assemblies. Genomic landscape analysis revealed that GC content is positively associated with both coding and repetitive fractions. Notably, CDS and repeat proportions showed a significant mutual exclusion, evidenced by their strong inverse correlation in the three analyzed populations (Supplementary Fig. 7).

Breakpoint regions exhibited consistent, assembly-specific associations with underlying genomic features, most notably characterized by CDS depletion and GC enrichment across all genomes, regardless of whether chromosomal ends were included or excluded from the analysis (Fig. 2; Supplementary Table 2). Among repeat families, LTR-gypsy elements were the only repeats consistently associated with breakpoint regions across all genomes when chromosomal ends were excluded. However, this association was significant only in the CAD genome and marginally (*p* < 0.1) significant in the EDI and ALF genomes (Fig. 2).

**Figure 2.**
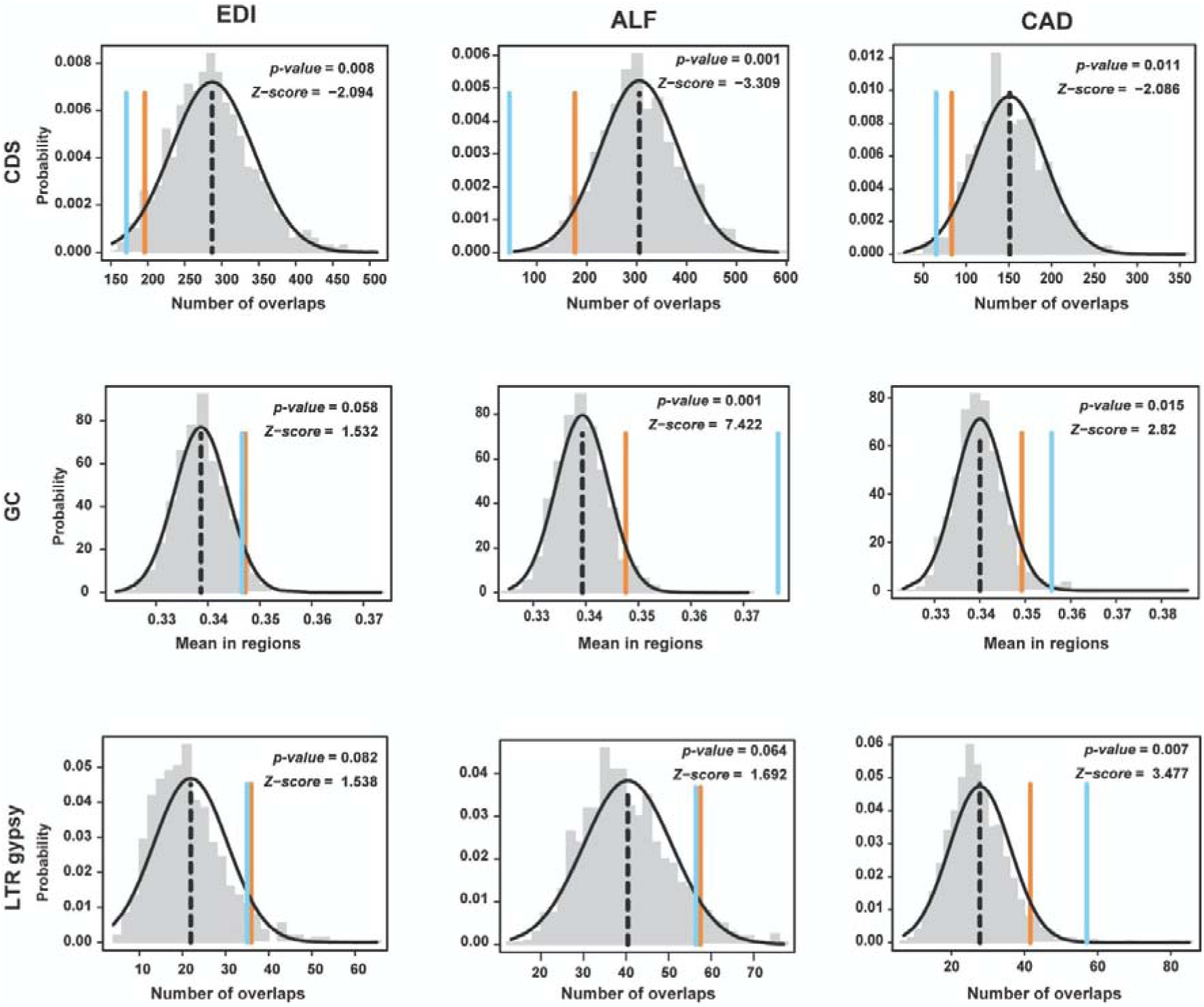
Enrichment of genomic features in breakpoint regions. Panels show the enrichment for each genome (columns) and each genomic feature (rows), including coding sequences (CDS), GC content, and LTR/Gypsy elements. In each panel, the black dashed line represents the expected mean overlap under the null model, the orange line marks the 95% expected interval, and the blue line indicates the observed overlap. Empirical *p*-values and *z*-scores are shown, with positive *z*-scores indicating over-representation and negative values indicating depletion within breakpoint regions. Chromosome ends were excluded from the enrichment analysis to prevent telomeric bias.

Analyses of individual breakpoint regions associated with CRs in each assembly partially confirmed the patterns described above (Supplementary Tables 3-5). One notable exception was CDS content, which generally showed only marginally significant depletion across breakpoint regions (Supplementary Tables 3-5). GC proportion exhibited significant enrichment in many breakpoint regions; however, patterns were variable, as numerous breakpoints showed no significant enrichment or instead displayed GC depletion (Supplementary Tables 3-5). This analysis also uncovered highly specific associations with repeat families that were not apparent in the genome-wide analyses. Overall, breakpoint regions in the EDI genome were predominantly enriched for DNA transposon families, LTR Gypsy elements, and low-complexity regions (Supplementary Table 3). In the ALF genome, breakpoint enrichment was also observed for LTR-Gypsy elements as well as for several DNA transposon families, including EnSpm, TcMar, hAT-Ac, and hAT-Tip100, as well as satellite and simple repeats (Supplementary Table 4). In the CAD genome, breakpoint enrichment was associated with both DNA transposons and retrotransposons. Enriched DNA transposon families included PIF-Harbinger and hAT-Ac, whereas enriched retrotransposons comprised LINE-L1, LINE-RTE-BovB, LTR-ERV1, LTR-Copia, and LTR-Gypsy elements (Supplementary Table 5). Beyond genome-specific patterns, we identified homologous breakpoint regions across assemblies that shared common genomic signatures, characterized by an enrichment for the same repeat families. For instance, the breakpoint on chromosome 1 of the EDI genome is significantly enriched for TcMar and hAT-Tip100 transposable elements (Supplementary Table 3). Notably, one of the fissioned counterparts (chromosome 8) in the ALF assembly exhibits enrichment for the same repeat families (Supplementary Table 4). Similarly, low-complexity regions are significantly enriched at a breakpoint on chromosome 6 in the EDI genome (Supplementary Table 3) and show marginal enrichment at the corresponding breakpoint on fissioned chromosome 27 in the ALF genome (Supplementary Table 4). Finally, LTR-Gypsy elements are significantly enriched in a breakpoint region on chromosome 16 of the ALF genome (Supplementary Table 4) as well as in the corresponding breakpoint region on chromosome 29 of the CAD genome (Supplementary Table 5).

### Genotype validation & RAD-seq data processing

Microsatellites confirmed that the F individuals from both crosses matched the genotypes of their respective parents (Supplementary Table 6). Among the cultivated F progeny, 55 individuals from the CAD × ALF cross and 144 individuals from the CAD × HUE cross showed microsatellite profiles consistent with their parental genotypes and were therefore selected for RAD-seq (Supplementary Table 6), generating a total of 2,171,612,556 reads. Of these, 15,931,539 reads were removed due to ambiguous RAD tags, 445,027,601 due to barcode mismatches, and 1,403,076 due to low quality. In total, 1,723,119,915 reads (79.3% of the raw dataset) were retained for downstream analyses, with per-individual coverage ranging from 2,776,724 to 18,379,774 reads.

For the CAD × ALF cross, *gstacks* identified 555,390 loci in the catalog. The *populations* module retained 43,468 and 43,767 loci when aligned to the ALF and CAD reference genome, respectively. After applying the final filtering criteria, the resulting VCF files contained 13,922 loci (ALF as reference) and 13,932 loci (CAD as reference). For the CAD × HUE cross, *gstacks* assembled a catalog of 1,884,372 loci. Of these, the *populations* module retained 97,374 loci, and the final filtered VCF comprised 12,287 loci.

The *diemR* analysis revealed several individuals whose genotype profiles were inconsistent with the expectations for true F progeny. In the CAD × ALF cross, one putative F was identified as a backcross between the F and the CAD parent, and three additional individuals displayed biologically implausible patterns of heterozygosity (Supplementary Fig. 8). Specifically, in the *diemR* polarity plots, closely linked markers occasionally exhibited opposite polarities, an unlikely scenario given their physical proximity, whereas in genuine F individuals polarity switches typically occur only across larger genomic regions. These four individuals were therefore excluded from all downstream analyses, resulting in a dataset of 51 F individuals. In the CAD × HUE cross, fifteen individuals were either identified as backcrosses between an F and a parent or exhibited similarly improbable polarity patterns inconsistent with F Mendelian expectations (Supplementary Fig. 8). These individuals were likewise removed to avoid distortions in subsequent linkage and recombination analyses, resulting in a dataset of 116 F individuals.

### Linkage mapping and segregation distortion

Subdivision of linkage groups across all mapping populations effectively mitigated genetic distance inflation caused by localized clusters of segregation distortion. For the CAD × ALF cross, F individuals were aligned to both parental reference genomes (CAD and ALF). After filtering for outliers and retaining markers spaced at least 50 kb apart, ∼3,400 high-quality SNPs were used for linkage map construction (Supplementary Table 9). Initial maps produced one linkage group per chromosome, but several chromosomes showed inflated genetic distances associated with clusters of segregation-distorted markers. Detailed inspection of recombination fraction matrices (Supplementary Figs. 9-11), synteny analyses (Fig. 1a), and segregation distortion profiles revealed the presence of distinct segregation zones, requiring manual subdivision of affected linkage groups. In the CAD-based map, chromosome 29 was split into four linkage groups, while chromosomes 22 and 33 were each split into two. Similarly, in the ALF-based map, chromosomes 15, 22, and 32 required subdivision. These adjustments substantially reduced map inflation and improved marker ordering, as reflected by decreased total and mean linkage group lengths (Supplementary Table 8; Supplementary Figs. 12 and 13). A comparable pattern was observed in the CAD × HUE cross, where markers were aligned to the CAD reference genome. Following filtering, 3,200 markers were retained (Supplementary Table 7). As in the CAD × ALF maps, chromosomes exhibiting strong segregation distortion, most notably chromosome 29, as well as chromosomes 22 and 33, required subdivision into multiple linkage groups (Supplementary Table 4; Supplementary Fig. 14). Splitting these chromosomes markedly reduced inflated genetic distances and improved overall map quality. Markers exhibiting segregation distortion were more abundant in chromosomes with CRs in all datasets (Fig. 3a).

**Figure 3.**
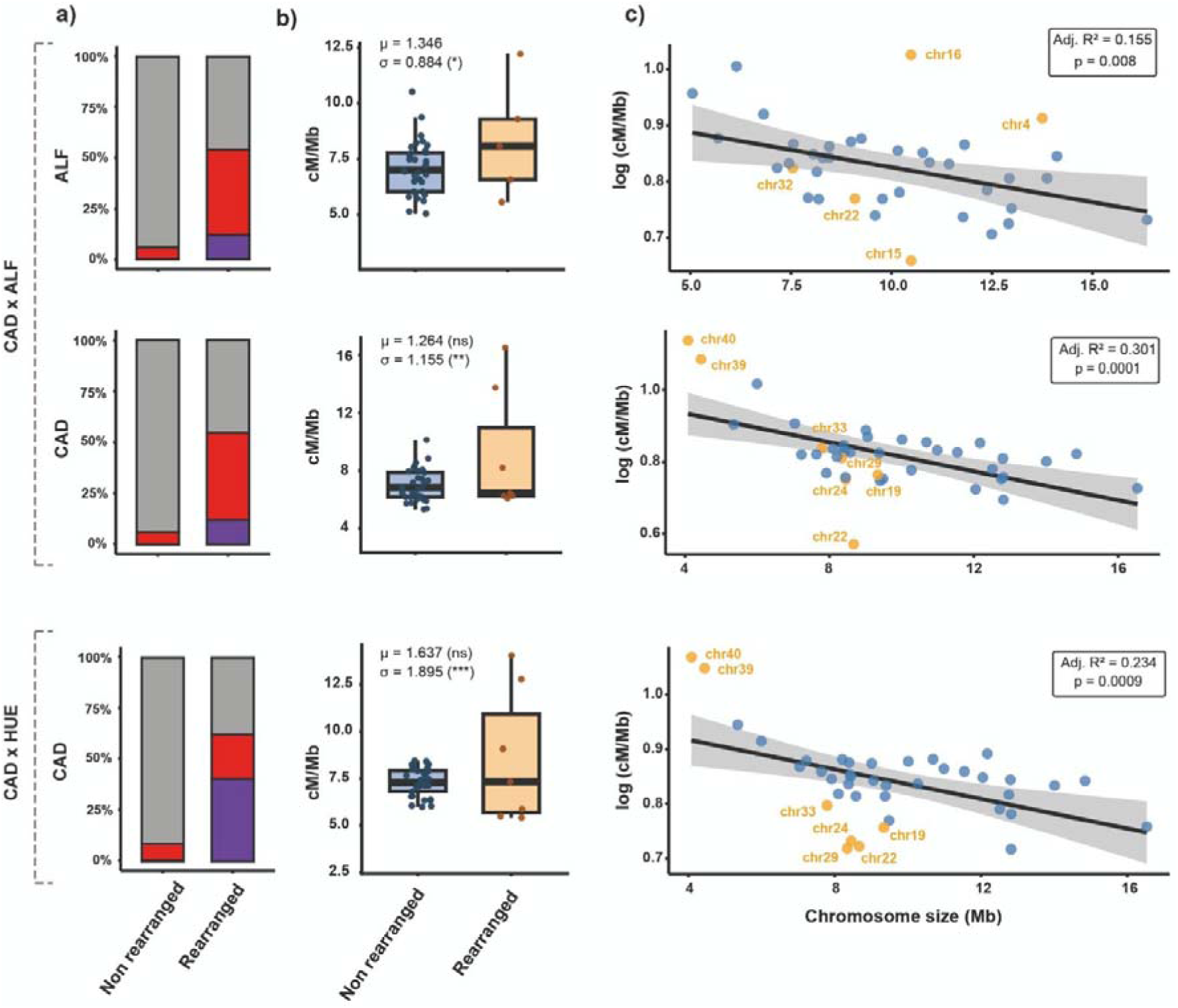
Segregation distortion and recombination in rearranged and non-rearranged chromosomes. **a)** Proportion of markers showing segregation distortion in rearranged versus non-rearranged chromosomes: Severely distorted markers (purple, *p* < 0.001), moderately distorted markers (red, 0.001 < *p* ≤ 0.05), and grey markers without significant distortion. **b)** Boxplots showing chromosome-wide recombination rates (cM/Mb) for rearranged and non-rearranged chromosomes. Dots represent individual chromosomes colored according to their respective category. Inserted text indicates the GAMLSS effect sizes and significance levels for the mean (μ) and dispersion/variance (σ) parameters after controlling for chromosome size. **c)** Relationship between chromosome size and mean recombination rate. Orange points represent chromosomes with CRs. The line shows the fitted linear model with the 95% CI highlighted in grey. Statistical parameters of the fitted linear models are specified in Supplementary Table 4. Significance levels: *p* < 0.05 (*), *p* < 0.01 (**), *p* < 0.001 (***), *ns* = non-significant.

### Recombination across genomes and CRs

Mean recombination rates were broadly similar among crosses and reference genomes, ranging from 6.48 to 6.96 cM/Mb on average, although substantial inter-chromosomal variation was observed in all cases (Supplementary Table 9). In both the CAD × ALF and CAD × HUE crosses, rearranged chromosomes consistently exhibited higher recombination rates than non-rearranged chromosomes (Fig. 3b; Supplementary table 10). Marker-level analyses using LMMs further supported this trend, revealing significantly elevated local recombination rates in rearranged chromosomes in both parental maps of the CAD × ALF cross (ALF: β = 1.485 ± 0.643, *p* = 0.027; CAD: β = 1.391 ± 0.673, *p* = 0.047), whereas the same tendency in the CAD × HUE cross was not statistically significant (*p* = 0.236) (Supplementary Table 10). At the chromosomal scale, GAMLSS analyses indicated that rearranged chromosomes did not exhibit significant shifts in mean recombination rates after accounting for chromosome size (μ parameter; all *p* > 0.2). In contrast, rearranged chromosomes consistently showed significantly higher recombination variance (σ parameter) across all datasets (ALF: σ = 0.884 ± 0.341, *p* = 0.014; CAD: σ = 1.155 ± 0.362, *p* = 0.003; CAD × HUE: μ = 1.895 ± 0.383, *p* < 0.001), indicating increased heterogeneity in recombination landscapes associated with CRs (Supplementary Table 10).

Across all three datasets, chromosome size was significantly and negatively associated with mean recombination rate, with smaller chromosomes exhibiting higher recombination rates (Fig. 3c). The exclusion of rearranged chromosomes from the datasets improved model fits in all cases except for the CAD genome in the CAD × ALF cross (Supplementary Figs. 15 to 17).

Recombination landscapes of non-rearranged chromosomes were characterized by moderate variation in recombination rate along the chromosome without abrupt transitions or extended regions of suppressed recombination (Supplementary Figs. 9-11). Larger chromosomes generally showed increased recombination rates toward subtelomeric regions, while smaller ones showed a more homogeneous pattern. Regions showing segregation distortion were typically restricted to telomeric regions, as illustrated by chromosomes 4, 14, and 21 of the CAD genome in the CAD × HUE cross (Supplementary Fig. 11). Segregation distortion was occasionally detected in rearranged chromosomes relative to the EDI assembly, such as chromosomes 1 and 8 in the ALF genome and chromosomes 1 and 5 in the CAD genome in the CAD × ALF cross (Supplementary Figs. 9 and 10).

Rearranged chromosomes exhibited recombination landscapes that differed markedly from those of non-rearranged chromosomes, displaying highly heterogeneous recombination patterns, characterized by sharp transitions in recombination rate, with localized peaks interspersed by extended regions of suppressed recombination. In the CAD × ALF cross, the chromosomal fusion/fission (Fig. 4a) showed pronounced suppression of recombination not only in the vicinity of the breakpoint region on chromosome 4 of the ALF genome, but also across a large central region of this chromosome and in the syntenic regions of chromosomes 19 and 39 of the CAD genome. Elevated recombination rates were observed in subtelomeric regions of these chromosomes. The reciprocal translocation (Fig. 4b), exhibited reduced recombination relative to subtelomeric regions and the translocated segments especially for interstitial segments of chromosome 22 in the ALF genome and chromosome 33 in the CAD genome. This rearrangement was also associated with pronounced segregation distortion in breakpoint-proximal regions. In the complex rearrangement (Fig. 4c), recombination was especially suppressed near breakpoint regions across all chromosomes involved. Notably, inverted regions on chromosomes 15 and 16 of the ALF genome, as well as their corresponding regions in the CAD genome, exhibited near-complete suppression of recombination. As in other rearrangements, recombination was largely confined to subtelomeric regions. Strong segregation distortion was again detected near breakpoint regions and was particularly pronounced on chromosome 15 of the ALF genome and chromosome 24 of the CAD genome, where nearly half of the markers displayed severe segregation distortion.

**Figure 4.**
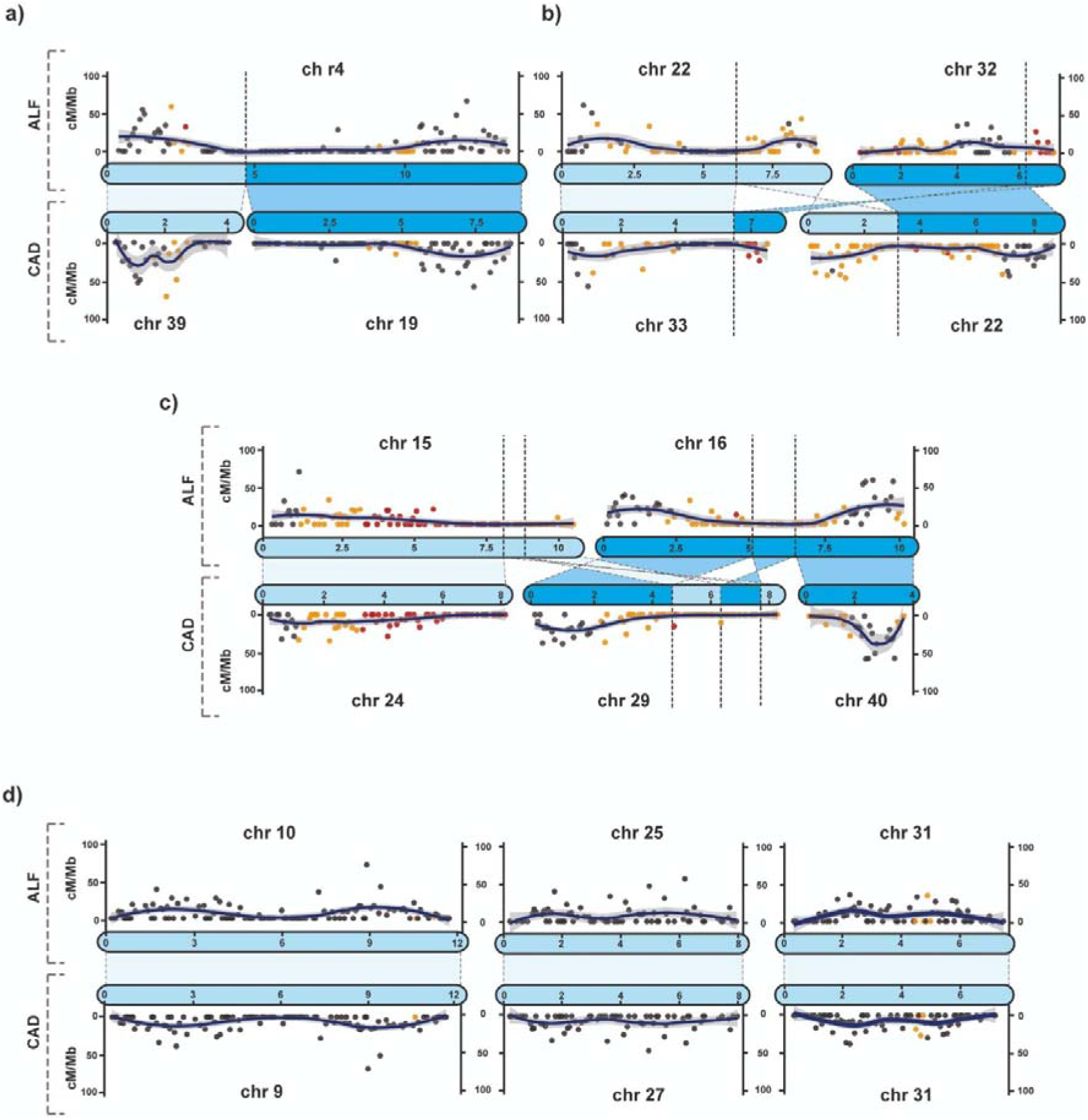
Recombination landscapes in rearranged and non-rearranged chromosomes in the CAD × ALF cross. Panels **a–d** show recombination landscapes for chromosomes involved (**a-c**) and not involved (**d**) in different CRs between the ALF (top) and CAD (bottom) genomes. In each panel, the coloured schematic blocks illustrate the orientation and relative positions of chromosomal segments in each genome, where dark and light blue indicate rearranged and non-rearranged segments, respectively, and dashed lines connecting chromosomes delimitate homologous regions. Scatter points represent local recombination rate estimates (cM/Mb), and solid lines show the fitted LOESS curves. Vertical dashed lines mark the inferred breakpoints of each rearrangement. Red dots indicate severely distorted markers (*p* < 0.001), orange dots moderately distorted markers (0.001 < *p* ≤ 0.05), and grey dots non-distorted markers.

With respect to the CAD × HUE cross, rearranged chromosomes in the CAD genome displayed recombination patterns similar to those observed in the CAD × ALF cross, but with markedly stronger segregation distortion (Supplementary Fig. 10). Most rearranged chromosomes (chromosomes 19, 22, 24, 29, 33, 39, and 40) contained a high proportion of severely distorted markers (Supplementary Fig. 10). As in the CAD × ALF cross, extended regions of suppressed recombination were detected in these chromosomes, with recombination largely confined to telomeric and subtelomeric regions.

### QTL analysis

The F_2_ cohort exhibited substantial variation across all 15 measured traits. Descriptive statistics revealed that while vegetative vigor remained relatively stable, reproductive effort was highly variable, with higher coefficients of variation (Supplementary Table 11). Reproductive, vegetative, and biomass-related traits showed strong self-correlation within each trait category (Supplementary Fig. 18). However, correlations between reproductive and biomass related traits were notably lower.

Genome-wide scans identified three major regions associated with reproductive success: on chromosome 16 of the ALF genome and chromosome 40 of the CAD genome in the CAD × ALF cross, and on chromosome 4 of the CAD genome in the CAD × HUE cross. In the CAD × ALF cross, these regions were consistently associated with both the number of floral stems and the number of male spikes (Fig. 5; Supplementary Figs. 19 and 20; Supplementary Table 12), whereas in the CAD × HUE cross the signal was linked to floral stem weight and total reproductive biomass (Fig. 5; Supplementary Fig. 21; Supplementary Table 12). Notably, the two regions detected in the CAD × ALF cross are syntenic and map to the same large-scale chromosomal region near the breakpoint of a major inversion (Fig. 4c). In this cross, a QTL peak in the ALF genome was identified at 63.35 cM (8,354,929 bp), with a Logarithm of the Odds (LOD) score of ∼8.8 exceeding permutation-based thresholds (7.88–8.01) and a ∼20 cM 95% Bayesian credible interval (60–82/83 cM); individuals carrying the BB genotype (homozygous for CAD alleles) exhibited higher reproductive output, indicating a dominant effect and explaining ∼12.3% of variance (Supplementary Table 12). This interval contained 65 annotated genes. (Fig. 5a). A second QTL in the CAD genome (chromosome 40) was detected at 8.24 cM (1,925,252 bp), showing similar statistical support (LOD ∼8.8; thresholds 8.03–8.09; ∼20 cM interval at 5–27/28 cM) but a contrasting additive genetic architecture, with reproductive output increasing with the number of ALF alleles (Fig. 55b), and accounting for a comparable proportion of phenotypic variance (∼12.3%; Supplementary Table 12). The corresponding Bayesian interval included 59 annotated genes (Fig. 5a). In the CAD × HUE cross, the detected region on chromosome 4 of the CAD genome is not associated with a chromosomal rearrangement and was identified at 54 cM (9,749,532 bp), with a peak LOD score of 8.98 exceeding permutation-based thresholds (8.00–8.05) for both floral stem weight and total reproductive biomass; the 95% Bayesian credible interval was narrow (54–54 cM), and the locus explained approximately 8.2–8.24% of the phenotypic variance (Fig. 5; Supplementary Fig. 21; Supplementary Table 12). Individuals with homozygous genotypes (AA or BB) exhibited higher reproductive output than heterozygotes (AB), with the strongest effect observed in BB individuals (homozygous for the HUE parent; Fig. 5b). This interval contained only five annotated genes (Fig. 5a).

**Figure 5.**
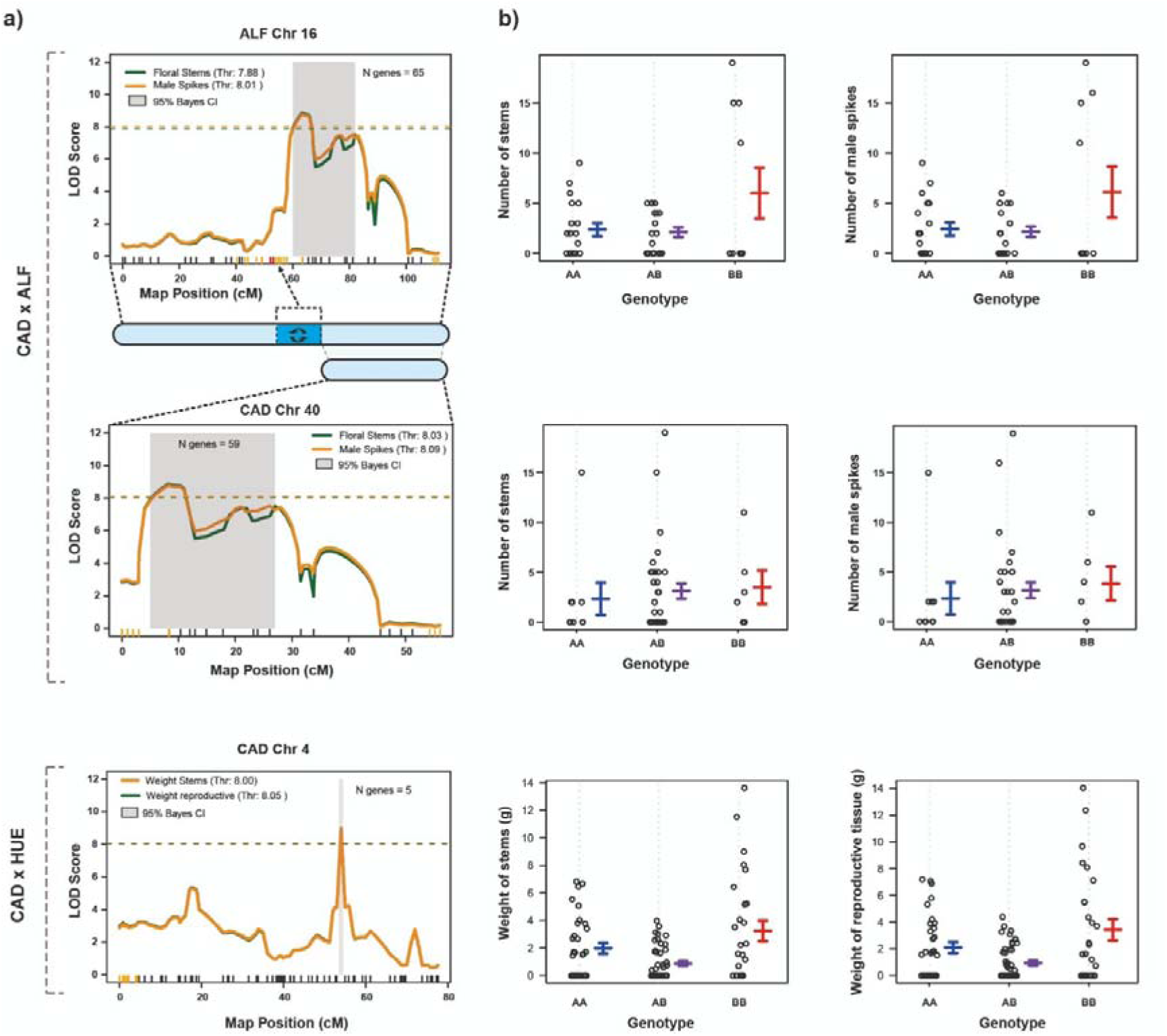
QTL mapping and phenotypic effects for reproductive effort. **a)** For the CAD × ALF cross, Logarithm of the Odds (LOD) score profiles for number of floral stems (green) and number of male spikes (orange) on ALF chromosome 16 and CAD chromosome 40. For the CAD × HUE, LOD score profiles for weight of floral stems (orange) and weight of reproductive biomass (green) on CAD chromosome 4. Horizontal dashed lines indicate significance thresholds (α = 0.05) based on 1,000 permutations. Shaded gray areas represent the 95% Bayesian credible intervals for the identified QTL. Ticks along the x-axis correspond to the genetic markers used to construct the linkage groups; orange and red ticks indicate markers with moderate and severe segregation distortion, respectively, whereas black ticks indicate no segregation distortion. The dark blue region in the schematic representation of CAD chromosome 16 corresponds to the chromosomal inversion. **b)** Genotype effect plots at the peak markers for each chromosome. Points represent individual observations, while colored bars indicate the mean standard error for the three genotypes: homozygous for the maternal genome (AA), heterozygous (AB), and homozygous for the paternal genome (BB). N genes indicate the number of genes found in the calculated Bayesian interval.

## Discussion

### CRs in *Carex laevigata* and their genomic characterization

*Carex laevigata* exhibits remarkable variation in chromosome number across its distribution (2n = 69-84; Luceño & Castroviejo, 1991; Escudero et al., 2013b; Fig. 1b). Our genomic analyses revealed a greater extent of structural variability than previously anticipated, as many CRs remained undetected in previous cytogenetic studies (Luceño and Castroviejo, 1991; Escudero et al., 2013b). Notably, the number of scaffolds in our ALF assembly (n = 38) does not match the most common karyotype in the germ line (2n = 37^II^). This discrepancy likely reflects intra-individual chromosomal variation, as different chromosome numbers have been reported within the germ line of single individuals of three different *Carex* species (Schmid, 1982; Luceño, 1992; Escudero et al., 2013a). It is therefore possible that the genome assembly, derived from vegetative tissue, displays the infrequent karyotype in the germ line with higher chromosome number.

Breakpoint regions associated with CRs were depleted for CDS across most genomes (Fig. 2; Supplementary Tables 2-5). This is expected, as breaks occurring within CDS are likely to disrupt gene function and reduce individual fitness, making them less likely to persist or become fixed in populations. In addition, although GC content appears broadly homogeneous across each genome (Supplementary Fig. 4-6), it is significantly elevated in most breakpoint regions (Fig. 2; Supplementary Tables 2 to 5). GC-rich regions are known to promote mutational events and increase the likelihood of double strand breaks (DSBs) (Fullerton et al., 2001; Kiktev et al., 2018). Because recombination can itself generate CRs (Farré et al., 2015), GC-enriched regions may therefore be more prone to structural instability. This intrinsic susceptibility is further amplified by the enrichment of specific transposon families at breakpoint regions (Supplementary Tables 2-5), especially LTR-Gypsy retrotransposons (Fig. 2). These elements can promote homology-driven ectopic recombination (Oliver et al., 2013), and facilitate mispairing during meiosis or DNA repair (Delprat et al., 2009). The combination of GC and transposable element enrichment, provide homologous substrates for ectopic recombination, likely making these genomic regions prone to breakpoint formation and CRs. This is further supported by our fine-scale analysis of individual breakpoints, uncovering specific associations with some repeat families (Supplementary Tables 3-5), providing evidence for the existence of recurrent CR hotspots. Likewise, the conserved enrichment of specific repeat elements at homologous breakpoints suggests that these regions are not only prone to breakage but also exhibit a persistent architectural vulnerability maintained over evolutionary time.

Overall, these results indicate that certain genomic regions are more susceptible to CRs in *C. laevigata*. Our synteny analyses reveal that some chromosomes (e.g. 8 and 27 in the EDI assembly), have undergone multiple independent rearrangements, forming dense clusters of breakpoints and repeated structural transitions (Fig. 1a; Supplementary Fig. 3). Consistent with hypothesis and findings that genomic hotspots of CRs are often associated with breakage-facilitating features like GC-rich windows and repetitive elements (Delprat et al., 2009; Farré et al., 2015; Kiktev et al., 2018; Escudero et al., 2023), these structural regions likely contribute disproportionately to karyotype diversification within *C. laevigata*, thereby explaining the remarkable variability observed across populations (Fig. 1a-b).

### Hybrid Dysfunction and Meiotic Barriers Associated with Chromosomal Variation

Hybrid dysfunction has long been recognized as a major driver of evolution and speciation (Coyne and Orr, 2004; Faria and Navarro, 2010). Karyotypic characterization of the *C. laevigata* crosses revealed that differences in CRs between parental lines with different chromosome numbers lead to abnormal configurations during Metaphase I of meiosis in F individuals, including univalents, trivalents, and tetravalents (Fig. 1b). These aberrant pairings can cause missegregation, reducing gamete viability or rendering some gametes nonfunctional, even in holocentric species (e.g., Lukhatanov et al., 2005; Escudero et al., 2016b). Indeed, mean germination success was significantly higher in selfed parental lines than in the F_1_, and, especially, the F progeny (Fig. 1c; Supplementary Fig. 1), suggesting that chromosomal mismatches in hybrids compromise gamete quality and seed viability. Interestingly, germination success in F_1_ and F_2_ was lower for crosses involving higher karyotypic differences (2n=80 × 74 vs 2n=80 × 76; Supplementary Fig. 1; Supplementary Table 1), which suggests that this effect could be stronger with increasing chromosomal divergence (Escudero et al., 2016).

Genomic markers showing segregation distortion were more abundant on rearranged chromosomes (Fig. 3a; Supplementary Figs. 9-11), suggesting that structural rearrangements, particularly those that disrupt proper pairing and segregation, generate systematic deviations from Mendelian expectations, likely by reducing the viability of some meiotic products. In the CAD × ALF cross, the fusion/fission rearrangement showed the lowest level of segregation distortion (Fig. 4a), consistent with the formation of a properly segregating trivalent configuration in heterozygotes (Nokkala et al., 2006; Lukhtanov et al., 2018), supported by recombination hotspots detected in the corresponding subtelomeric regions. This trivalent configuration would facilitate proper orientation and balanced segregation of the fused and fissioned chromosomes during meiosis, thereby reducing the production of unbalanced gametes and minimizing segregation distortion. The two other CRs betweenALF and CAD (Fig. 4b,c), i.e. a reciprocal translocation and a complex rearrangement, showed substantially higher levels of segregation distortion. In the reciprocal translocation with inverted segments (Fig. 4b), distorted markers were mainly concentrated within the translocated regions, consistent with impaired pairing and abnormal segregation during meiosis, as previously reported in *Musa* (Martin et al., 2017). In the more complex rearrangement, distortion was particularly severe near breakpoint regions, especially on chromosome 15 of the ALF genome and chromosome 24 of the CAD genome (Fig. 4c), suggesting strong constraints on homologous pairing and segregation.

Structural disparities between karyotypes can result in incompatible meiotic configurations in hybrids, producing inviable or low-fitness offspring, and thereby promoting genetic divergence and, ultimately, speciation. In *Carex*, chromosomal hybrid dysfunction (through incompatibility to form F_1_s) has been documented both within and between closely related species (Escudero et al., 2016b; Whitkus, 1988). Strong effects on segregation distortion in F_2_s from intraspecific crosses with different chromosome numbers have also been reported (Escudero et al., 2018). Although hybrid dysfunction has been proposed as a mechanism driving, at least partially, karyotype differentiation in *C. laevigata* (Escudero et al., 2013b; Márquez-Corro et al., 2024), there was no empirical evidence supporting this hypothesis. Our results indicate that, despite the inherent flexibility of holocentric chromosomes, hybrid dysfunction in *C. laevigata* reduces gamete viability in crosses between divergent karyotypes, generating postzygotic barriers and limiting gene flow, thereby reinforcing differentiation among populations.

### Recombination and CRs

The observed mean recombination rate across genomes (6.5-7 cM/Mb) falls within the expected range for holocentric organisms with 38-40 chromosomes (Zedek et al., 2026). Recombination rate was negatively associated with chromosome size (Fig. 3c; Supplementary Figs. 14-16), likely reflecting crossover interference and the generally low number of COs per chromosome in holocentric taxa (Nokkala et al., 2004; Hunter, 2015), which proportionally constrains recombination in larger chromosomes. Consistent with this, larger chromosomes showed elevated recombination toward telomeric regions, whereas smaller chromosomes exhibited a more uniform recombination landscape (Supplementary Figs. 9-11), potentially due to spatial constraints on crossover placement imposed by interference. Rearranged chromosomes also exhibited higher overall recombination rates than non-rearranged chromosomes, although these differences were not statistically significant (Fig. 3b; Supplementary Table 10). Nevertheless, rearranged chromosomes consistently showed significantly greater variation in recombination rate across all reference genomes examined (Fig. 3b; Supplementary Table 10). This pattern may result from multiple factors. Segregation distortion can inflate genetic map lengths and thus increase apparent recombination rates (Hackett and Broadfoot, 2003). In addition, CRs likely constrain crossover distribution during meiosis, restricting recombination to regions that retain sufficient homology for proper chromosomal pairing and segregation, while suppressing crossover formation in structurally divergent regions. As a result, recombination may become concentrated in a smaller subset of chromosomal regions, leading to greater heterogeneity in recombination rates along rearranged chromosomes.

The heterogeneous recombination landscapes observed across rearranged chromosomes may be driven by meiotic mechanical constraints that alter crossover distribution (Fig. 4; Supplementary Figs. 9–11). Near chromosomal breakpoints and within inverted segments, recombination may be reduced due to impaired synapsis and incomplete homologous pairing, which can hinder stable formation of the synaptonemal complex (Li et al., 2023) and contribute to the pronounced segregation distortion observed in these regions (Fig. 4). In fusion/fission and translocation heterozygotes, regular bivalent pairing is not possible, making multivalent configurations (trivalents and tetravalents) the preferred alternative to univalents to ensure balanced segregation. However, the formation of these complex structures may impose additional spatial constraints on crossover placement, favoring recombination in subterminal rather than interstitial regions (Strickberger, 1982; Lukhtanov et al., 2018). Consequently, the observed reduction in recombination in rearranged regions likely reflects a combination of suppressed crossover formation in structurally mismatched regions and selection against unbalanced recombinant gametes generated within rearranged chromosomal configurations (Wellenreuther & Bernatchez, 2018).

Collectively, our results demonstrate that CRs in *C. laevigata* locally suppress recombination, supporting the hypothesis that such structural shifts are primary drivers of karyotype differentiation (Faria and Navarro, 2010). While the theoretical framework for recombination suppression has focused on inversions (e.g., Faria et al., 2019; Huang and Riesberg, 2020), our work supports the growing evidence that diverse rearrangements, including fissions, fusions, and reciprocal translocations, exert similar constraints on genetic exchange (Martin et al., 2020; Vara et al., 2021). These findings are particularly significant for holocentric organisms, where empirical evidence of suppression of recombination remains scarce and has only recently been documented in few taxa such as nematodes (*Pristionchus*; Yoshida et al., 2023) and butterflies (*Leptidea sinapis*; Näsvall et al., 2023). By confirming these patterns in *C. laevigata*, we provide experimental support for the idea that recombination suppression is a universal consequence of chromosomal evolution.

### Adaptive Potential of CRs: Insights from QTL Mapping

The QTL mapping results reveal a complex genetic architecture underlying reproductive success in *C. laevigata*. In CAD × ALF individuals, the QTL profiles of chromosome 16 in the ALF genome and chromosome 40 in the CAD genome were highly similar, consistent with their syntenic relationship (Figs. 4c and 5a; Supplementary Table 12; Supplementary Figs. 19 and 20). Given the strong correlation between phenotypic traits (Supplementary Fig. 18), this QTL likely reflects either pleiotropic effects of a single gene or a cluster of tightly linked loci controlling inflorescence development. In both cases, QTLs represent a key component of the genetic architecture underlying reproductive success in *C. laevigata*. Despite the correspondence between QTLs in the ALF and CAD genomes, their genetic architectures differ: the ALF locus shows a stronger dominance effect of the paternal allele, whereas the CAD locus displays a predominantly additive pattern (Fig. 5b). These differences may reflect background-dependent allele expression or divergent selective histories, although the limited sample size (N = 48) in the CAD × ALF F_2_s limits the power to distinguish between these effects. Notably, the marker associated with these QTLs lies at ∼ 2 Mb from the breakpoint of a chromosomal inversion. Although recombination is not completely suppressed in this region (Fig. 4c), the presence of moderate segregation distortion suggests that the locus remains influenced by the rearrangement. This effect may arise from reduced recombination near the breakpoint, linkage with adaptive alleles, or post-zygotic selection acting against recombinant genotypes.

The genetic architecture of the CAD × HUE locus reveals a distinct evolutionary signature. Unlike the CAD × ALF loci, which show dominance or additive effects, the CAD × HUE locus on chromosome 4 exhibits a pattern of underdominance, where individuals with homozygous genotypes (particularly the HUE parent alleles) show significantly higher reproductive output than heterozygotes (Fig. 5b). This reduction in fitness among heterozygotes at a collinear locus is consistent with genetic incompatibility between the CAD and HUE. While the CAD × ALF reproductive success appears influenced by the structural constraints of chromosomal inversions and segregation distortion, the CAD × HUE cross points toward intrinsic post-zygotic barriers. This type of incompatibility, where hybrid genotypes are less fit than parental ones, is a hallmark of diverging lineages (Mack & Nachman, 2017) and suggests that even in the absence of CRs, strong selective barriers are maintaining the integrity of these genomic regions.

CRs in *C. laevigata* have been suggested to promote local adaptation (Escudero et al., 2013b; Márquez-Corro et al., 2024). Chromosomal inversions are well known to suppress recombination and facilitate adaptive divergence, often acting as “supergenes” that protect clusters of co-adapted genes (Ayala and Coluzzi, 2005; Fransz et al. 2016; Todesco et al., 2020; Kollar, 2025). Our results in CAD × ALF cross support this, as the proximity to the inversion breakpoint likely prevents the breakdown of favorable allele combinations. However, the pattern observed in the CAD × HUE cross provides an important contrast, demonstrating that strong barriers to gene flow can also arise in non-rearranged, collinear genomic regions through fitness-related incompatibilities (Reifová et al., 2023). Together, these results suggest that the adaptive landscape of *C. laevigata* is shaped by a mosaic of CRs and specific incompatible loci that limit hybrid fitness.

## Conclusions

Ultimately, the exceptional karyotypic diversity observed in *Carex laevigata* is driven by a complex interplay of inherent genomic vulnerabilities and different evolutionary forces. Our genomic characterization reveals that structural variation extends far beyond simple fusion and fission events, with a hidden landscape of CRs including inversions and reciprocal translocations catalyzed by GC-rich hotspots and LTR-Gypsy retrotransposons. Crucially, while the holocentric nature of *Carex* chromosomes provides some meiotic flexibility, these structural rearrangements act as a strong barrier to gene flow.

By disrupting homologous pairing, causing severe segregation distortion, and suppressing recombination—particularly around breakpoints and within complex structural shifts—these CRs drive significant hybrid dysfunction. Furthermore, the discovery of key reproductive QTLs linked to inversion breakpoints underscores the adaptive potential of these regions. Taken together, our findings corroborate theoretical predictions (Lucek et al., 2022) by providing compelling empirical evidence that CRs are not merely byproducts of genomic instability, but are primary architects of reproductive isolation, karyotypic differentiation, and speciation within holocentric lineages.

## Acknowledgements

This study was funded by the Andalusian regional government (Junta de Andalucía, Spain) through project ProyExcel-00125, and the Agencia Estatal de Investigación-Ministerio de Ciencia, Innovación y Universidades (AEI-MICIU) through projects PID2021-122715NB-I00 and PID2024-157198NB-I00. RSV was supported by projects ProyExcel-00125 and AEI-MICIU PID2023-147332NB-I00. IGR was supported by the Andalusian Regional Ministry of Economy, Knowledge, Business, and University (PREDOC_00632). AVM and KL were supported by the Swiss National Science Foundation (SNSF) grants 202869 and 220868, the Fondation Pierre Mercier pour la science and the Fondation du Jardin Botanique de Neuchâtel. The authors thank C. Cornet and P. Escuer for invaluable discussions. F.J. Fernández, A. Montero, M. Sanz-Arnal, M. Sánchez-Villegas, P. Pineda and C. Barciela-Torres for their help with greenhouse work. The computations and simulations presented in this study were carried out using the Hércules supercomputer at the Centro Informático Científico de Andalucía (CICA), managed by the Agencia Digital de Andalucía.

