## Supplementary information for "Dynamic holocentric genomes facilitate divergent evolutionary paths through chromosomal rearrangements, hybrid dysfunction, and recombination suppression"

**Supplementary Tables**

**Supplementary Table 1. Germination rate of selfed parents, F₁ and F₂ individuals.** Number of seeds collected, number of germinated, and the resulting germination rates for selfed parental individuals (ALF, CAD, and HUE) and for F₂ offspring derived from crosses between these populations. Each row corresponds to an independent cross.

| **Cross id** | **Cross type** | **Population** | **Seeds collected** | **Seeds germinated** | **Germination rate** |
| --- | --- | --- | --- | --- | --- |
| 1 | selfing P | ALF | 44 | 40 | 0.909 |
| 2 | selfing P | ALF | 71 | 17 | 0.239 |
| 3 | selfing P | ALF | 57 | 28 | 0.491 |
| 4 | selfing P | ALF | 52 | 48 | 0.923 |
| 5 | selfing P | ALF | 93 | 13 | 0.140 |
| 6 | selfing P | ALF | 49 | 31 | 0.633 |
| 7 | selfing P | ALF | 48 | 0 | 0.000 |
| 8 | selfing P | CAD | 50 | 36 | 0.720 |
| 9 | selfing P | CAD | 69 | 60 | 0.870 |
| 10 | selfing P | CAD | 12 | 11 | 0.917 |
| 11 | selfing P | CAD | 54 | 50 | 0.926 |
| 12 | selfing P | CAD | 71 | 58 | 0.817 |
| 13 | selfing P | CAD | 50 | 0 | 0.000 |
| 14 | selfing P | HUE | 112 | 78 | 0.696 |
| 15 | selfing P | HUE | 58 | 4 | 0.069 |
| 16 | selfing P | HUE | 164 | 5 | 0.030 |
| 17 | selfing P | HUE | 211 | 94 | 0.445 |
| 18 | F1 (crossing P) | CAD x ALF | 29 | 14 | 0.483 |
| 19 | F1 (crossing P) | CAD x ALF | 34 | 14 | 0.412 |
| 20 | F1 (crossing P) | CAD x ALF | 54 | 1 | 0.019 |
| 21 | F1 (crossing P) | CAD x ALF | 48 | 0 | 0.000 |
| 22 | F1 (crossing P) | CAD x ALF | 10 | 3 | 0.300 |
| 23 | F1 (crossing P) | CAD x HUE | 61 | 56 | 0.918 |
| 24 | F1 (crossing P) | CAD x HUE | 50 | 31 | 0.620 |
| 25 | F1 (crossing P) | CAD x HUE | 42 | 35 | 0.833 |
| 26 | F1 (crossing P) | CAD x HUE | 65 | 0 | 0.000 |
| 27 | F2 (selfing F1) | CAD x ALF | 123 | 13 | 0.106 |
| 28 | F2 (selfing F1) | CAD x ALF | 76 | 46 | 0.605 |
| 29 | F2 (selfing F1) | CAD x ALF | 43 | 3 | 0.070 |
| 30 | F2 (selfing F1) | CAD x ALF | 56 | 25 | 0.446 |
| 31 | F2 (selfing F1) | CAD x ALF | 35 | 0 | 0.000 |
| 32 | F2 (selfing F1) | CAD x ALF | 163 | 5 | 0.031 |
| 33 | F2 (selfing F1) | CAD x ALF | 56 | 1 | 0.018 |
| 34 | F2 (selfing F1) | CAD x ALF | 112 | 18 | 0.161 |
| 35 | F2 (selfing F1) | CAD x ALF | 46 | 0 | 0.000 |
| 36 | F2 (selfing F1) | CAD x ALF | 82 | 14 | 0.171 |
| 37 | F2 (selfing F1) | CAD x ALF | 85 | 0 | 0.000 |
| 38 | F2 (selfing F1) | CAD x ALF | 132 | 0 | 0.000 |
| 39 | F2 (selfing F1) | CAD x ALF | 98 | 1 | 0.010 |
| 40 | F2 (selfing F1) | CAD x ALF | 98 | 0 | 0.000 |
| 41 | F2 (selfing F1) | CAD x HUE | 64 | 1 | 0.016 |
| 42 | F2 (selfing F1) | CAD x HUE | 175 | 0 | 0.000 |
| 43 | F2 (selfing F1) | CAD x HUE | 43 | 2 | 0.047 |
| 44 | F2 (selfing F1) | CAD x HUE | 26 | 3 | 0.115 |
| 45 | F2 (selfing F1) | CAD x HUE | 124 | 0 | 0.000 |
| 46 | F2 (selfing F1) | CAD x HUE | 87 | 0 | 0.000 |
| 47 | F2 (selfing F1) | CAD x HUE | 22 | 5 | 0.227 |
| 48 | F2 (selfing F1) | CAD x HUE | 26 | 14 | 0.538 |
| 49 | F2 (selfing F1) | CAD x HUE | 109 | 48 | 0.440 |
| 50 | F2 (selfing F1) | CAD x HUE | 66 | 3 | 0.045 |
| 51 | F2 (selfing F1) | CAD x HUE | 144 | 7 | 0.049 |
| 52 | F2 (selfing F1) | CAD x HUE | 83 | 1 | 0.012 |
| 53 | F2 (selfing F1) | CAD x HUE | 89 | 0 | 0.000 |
| 54 | F2 (selfing F1) | CAD x HUE | 55 | 6 | 0.109 |
| 55 | F2 (selfing F1) | CAD x HUE | 53 | 28 | 0.528 |
| 56 | F2 (selfing F1) | CAD x HUE | 77 | 38 | 0.494 |
| 57 | F2 (selfing F1) | CAD x HUE | 23 | 6 | 0.261 |
| 58 | F2 (selfing F1) | CAD x HUE | 52 | 0 | 0.000 |
| 59 | F2 (selfing F1) | CAD x HUE | 60 | 7 | 0.117 |
| 60 | F2 (selfing F1) | CAD x HUE | 50 | 1 | 0.020 |
| 61 | F2 (selfing F1) | CAD x HUE | 45 | 0 | 0.000 |
| 62 | F2 (selfing F1) | CAD x HUE | 52 | 0 | 0.000 |
| 63 | F2 (selfing F1) | CAD x HUE | 51 | 6 | 0.118 |
| 64 | F2 (selfing F1) | CAD x HUE | 53 | 26 | 0.491 |
| 65 | F2 (selfing F1) | CAD x HUE | 11 | 4 | 0.364 |
| 66 | F2 (selfing F1) | CAD x HUE | 52 | 4 | 0.077 |
| 67 | F2 (selfing F1) | CAD x HUE | 61 | 17 | 0.279 |
| 68 | F2 (selfing F1) | CAD x HUE | 64 | 21 | 0.328 |
| 69 | F2 (selfing F1) | CAD x HUE | 167 | 0 | 0.000 |
| 70 | F2 (selfing F1) | CAD x HUE | 24 | 14 | 0.583 |
| 71 | F2 (selfing F1) | CAD x HUE | 22 | 0 | 0.000 |
| 72 | F2 (selfing F1) | CAD x HUE | 33 | 23 | 0.697 |
| 73 | F2 (selfing F1) | CAD x HUE | 68 | 9 | 0.132 |
| 74 | F2 (selfing F1) | CAD x HUE | 122 | 2 | 0.016 |
| 75 | F2 (selfing F1) | CAD x HUE | 57 | 36 | 0.632 |
| 76 | F2 (selfing F1) | CAD x HUE | 34 | 1 | 0.029 |
| 77 | F2 (selfing F1) | CAD x HUE | 77 | 9 | 0.117 |
| 78 | F2 (selfing F1) | CAD x HUE | 52 | 4 | 0.077 |
| 79 | F2 (selfing F1) | CAD x HUE | 177 | 43 | 0.243 |
| 80 | F2 (selfing F1) | CAD x HUE | 125 | 8 | 0.064 |
| 81 | F2 (selfing F1) | CAD x HUE | 43 | 12 | 0.279 |
| 82 | F2 (selfing F1) | CAD x HUE | 121 | 14 | 0.116 |
| 83 | F2 (selfing F1) | CAD x HUE | 30 | 18 | 0.600 |

**Supplementary Table 2. Enrichment analysis of genomic features in breakpoint regions of the DToL, ALF and CAD genomes.** Results from permutation-based enrichment tests comparing genomic windows classified as *breakpoints* versus *syntenic* regions. For each feature or repeat family, the table reports the *p*-value and *z*-score of the enrichment test, where positive *z*-scores indicate relative enrichment in breakpoint regions and negative *z*-scores indicate depletion. Two analyses were conducted: one including chromosomal ends and one excluding them. In the EDI genome, all identified breakpoints were located within chromosomes. “Repeat element” entries correspond to specific repeat families annotated in the genome. Significance codes: *p* < 0.05 (*), *p* < 0.01 (**), *p* < 0.001 (***), *p* < 0.10 (.).

| **Genome** | **Genomic feature** | **Repeat class/family** | **Including chromosomal ends** | | **Excluding chromosomal ends** | |
| --- | --- | --- | --- | --- | --- | --- |
|  |  |  | ***p-value*** | ***z-score*** | ***p-value*** | ***z-score*** |
| EDI | CDS | - | - | - | 0.008 ** | -2.094 |
| EDI | GC proportion | - | - | - | 0.058 . | 1.532 |
| EDI | Repeat elements | - | - | - | 0.094 . | -1.229 |
| EDI | Repeat element | DNA | - | - | 0.884 | -0.347 |
| EDI | Repeat element | DNA/CMC-EnSpm | - | - | 0.145 | 1.059 |
| EDI | Repeat element | DNA/hAT | - | - | 0.21 | 0.996 |
| EDI | Repeat element | DNA/hAT-Ac | - | - | 0.042 * | 2.128 |
| EDI | Repeat element | DNA/hAT-Charlie | - | - | 0.782 | -0.465 |
| EDI | Repeat element | DNA/hAT-Tag1 | - | - | 0.013 * | 3.439 |
| EDI | Repeat element | DNA/hAT-Tip100 | - | - | 0.491 | -0.392 |
| EDI | Repeat element | DNA/MULE-MuDR | - | - | 0.016 * | -1.829 |
| EDI | Repeat element | DNA/PIF-Harbinger | - | - | 0.37 | 0.479 |
| EDI | Repeat element | DNA/TcMar | - | - | 0.1 | 1.403 |
| EDI | Repeat element | DNA/Zisupton | - | - | 0.438 | -0.298 |
| EDI | Repeat element | LINE/I | - | - | 0.794 | -0.467 |
| EDI | Repeat element | LINE/L1 | - | - | 0.364 | 0.38 |
| EDI | Repeat element | LINE/L2 | - | - | 0.825 | -0.413 |
| EDI | Repeat element | LINE/RTE | - | - | 0.6 . | -0.669 |
| EDI | Repeat element | LINE/RTE-BovB | - | - | 0.453 | -0.434 |
| EDI | Repeat element | Low_complexity | - | - | 0.473 | -0.125 |
| EDI | Repeat element | LTR | - | - | 0.909 | -0.297 |
| EDI | Repeat element | LTR/Copia | - | - | 0.348 | 0.395 |
| EDI | Repeat element | LTR/ERVK | - | - | 0.276 | 0.959 |
| EDI | Repeat element | LTR/ERVL | - | - | 0.969 | -0.127 |
| EDI | Repeat element | LTR/Gypsy | - | - | 0.082 . | 1.591 |
| EDI | Repeat element | LTR/Pao | - | - | 0.937 | -0.162 |
| EDI | Repeat element | RC/Helitron | - | - | 0.115 | -1.158 |
| EDI | Repeat element | Satellite | - | - | 0.052 . | -1.493 |
| EDI | Repeat element | Simple_repeat | - | - | 0.161 | -0.903 |
| EDI | Repeat element | SINE/U | - | - | 0.936 | -0.238 |
| EDI | Repeat element | Unknown | - | - | 0.08 . | -1.381 |
| ALF | CDS | - | 0.002 ** | -2.923 | < 0.001 *** | -3.309 |
| ALF | GC proportion | - | < 0.001 *** | 8.643 | < 0.001 *** | 7.422 |
| ALF | Repeat elements | - | 0.411 | -0.216 | 0.302 | 0.492 |
| ALF | Repeat element | DNA | 0.241 | -0.928 | 0.421 | -0.462 |
| ALF | Repeat element | DNA/CMC-EnSpm | 0.539 | -0.063 | 0.468 | -0.293 |
| ALF | Repeat element | DNA/hAT-Ac | 0.424 | -0.330 | 0.395 | -0.423 |
| ALF | Repeat element | DNA/hAT-Tag1 | 0.286 | 0.667 | 0.217 | 0.971 |
| ALF | Repeat element | DNA/hAT-Tip100 | 0.585 | -0.080 | 0.414 | 0.281 |
| ALF | Repeat element | DNA/MULE-MuDR | 0.168 | -1.005 | 0.455 | -0.209 |
| ALF | Repeat element | DNA/PIF-Harbinger | 0.523 | -0.346 | 0.629 | -0.123 |
| ALF | Repeat element | DNA/TcMar | 0.328 | 0.467 | 0.436 | -0.392 |
| ALF | Repeat element | DNA/Zisupton | 0.339 | -0.584 | 0.454 | -0.293 |
| ALF | Repeat element | LINE/I | 0.596 | -0.671 | 0.67 | -0.605 |
| ALF | Repeat element | LINE/L1 | 0.176 | -0.974 | 0.357 | -0.47 |
| ALF | Repeat element | LINE/L1-Tx1 | 0.730 | -0.544 | 0.786 | -0.471 |
| ALF | Repeat element | LINE/RTE-BovB | 0.272 | -0.300 | 0.367 | -0.189 |
| ALF | Repeat element | Low_complexity | 0.243 | 0.730 | 0.29 | 0.55 |
| ALF | Repeat element | LTR | 0.110 | -1.304 | 0.219 | -0.894 |
| ALF | Repeat element | LTR/Copia | 0.401 | -0.343 | 0.439 | 0.113 |
| ALF | Repeat element | LTR/ERV1 | 0.372 | -0.887 | 0.448 | -0.822 |
| ALF | Repeat element | LTR/ERVK | 0.932 | -0.155 | 0.931 | -0.15 |
| ALF | Repeat element | LTR/Gypsy | 0.046 * | 1.899 | 0.064 . | 1.692 |
| ALF | Repeat element | RC/Helitron | 0.286 | -0.626 | 0.423 | 0.085 |
| ALF | Repeat element | Satellite | 0.172 | 0.997 | 0.06 | 1.65 |
| ALF | Repeat element | Simple_repeat | 0.213 | -0.747 | 0.351 | -0.43 |
| ALF | Repeat element | SINE/U | 0.886 | -0.297 | 0.912 | -0.291 |
| ALF | Repeat element | Unknown | 0.453 | 0.109 | 0.225 | 0.735 |
| CAD | CDS | - | 0.453 | -0.162 | 0.011 * | -2.086 |
| CAD | GC proportion | - | < 0.001 *** | 9.341 | 0.015 * | 2.82 |
| CAD | Repeat elements | - | < 0.001 *** | -5.603 | < 0.001 *** | -4.494 |
| CAD | Repeat element | DNA | 0.498 | -0.520 | 0.509 | 0.196 |
| CAD | Repeat element | DNA/CMC-EnSpm | 0.102 | 1.397 | 0.474 | -0.255 |
| CAD | Repeat element | DNA/hAT | 0.342 | 0.579 | 0.351 | 0.657 |
| CAD | Repeat element | DNA/hAT-Ac | 0.025 * | -1.800 | 0.151 | -1.127 |
| CAD | Repeat element | DNA/hAT-Tag1 | 0.124 | -1.262 | 0.383 | -0.84 |
| CAD | Repeat element | DNA/hAT-Tip100 | 0.436 | -0.460 | 0.238 | -1.087 |
| CAD | Repeat element | DNA/Maverick | 0.014 | -2.141 | 0.023 | -1.765 |
| CAD | Repeat element | DNA/MULE-MuDR | 0.017 | -2.229 | 0.517 | -0.127 |
| CAD | Repeat element | DNA/PIF-Harbinger | 0.147 | -1.186 | 0.456 | -0.32 |
| CAD | Repeat element | DNA/TcMar-Tigger | 0.470 | -0.815 | 0.726 | -0.52 |
| CAD | Repeat element | DNA/Zisupton | 0.226 | -0.852 | 0.396 | -0.472 |
| CAD | Repeat element | LINE/I | 0.141 | -1.349 | 0.417 | -0.9 |
| CAD | Repeat element | LINE/L1 | 0.005 ** | -2.512 | 0.289 | -0.669 |
| CAD | Repeat element | LINE/RTE-BovB | 0.367 | -0.327 | 0.618 | -0.113 |
| CAD | Repeat element | LTR | 0.233 | -1.173 | 0.549 | -0.737 |
| CAD | Repeat element | LTR/Copia | 0.015 * | -2.102 | 0.33 | 0.439 |
| CAD | Repeat element | LTR/ERV1 | 0.656 | -0.237 | 0.331 | 0.385 |
| CAD | Repeat element | LTR/ERVK | 0.475 | -0.379 | 0.475 | 0.26 |
| CAD | Repeat element | LTR/ERVL | 0.219 | 1.042 | 0.333 | 0.798 |
| CAD | Repeat element | LTR/Gypsy | 0.024 * | 2.094 | 0.007 ** | 3.477 |
| CAD | Repeat element | PLE/Chlamys | 0.550 | 0.019 | 0.64 | -0.242 |
| CAD | Repeat element | RC/Helitron | 0.004 ** | -2.485 | 0.127 | -1.189 |
| CAD | Repeat element | Satellite | 0.447 | -0.269 | 0.364 | 0.365 |
| CAD | Repeat element | Unknown | < 0.001 *** | -4.336 | 0.124 | -1.201 |

**Supplementary Table 3. Fine-grained enrichment analysis of individual breakpoint regions in the DToL genome.** Values represent the Z-score and the empirical p-value (in parentheses) obtained from 1,000 permutations using *regioneR*. Positive Z-scores indicate enrichment (presence higher than expected), while negative Z-scores indicate depletion of the genomic feature within the breakpoint interval. CDS: Coding Sequence; GC: GC content; bp: base pairs; NA: feature not present in the randomized background for that specific interval. Significance codes: *p* < 0.05 (*), *p* < 0.01 (**), *p* < 0.001 (***), *p* < 0.10 (.).

| **Chromosome** | 1 | 5 | 6 | 6 | 6 | 8 | 8 | 8 | 18 | 18 | 27 |
| --- | --- | --- | --- | --- | --- | --- | --- | --- | --- | --- | --- |
| **Start bp** | 7,877,580 | 1,466,098 | 4,232,133 | 5,613,930 | 7,939,300 | 8,746,215 | 9,910,730 | 11,792,160 | 5,481,238 | 9,688,749 | 4,205,702 |
| **End bp** | 7,960,197 | 1,467,833 | 4,249,085 | 5,633,656 | 7,963,241 | 8,755,674 | 9,986,028 | 11,803,536 | 5,501,934 | 9,818,192 | 4,455,734 |
| **CDS** | -0.564 (0.319) | 0.112 (0.378) | -0.144 (0.574) | -0.434 (0.398) | 0.028 (0.402) | -0.676 (0.364) | -0.853 (0.2) | -0.845 (0.266) | -0.649 (0.323) | -0.502 (0.342) | -1.547 (0.051.) |
| **GC** | 2.479 (0.02*) | -0.245 (0.437) | 0.344 (0.297) | -0.818 (0.206) | 0.137 (0.418) | 0.187 (0.392) | 0.494 (0.287) | 0.245 (0.356) | -0.598 (0.287) | 3.248 (0.011*) | -1.317 (0.075.) |
| **All Repeats** | -0.396 (0.323) | 1.018 (0.21) | 1.042 (0.164) | -0.097 (0.478) | 0.142 (0.478) | 0.959 (0.197) | -0.202 (0.42) | -1.048 (0.161) | 0.389 (0.377) | -2.724 (0.007**) | 0.147 (0.422) |
| **DNA** | -0.131 (0.983) | -0.032 (0.999) | -0.032 (0.999) | -0.084 (0.993) | -0.067 (0.995) | -0.063 (0.996) | -0.128 (0.984) | -0.045 (0.998) | -0.1 (0.99) | -0.123 (0.985) | -0.209 (0.954) |
| **DNA/CMC-EnSpm** | -0.622 (0.445) | -0.166 (0.97) | 0.805 (0.185) | -0.41 (0.777) | 0.592 (0.256) | -0.29 (0.888) | -0.128 (0.729) | -0.27 (0.878) | -0.383 (0.794) | 3.34 (0.015*) | -0.543 (0.412) |
| **DNA/MULE-MuDR** | -0.721 (0.343) | -0.277 (0.917) | 0.811 (0.222) | -0.703 (0.522) | -0.172 (0.677) | -0.509 (0.693) | -0.659 (0.353) | -0.554 (0.658) | -0.717 (0.491) | -1.22 (0.132) | -0.909 (0.199) |
| **DNA/PIF-Harbinger** | -0.344 (0.87) | -0.06 (0.996) | -0.171 (0.968) | -0.157 (0.971) | 4.114 (0.034*) | -0.12 (0.983) | -0.33 (0.879) | -0.145 (0.976) | 5.032 (0.03*) | -0.421 (0.809) | -0.611 (0.62) |
| **DNA/TcMar** | 6.499 (0.003**) | -0.063 (0.996) | -0.283 (0.918) | -0.288 (0.912) | -0.348 (0.882) | -0.222 (0.95) | 1.002 (0.304) | -0.249 (0.935) | -0.306 (0.9) | -0.668 (0.53) | -0.242 (0.624) |
| **DNA/Zisupton** | -0.733 (0.38) | -0.254 (0.933) | 0.674 (0.339) | -0.614 (0.617) | 0.348 (0.419) | 1.265 (0.222) | -0.486 (0.408) | -0.503 (0.749) | 0.29 (0.404) | -1.108 (0.154) | 0.654 (0.242) |
| **DNA/hAT** | -0.494 (0.756) | -0.105 (0.989) | -0.236 (0.944) | -0.263 (0.931) | 2.688 (0.091.) | -0.138 (0.98) | -0.443 (0.799) | -0.199 (0.959) | -0.298 (0.913) | 0.679 (0.347) | 0.878 (0.252) |
| **DNA/hAT-Ac** | 6.441 (0.003**) | -0.161 (0.971) | -0.317 (0.884) | -0.337 (0.863) | -0.411 (0.807) | -0.261 (0.919) | 0.735 (0.24) | -0.279 (0.907) | -0.372 (0.835) | -0.799 (0.408) | 0.04 (0.455) |
| **DNA/hAT-Charlie** | -0.154 (0.975) | NA (1) | -0.097 (0.99) | -0.107 (0.988) | -0.12 (0.985) | -0.053 (0.997) | -0.16 (0.972) | -0.063 (0.996) | -0.071 (0.995) | -0.18 (0.967) | -0.3 (0.91) |
| **DNA/hAT-Tag1** | -0.241 (0.927) | -0.052 (0.997) | -0.09 (0.991) | -0.115 (0.983) | -0.135 (0.978) | -0.1 (0.989) | -0.258 (0.916) | -0.113 (0.985) | -0.141 (0.976) | 5.549 (0.014*) | 2.044 (0.07.) |
| **DNA/hAT-Tip100** | 0.78 (0.231) | -0.09 (0.992) | -0.244 (0.93) | -0.238 (0.937) | -0.24 (0.926) | -0.149 (0.976) | -0.47 (0.754) | -0.175 (0.956) | -0.212 (0.941) | -0.519 (0.641) | -0.082 (0.723) |
| **LINE/I** | -0.15 (0.978) | NA (1) | -0.042 (0.998) | -0.063 (0.996) | -0.086 (0.992) | -0.045 (0.998) | -0.176 (0.97) | -0.055 (0.997) | -0.078 (0.994) | -0.197 (0.961) | -0.255 (0.938) |
| **LINE/L1** | -1.387 (0.092.) | -0.323 (0.883) | 0.49 (0.324) | 2.169 (0.065.) | 0.769 (0.256) | -0.62 (0.619) | 0.833 (0.22) | -0.678 (0.556) | 2.253 (0.056.) | -1.496 (0.056.) | 0.64 (0.278) |
| **LINE/L2** | -0.168 (0.971) | NA (1) | -0.063 (0.996) | -0.073 (0.994) | -0.102 (0.989) | -0.063 (0.996) | -0.131 (0.978) | -0.071 (0.995) | -0.09 (0.992) | -0.182 (0.961) | -0.24 (0.927) |
| **LINE/RTE** | -0.225 (0.945) | -0.032 (0.999) | -0.128 (0.982) | -0.124 (0.983) | -0.131 (0.98) | -0.071 (0.995) | -0.199 (0.956) | -0.084 (0.993) | -0.108 (0.987) | -0.298 (0.907) | -0.411 (0.831) |
| **LINE/RTE-BovB** | -0.437 (0.806) | -0.071 (0.995) | -0.222 (0.95) | -0.076 (0.948) | -0.261 (0.93) | -0.157 (0.975) | -0.434 (0.814) | -0.189 (0.964) | -0.235 (0.944) | -0.534 (0.694) | -0.027 (0.806) |
| **LTR** | -0.111 (0.987) | NA (1) | -0.045 (0.998) | -0.055 (0.997) | -0.055 (0.997) | -0.055 (0.997) | -0.128 (0.983) | -0.045 (0.998) | -0.055 (0.997) | -0.114 (0.984) | -0.172 (0.967) |
| **LTR/Copia** | -0.208 (0.552) | -0.441 (0.808) | 0.416 (0.329) | 0.821 (0.225) | 0.643 (0.26) | -0.667 (0.555) | 1.53 (0.099.) | 0.108 (0.467) | -0.344 (0.593) | -0.908 (0.216) | 0.378 (0.338) |
| **LTR/ERVK** | -0.343 (0.879) | -0.055 (0.997) | -0.138 (0.98) | -0.157 (0.972) | -0.169 (0.968) | -0.14 (0.98) | -0.326 (0.895) | -0.143 (0.979) | -0.179 (0.968) | -0.417 (0.827) | 2.454 (0.074.) |
| **LTR/ERVL** | NA (1) | NA (1) | -0.048 (0.997) | NA (1) | -0.048 (0.997) | NA (1) | -0.032 (0.999) | NA (1) | -0.036 (0.998) | -0.062 (0.996) | -0.071 (0.993) |
| **LTR/Gypsy** | 0.324 (0.317) | -0.31 (0.903) | -0.434 (0.661) | 0.248 (0.364) | -0.593 (0.59) | -0.425 (0.776) | -0.193 (0.622) | -0.483 (0.731) | 0.143 (0.372) | 2.225 (0.037*) | 1.038 (0.151) |
| **LTR/Pao** | -0.051 (0.995) | NA (1) | -0.06 (0.996) | -0.063 (0.996) | -0.042 (0.998) | NA (1) | -0.05 (0.994) | NA (1) | -0.039 (0.998) | -0.062 (0.986) | -0.085 (0.972) |
| **Low complexity** | 1.705 (0.07.) | -0.284 (0.92) | -0.772 (0.517) | -0.01 (0.735) | 2.554 (0.041*) | -0.466 (0.693) | -0.604 (0.401) | 0.566 (0.38) | 1.134 (0.213) | -0.589 (0.142) | -0.486 (0.292) |
| **RC/Helitron** | -0.737 (0.334) | -0.238 (0.937) | -0.618 (0.617) | 0.285 (0.419) | -0.721 (0.513) | -0.507 (0.746) | -0.713 (0.359) | -0.521 (0.721) | 1.247 (0.149) | -0.74 (0.27) | -0.376 (0.429) |
| **SINE/U** | -0.097 (0.99) | NA (1) | NA (1) | -0.045 (0.998) | -0.052 (0.997) | -0.045 (0.998) | -0.042 (0.998) | NA (1) | -0.045 (0.998) | -0.084 (0.992) | -0.155 (0.972) |
| **Satellite** | 1.487 (0.111) | -0.357 (0.856) | -0.089 (0.693) | -0.204 (0.645) | -0.401 (0.552) | -0.688 (0.572) | -0.722 (0.182) | -0.766 (0.501) | -0.563 (0.29) | -1.168 (0.137) | -1.203 (0.121) |
| **Simple repeat** | -0.223 (0.458) | 0.863 (0.278) | 1.056 (0.163) | -0.852 (0.256) | -0.744 (0.289) | 0.357 (0.389) | 0.409 (0.328) | 0.135 (0.482) | 1.179 (0.129) | -1.63 (0.042*) | -0.614 (0.255) |
| **Unknown** | -0.884 (0.191) | 1.635 (0.111) | 0.756 (0.237) | -0.174 (0.457) | -0.023 (0.521) | 1.696 (0.065.) | -0.392 (0.352) | -0.942 (0.219) | -0.234 (0.447) | -3.106 (0.004**) | 0.42 (0.334) |

**Supplementary Table 4. Fine-grained enrichment analysis of individual breakpoint regions in the ALF genome.** Values represent the Z-score and the empirical p-value (in parentheses) obtained from 1,000 permutations using *regioneR*. Positive Z-scores indicate enrichment (presence higher than expected), while negative Z-scores indicate depletion of the genomic feature within the breakpoint interval. CDS: Coding Sequence; GC: GC content; bp: base pairs; NA: feature not present in the randomized background for that specific interval. Significance codes: *p* < 0.05 (*), *p* < 0.01 (**), *p* < 0.001 (***), *p* < 0.10 (.).

| **Chromosome** | 1 | 4 | 8 | 15 | 15 | 16 | 16 | 16 | 22 | 27 | 27 | 27 | 32 | 34 | 37 | 38 |
| --- | --- | --- | --- | --- | --- | --- | --- | --- | --- | --- | --- | --- | --- | --- | --- | --- |
| **Start bp** | 4,154,253 | 4,319,927 | 1 | 8,093,120 | 8,587,773 | 4,035,832 | 5,025,707 | 6,292,071 | 2,862,793 | 4,365,670 | 5,812,307 | 8,150,000 | 1,638,528 | 5,550,729 | 1 | 5,050,000 |
| **End bp** | 4,179,719 | 4,858,347 | 50,000 | 8,147,397 | 8,608,693 | 4,068,781 | 5,047,031 | 6,380,364 | 2,875,046 | 4,368,067 | 5,841,813 | 8,189,538 | 1,677,164 | 5,559,443 | 50,000 | 5,050,116 |
| **CDS** | -0.38 (0.469) | -2.703 (0.002**) | 0.11 (0.421) | -1.21 (0.092.) | -0.147 (0.591) | -0.373 (0.469) | -0.853 (0.288) | -1.037 (0.155) | -0.568 (0.45) | 0.034 (0.352) | -0.961 (0.189) | -0.195 (0.528) | -0.991 (0.166) | 1.466 (0.109) | 0.303 (0.357) | -0.352 (0.886) |
| **GC** | 1.422 (0.09.) | 8.617 (0.001***) | 1.724 (0.05*) | 0.238 (0.344) | 2.008 (0.036*) | -0.025 (0.543) | -0.159 (0.478) | -0.009 (0.556) | -1.824 (0.008**) | 0.396 (0.308) | -1.659 (0.027*) | 0.56 (0.237) | -0.849 (0.184) | -0.067 (0.526) | 1.779 (0.052.) | 5.064 (0.003**) |
| **All Repeats** | -1.232 (0.101) | 0.352 (0.342) | 1.129 (0.129) | 0.266 (0.404) | 1.224 (0.074.) | -0.04 (0.48) | -0.048 (0.484) | 0.307 (0.397) | 0.156 (0.449) | 0.493 (0.351) | 0.331 (0.389) | 0.183 (0.444) | -0.13 (0.448) | -0.787 (0.251) | -1.697 (0.058.) | 0.331 (0.666) |
| **DNA** | -0.394 (0.848) | 0.098 (0.497) | -0.507 (0.72) | -0.492 (0.718) | -0.302 (0.902) | -0.431 (0.811) | -0.293 (0.882) | -0.654 (0.563) | -0.269 (0.916) | -0.12 (0.983) | -0.408 (0.832) | -0.465 (0.787) | 1.278 (0.207) | -0.206 (0.957) | -0.521 (0.742) | -0.071 (0.995) |
| **DNA/CMC-EnSpm** | -0.519 (0.628) | -0.136 (0.537) | -0.779 (0.395) | -0.79 (0.378) | -0.527 (0.648) | 0.613 (0.213) | -0.504 (0.684) | -0.928 (0.199) | -0.42 (0.757) | -0.202 (0.949) | 0.081 (0.451) | 0.555 (0.253) | 2.588 (0.046*) | -0.357 (0.829) | -0.19 (0.681) | -0.15 (0.978) |
| **DNA/MULE-MuDR** | 2.073 (0.064.) | -0.524 (0.335) | -0.267 (0.572) | 0.411 (0.322) | 1.843 (0.08.) | 0.474 (0.291) | -0.725 (0.484) | -0.533 (0.397) | -0.633 (0.597) | -0.318 (0.874) | 0.189 (0.403) | -0.946 (0.27) | -0.518 (0.488) | -0.484 (0.737) | -1.09 (0.191) | -0.213 (0.955) |
| **DNA/PIF-Harbinger** | -0.228 (0.944) | -0.227 (0.647) | -0.264 (0.922) | -0.29 (0.901) | -0.213 (0.951) | 2.821 (0.059.) | -0.192 (0.958) | -0.387 (0.833) | -0.14 (0.979) | -0.078 (0.994) | -0.199 (0.953) | -0.263 (0.923) | -0.229 (0.941) | -0.107 (0.988) | -0.317 (0.891) | -0.063 (0.996) |
| **DNA/TcMar** | -0.452 (0.794) | -0.062 (0.619) | 5.74 (0.003**) | -0.649 (0.62) | -0.422 (0.82) | 1.135 (0.249) | -0.434 (0.817) | 0.149 (0.53) | -0.344 (0.884) | -0.148 (0.977) | -0.478 (0.761) | -0.533 (0.722) | -0.589 (0.697) | -0.267 (0.931) | -0.616 (0.666) | -0.055 (0.997) |
| **DNA/Zisupton** | -0.505 (0.747) | -0.276 (0.497) | -0.642 (0.608) | 0.359 (0.418) | -0.428 (0.814) | -0.57 (0.681) | 1.25 (0.207) | -0.902 (0.37) | -0.358 (0.875) | -0.194 (0.959) | -0.526 (0.723) | 0.545 (0.357) | 1.866 (0.116) | -0.303 (0.903) | -0.665 (0.594) | -0.074 (0.994) |
| **DNA/hAT-Ac** | 0.329 (0.371) | 0.224 (0.412) | 2.438 (0.041*) | -0.208 (0.655) | 1.606 (0.109) | -0.67 (0.557) | -0.544 (0.655) | -1.057 (0.214) | -0.435 (0.797) | -0.256 (0.932) | -0.628 (0.567) | -0.016 (0.735) | -0.767 (0.464) | -0.371 (0.844) | -0.836 (0.395) | -0.14 (0.98) |
| **DNA/hAT-Tag1** | -0.086 (0.992) | 1.399 (0.138) | -0.126 (0.983) | -0.124 (0.983) | -0.1 (0.989) | -0.109 (0.987) | -0.095 (0.99) | -0.128 (0.981) | -0.084 (0.993) | -0.032 (0.999) | -0.102 (0.989) | -0.124 (0.983) | -0.135 (0.98) | -0.071 (0.995) | -0.108 (0.987) | NA (1) |
| **DNA/hAT-Tip100** | -0.25 (0.923) | 0.186 (0.473) | -0.394 (0.827) | -0.416 (0.812) | 5.023 (0.016*) | -0.322 (0.876) | -0.277 (0.913) | -0.558 (0.711) | -0.229 (0.941) | -0.095 (0.99) | -0.315 (0.888) | -0.394 (0.836) | -0.352 (0.869) | -0.178 (0.966) | -0.384 (0.833) | NA (1) |
| **LINE/I** | -0.146 (0.979) | -0.459 (0.787) | -0.164 (0.973) | -0.128 (0.983) | -0.102 (0.989) | -0.111 (0.987) | -0.09 (0.992) | -0.18 (0.967) | -0.071 (0.995) | -0.045 (0.998) | -0.118 (0.985) | -0.12 (0.983) | -0.124 (0.984) | -0.063 (0.996) | -0.135 (0.978) | NA (1) |
| **LINE/L1** | -0.911 (0.335) | -0.151 (0.503) | -0.87 (0.28) | 0.358 (0.396) | -0.158 (0.676) | -1.021 (0.254) | -0.149 (0.693) | 0.826 (0.221) | -0.686 (0.569) | -0.321 (0.885) | 0.07 (0.473) | 1.287 (0.132) | -1.081 (0.204) | -0.6 (0.645) | -1.15 (0.144) | -0.15 (0.978) |
| **LINE/L1-Tx1** | -0.067 (0.995) | -0.393 (0.851) | -0.09 (0.992) | -0.158 (0.974) | -0.071 (0.995) | -0.095 (0.991) | -0.045 (0.998) | -0.175 (0.966) | -0.078 (0.994) | -0.032 (0.999) | -0.084 (0.993) | -0.128 (0.984) | -0.126 (0.983) | -0.055 (0.997) | -0.129 (0.981) | NA (1) |
| **LINE/RTE-BovB** | -0.07 (0.948) | -0.082 (0.638) | -0.326 (0.876) | -0.137 (0.882) | -0.202 (0.954) | -0.25 (0.926) | -0.077 (0.942) | -0.079 (0.821) | -0.155 (0.975) | -0.08 (0.993) | -0.229 (0.94) | -0.267 (0.918) | -0.285 (0.915) | -0.128 (0.981) | -0.355 (0.872) | -0.032 (0.999) |
| **LTR** | -0.6 (0.689) | -1.356 (0.112) | -0.823 (0.488) | -0.825 (0.471) | 2.633 (0.064.) | -0.693 (0.597) | 1.024 (0.263) | -1.074 (0.284) | -0.399 (0.848) | -0.19 (0.965) | 0.699 (0.353) | 0.403 (0.448) | 1.601 (0.136) | -0.352 (0.88) | -0.77 (0.507) | -0.09 (0.992) |
| **LTR/Copia** | -0.949 (0.306) | 0.302 (0.355) | -0.798 (0.309) | -0.522 (0.447) | -0.205 (0.656) | -1.07 (0.217) | 1.005 (0.193) | 0.776 (0.217) | -0.644 (0.554) | 1.365 (0.18) | -0.461 (0.527) | -0.236 (0.59) | 0.271 (0.37) | -0.53 (0.63) | -0.049 (0.635) | -0.29 (0.92) |
| **LTR/ERV1** | -0.141 (0.975) | -0.615 (0.636) | -0.209 (0.945) | -0.211 (0.95) | -0.134 (0.979) | -0.204 (0.957) | -0.162 (0.973) | -0.26 (0.917) | -0.1 (0.99) | -0.071 (0.995) | -0.139 (0.971) | -0.209 (0.95) | -0.2 (0.956) | -0.128 (0.983) | -0.205 (0.954) | NA (1) |
| **LTR/ERVK** | -0.051 (0.996) | -0.097 (0.966) | -0.067 (0.995) | -0.06 (0.996) | -0.055 (0.997) | -0.032 (0.999) | -0.032 (0.999) | -0.065 (0.991) | -0.032 (0.999) | NA (1) | -0.055 (0.997) | -0.049 (0.996) | -0.042 (0.998) | -0.032 (0.999) | -0.055 (0.997) | NA (1) |
| **LTR/Gypsy** | -0.097 (0.733) | -0.647 (0.315) | 0.747 (0.23) | -0.147 (0.656) | 0.036 (0.516) | 0.364 (0.351) | 3.063 (0.024*) | 3.115 (0.023*) | 0.468 (0.343) | -0.36 (0.864) | -0.799 (0.389) | 0.656 (0.24) | -0.876 (0.309) | -0.488 (0.693) | -0.117 (0.67) | -0.261 (0.936) |
| **Low complexity** | -0.972 (0.365) | 1.049 (0.164) | 1.571 (0.105) | 0.265 (0.452) | 1.018 (0.248) | 0.339 (0.415) | -0.867 (0.442) | -0.763 (0.322) | 0.518 (0.417) | 2.569 (0.1.) | -0.237 (0.622) | 2.152 (0.052.) | -1.117 (0.242) | -0.611 (0.677) | -0.128 (0.611) | -0.084 (0.993) |
| **RC/Helitron** | 1.746 (0.095.) | -0.586 (0.298) | -0.9 (0.286) | 1.955 (0.053.) | 1.123 (0.19) | 0.537 (0.339) | -0.671 (0.549) | -0.12 (0.601) | -0.555 (0.702) | -0.297 (0.907) | -0.787 (0.436) | -0.884 (0.327) | -0.244 (0.637) | 1.135 (0.227) | -1.008 (0.259) | -0.132 (0.982) |
| **SINE/U** | -0.032 (0.999) | -0.194 (0.942) | -0.051 (0.997) | -0.081 (0.992) | NA (1) | -0.055 (0.997) | -0.042 (0.998) | -0.068 (0.994) | -0.045 (0.998) | NA (1) | -0.032 (0.999) | -0.065 (0.995) | -0.052 (0.997) | -0.032 (0.999) | -0.06 (0.996) | NA (1) |
| **Satellite** | -0.773 (0.491) | 2.156 (0.025*) | -0.464 (0.523) | 0.028 (0.513) | -0.721 (0.55) | 0.703 (0.312) | -0.749 (0.525) | 0.863 (0.244) | -0.57 (0.685) | -0.318 (0.896) | -0.055 (0.724) | 0.419 (0.39) | -0.265 (0.614) | -0.499 (0.755) | -0.472 (0.513) | -0.105 (0.989) |
| **Simple repeat** | -0.502 (0.372) | -0.425 (0.345) | 0.802 (0.202) | 0.708 (0.221) | -0.271 (0.512) | 0.415 (0.32) | -0.84 (0.255) | 0.096 (0.445) | -0.026 (0.633) | 1.486 (0.134) | 0.192 (0.427) | 0.491 (0.323) | -0.972 (0.192) | 0.44 (0.369) | -1.067 (0.14) | 5.099 (0.035*) |
| **Unknown** | -1.55 (0.065.) | 0.515 (0.275) | 1.105 (0.131) | 0 (0.52) | 1.246 (0.128) | -0.21 (0.449) | 0.154 (0.454) | 0.324 (0.378) | 0.802 (0.23) | -0.18 (0.597) | 0.704 (0.232) | -0.13 (0.486) | 0.365 (0.377) | -0.698 (0.312) | -1.255 (0.103) | -0.712 (0.633) |

**Supplementary Table 5. Fine-grained enrichment analysis of individual breakpoint regions in the CAD genome.** Values represent the Z-score and the empirical p-value (in parentheses) obtained from 1,000 permutations using *regioneR*. Positive Z-scores indicate enrichment (presence higher than expected), while negative Z-scores indicate depletion of the genomic feature within the breakpoint interval. CDS: Coding Sequence; GC: GC content; bp: base pairs; NA: feature not present in the randomized background for that specific interval. Significance codes: *p* < 0.05 (*), *p* < 0.01 (**), *p* < 0.001 (***), *p* < 0.10 (.).

| **Chromosome** | 1 | 5 | 19 | 22 | 24 | 25 | 25 | 25 | 26 | 29 | 29 | 29 | 29 | 33 | 35 | 38 | 39 | 40 |
| --- | --- | --- | --- | --- | --- | --- | --- | --- | --- | --- | --- | --- | --- | --- | --- | --- | --- | --- |
| **Start bp** | 4,344,437 | 1 | 1 | 1 | 8,400,000 | 4,655,409 | 5,957,224 | 8,400,000 | 8,400,000 | 3,987,191 | 4,942,141 | 6,701,801 | 7,853,518 | 1,610,824 | 5,575,786 | 1 | 4,400,000 | 1 |
| **End bp** | 4,369,916 | 50,000 | 50,000 | 50,000 | 8,448,568 | 4,657,350 | 5,991,829 | 8,443,197 | 8,417,851 | 4,030,509 | 4,953,518 | 6,735,025 | 7,862,941 | 1,647,489 | 5,586,772 | 50,000 | 4,441,323 | 50,000 |
| **CDS** | -0.527 (0.417) | -1.215 (0.085.) | -1.207 (0.082.) | 0.146 (0.406) | -1.282 (0.066.) | -0.665 (0.35) | -0.649 (0.349) | -0.238 (0.49) | -1.112 (0.086.) | -0.686 (0.296) | -0.43 (0.472) | -1.245 (0.087.) | -0.43 (0.533) | -1.133 (0.088.) | -0.663 (0.388) | 0.233 (0.353) | -1.225 (0.078.) | -1.016 (0.14) |
| **GC** | 0.728 (0.195) | 13.519 (0.001***) | 4.948 (0.003**) | 0.032 (0.435) | 2.064 (0.025*) | 0.433 (0.287) | -0.918 (0.167) | -0.017 (0.543) | 5.575 (0.001***) | 0.203 (0.363) | -0.124 (0.491) | 1.029 (0.133) | 0.221 (0.367) | 4.289 (0.004**) | -0.459 (0.344) | 2.755 (0.011*) | 2.281 (0.018*) | -1.414 (0.05*) |
| **All_Repeats** | -0.9 (0.186) | -3.193 (0.024*) | -3.035 (0.018*) | -0.167 (0.369) | -3.029 (0.023*) | -0.374 (0.374) | -0.022 (0.504) | -0.802 (0.186) | -3.141 (0.027*) | -0.055 (0.472) | -1.113 (0.158) | 0.153 (0.449) | 1.398 (0.101) | -0.32 (0.363) | 2.259 (0.015*) | -0.642 (0.212) | -2.415 (0.023*) | -0.824 (0.185) |
| **DNA** | -0.394 (0.845) | -0.498 (0.766) | -0.534 (0.732) | -0.753 (0.538) | -0.518 (0.745) | -0.322 (0.894) | -0.414 (0.826) | 4.722 (0.003**) | -0.504 (0.743) | 3.196 (0.027*) | -0.213 (0.941) | 0.9 (0.273) | -0.238 (0.94) | -0.527 (0.727) | 2.706 (0.08.) | -0.456 (0.754) | -0.508 (0.727) | -0.553 (0.718) |
| **DNA/CMC-EnSpm** | -0.598 (0.606) | 0.788 (0.183) | 0.266 (0.324) | -1.057 (0.159) | -0.154 (0.707) | 0.534 (0.293) | -0.518 (0.637) | -0.172 (0.704) | -0.739 (0.423) | -0.255 (0.637) | -0.361 (0.822) | -0.246 (0.656) | -0.365 (0.823) | -0.753 (0.396) | -0.46 (0.748) | -0.678 (0.432) | -0.712 (0.433) | -0.669 (0.414) |
| **DNA/hAT-Ac** | 0.311 (0.387) | -0.14 (0.683) | -0.751 (0.452) | -1.099 (0.19) | -0.79 (0.444) | -0.467 (0.738) | 2.428 (0.045*) | -0.732 (0.448) | -0.787 (0.42) | 0.898 (0.211) | 0.92 (0.193) | -0.817 (0.405) | -0.376 (0.839) | -0.188 (0.66) | -0.463 (0.763) | -0.798 (0.436) | -0.795 (0.428) | -0.746 (0.435) |
| **DNA/hAT-Tag1** | -0.06 (0.996) | -0.09 (0.991) | -0.085 (0.991) | -0.148 (0.977) | -0.085 (0.992) | -0.06 (0.996) | -0.045 (0.998) | -0.128 (0.982) | -0.1 (0.989) | -0.096 (0.99) | -0.078 (0.994) | -0.107 (0.988) | NA (1) | -0.124 (0.983) | -0.074 (0.994) | -0.076 (0.992) | -0.104 (0.988) | -0.132 (0.981) |
| **DNA/hAT-Tip100** | -0.311 (0.898) | -0.382 (0.846) | -0.362 (0.845) | -0.568 (0.678) | -0.405 (0.823) | -0.25 (0.932) | -0.287 (0.9) | -0.388 (0.823) | -0.408 (0.819) | -0.436 (0.798) | -0.169 (0.969) | -0.414 (0.824) | -0.179 (0.964) | -0.412 (0.816) | -0.188 (0.958) | -0.381 (0.839) | -0.376 (0.839) | -0.391 (0.826) |
| **DNA/MULE-MuDR** | -0.79 (0.418) | -0.994 (0.252) | -1.004 (0.259) | 0.076 (0.473) | 0.608 (0.271) | -0.654 (0.549) | -0.181 (0.673) | -1.056 (0.224) | -1.001 (0.247) | 0.762 (0.245) | 1.227 (0.143) | 0.781 (0.228) | 0.488 (0.285) | 0.88 (0.217) | -0.598 (0.606) | -0.963 (0.271) | -1.044 (0.234) | -1.054 (0.228) |
| **DNA/PIF-Harbinger** | -0.156 (0.971) | -0.267 (0.919) | -0.286 (0.909) | -0.436 (0.795) | -0.277 (0.915) | -0.168 (0.967) | -0.227 (0.945) | -0.26 (0.925) | -0.29 (0.902) | 2.336 (0.108) | -0.135 (0.98) | -0.29 (0.905) | -0.131 (0.981) | -0.294 (0.902) | 8.316 (0.008**) | -0.292 (0.904) | -0.269 (0.917) | -0.277 (0.916) |
| **DNA/TcMar** | -0.414 (0.808) | -0.588 (0.684) | -0.583 (0.675) | -0.876 (0.427) | 0.713 (0.339) | -0.364 (0.861) | 1.506 (0.189) | -0.588 (0.681) | -0.585 (0.685) | -0.683 (0.614) | 2.753 (0.085.) | -0.647 (0.632) | 2.877 (0.086.) | -0.634 (0.641) | -0.314 (0.898) | -0.666 (0.627) | -0.609 (0.666) | -0.585 (0.694) |
| **DNA/Zisupton** | -0.488 (0.757) | -0.609 (0.636) | -0.626 (0.614) | 1.103 (0.198) | -0.703 (0.569) | -0.416 (0.821) | -0.448 (0.792) | -0.613 (0.631) | -0.696 (0.584) | 0.317 (0.41) | -0.324 (0.886) | 0.383 (0.398) | -0.328 (0.888) | 0.342 (0.421) | -0.374 (0.857) | -0.665 (0.599) | -0.676 (0.582) | -0.635 (0.621) |
| **LINE/I** | -0.085 (0.991) | -0.144 (0.977) | -0.133 (0.981) | -0.185 (0.96) | -0.128 (0.981) | -0.105 (0.989) | -0.124 (0.982) | -0.135 (0.979) | -0.117 (0.985) | -0.145 (0.976) | -0.085 (0.992) | -0.163 (0.969) | -0.062 (0.995) | -0.152 (0.974) | -0.09 (0.991) | -0.144 (0.974) | -0.141 (0.979) | -0.132 (0.98) |
| **LINE/L1** | -0.263 (0.634) | -1.113 (0.172) | -1.19 (0.157) | -0.758 (0.272) | 3.374 (0.008**) | 0.004 (0.497) | -0.814 (0.376) | -1.137 (0.176) | -1.117 (0.189) | -1.174 (0.161) | 0.389 (0.38) | 0.678 (0.261) | -0.556 (0.664) | 0.296 (0.37) | -0.627 (0.585) | -1.11 (0.188) | -1.147 (0.182) | -1.161 (0.167) |
| **LINE/L1-Tx1** | -0.095 (0.991) | -0.116 (0.986) | -0.11 (0.988) | -0.157 (0.975) | -0.14 (0.98) | -0.071 (0.995) | -0.107 (0.988) | -0.11 (0.988) | -0.116 (0.986) | -0.155 (0.975) | -0.055 (0.997) | -0.115 (0.987) | -0.032 (0.999) | -0.111 (0.987) | -0.052 (0.997) | -0.11 (0.988) | -0.118 (0.985) | -0.128 (0.983) |
| **LINE/RTE-BovB** | -0.22 (0.948) | -0.281 (0.912) | -0.244 (0.911) | 2.41 (0.058.) | -0.295 (0.903) | -0.2 (0.955) | -0.202 (0.954) | -0.333 (0.884) | -0.275 (0.913) | -0.341 (0.877) | 6.515 (0.021*) | -0.293 (0.906) | -0.147 (0.978) | -0.28 (0.907) | -0.128 (0.979) | -0.309 (0.897) | -0.305 (0.901) | -0.309 (0.895) |
| **Low_complexity** | 0.783 (0.309) | -0.596 (0.428) | -0.626 (0.423) | 2.112 (0.037*) | -0.114 (0.607) | 0.169 (0.499) | -0.896 (0.422) | -1.158 (0.23) | -1.175 (0.213) | -0.77 (0.359) | 0.686 (0.366) | 0.351 (0.423) | -0.568 (0.705) | 1.427 (0.115) | -0.74 (0.551) | -1.22 (0.191) | -1.175 (0.222) | -1.167 (0.208) |
| **LTR** | -0.599 (0.694) | -0.749 (0.525) | -0.793 (0.502) | 0.934 (0.247) | -0.813 (0.502) | -0.506 (0.77) | -0.595 (0.678) | -0.758 (0.531) | -0.797 (0.506) | 1.029 (0.226) | -0.389 (0.842) | -0.837 (0.468) | -0.322 (0.896) | -0.79 (0.484) | -0.422 (0.819) | -0.766 (0.519) | -0.763 (0.522) | -0.773 (0.509) |
| **LTR/Copia** | -0.867 (0.358) | -0.729 (0.35) | -1.12 (0.168) | -1.093 (0.145) | -0.747 (0.352) | 0.567 (0.268) | -0.866 (0.343) | -1.12 (0.168) | -1.083 (0.174) | 4.637 (0.003**) | -0.654 (0.585) | 1.022 (0.179) | -0.594 (0.649) | -0.095 (0.608) | -0.724 (0.504) | -1.115 (0.183) | -1.18 (0.154) | -1.106 (0.18) |
| **LTR/ERV1** | -0.182 (0.962) | -0.225 (0.945) | -0.192 (0.959) | -0.293 (0.905) | -0.158 (0.965) | -0.139 (0.981) | -0.177 (0.965) | 9.406 (0.003**) | -0.171 (0.966) | -0.209 (0.95) | -0.11 (0.988) | -0.227 (0.94) | -0.102 (0.989) | -0.22 (0.947) | -0.121 (0.984) | -0.204 (0.953) | -0.197 (0.955) | -0.202 (0.955) |
| **LTR/ERVK** | -0.032 (0.999) | -0.067 (0.995) | -0.063 (0.996) | -0.102 (0.989) | -0.041 (0.997) | -0.032 (0.999) | NA (1) | -0.083 (0.991) | -0.042 (0.996) | -0.09 (0.991) | -0.045 (0.998) | -0.06 (0.996) | NA (1) | -0.034 (0.998) | -0.04 (0.998) | -0.067 (0.995) | -0.05 (0.997) | -0.043 (0.997) |
| **LTR/Gypsy** | -0.026 (0.755) | 5.038 (0.002**) | 1.907 (0.07.) | -0.669 (0.337) | -0.901 (0.276) | 0.207 (0.407) | -0.058 (0.737) | 2.163 (0.044*) | -0.935 (0.269) | 2.247 (0.048*) | 0.542 (0.311) | 0.395 (0.327) | 0.709 (0.273) | -0.948 (0.246) | -0.538 (0.648) | -1 (0.252) | -0.957 (0.248) | -0.876 (0.261) |
| **RC/Helitron** | -0.712 (0.533) | -0.928 (0.315) | -0.927 (0.292) | 0.349 (0.336) | -0.397 (0.538) | 0.43 (0.371) | -0.716 (0.508) | -0.967 (0.294) | -0.917 (0.301) | 1.816 (0.065.) | 0.73 (0.288) | -0.976 (0.279) | -0.484 (0.762) | 0.633 (0.268) | -0.612 (0.648) | -0.921 (0.295) | -0.911 (0.282) | -0.87 (0.309) |
| **Satellite** | 0.148 (0.494) | 0.245 (0.459) | -1.056 (0.28) | 0.124 (0.467) | -0.396 (0.548) | 1.476 (0.161) | 1.077 (0.221) | -1.012 (0.307) | -1.048 (0.275) | -0.526 (0.472) | -0.555 (0.696) | 0.701 (0.296) | -0.483 (0.77) | -1.077 (0.238) | -0.566 (0.688) | -1.064 (0.258) | -0.978 (0.305) | -1.025 (0.276) |
| **Simple_repeat** | -0.202 (0.504) | -0.675 (0.268) | -1.458 (0.081.) | 0.102 (0.483) | -0.039 (0.536) | -0.619 (0.377) | 1.095 (0.164) | -0.971 (0.162) | -1.774 (0.061.) | -0.16 (0.473) | 1.5 (0.105) | -0.683 (0.244) | -0.609 (0.422) | 0.06 (0.505) | 1.424 (0.111) | -1.751 (0.065.) | -1.879 (0.052.) | -1.724 (0.065.) |
| **SINE/U** | -0.074 (0.994) | -0.078 (0.994) | -0.045 (0.998) | -0.123 (0.981) | -0.055 (0.996) | -0.055 (0.997) | -0.063 (0.996) | -0.06 (0.996) | -0.085 (0.992) | -0.08 (0.993) | -0.032 (0.999) | -0.055 (0.996) | -0.045 (0.998) | -0.06 (0.996) | -0.032 (0.999) | -0.06 (0.996) | -0.08 (0.993) | -0.055 (0.997) |
| **Unknown** | -0.286 (0.391) | -1.857 (0.065.) | -1.478 (0.087.) | 0.357 (0.363) | -0.651 (0.218) | -0.22 (0.442) | 0.865 (0.203) | -1.529 (0.085.) | -2.417 (0.051.) | -0.767 (0.192) | 2.061 (0.032*) | 0.006 (0.547) | -0.806 (0.276) | 0.002 (0.533) | 0.376 (0.371) | -2.257 (0.052.) | -2.329 (0.058.) | -2.354 (0.053.) |

**Supplementary Table 6. Microsatellite genotypes for all individuals included in the crosses.** Genotypes at 4 microsatellite loci (WE1, 468, EG7 and XAF) for the parental individuals, F₁ hybrids, and F₂ progeny used in the analyses. Columns correspond to the alleles detected at each locus, and rows indicate the individuals assigned to each generation and cross. “1” indicates presence of an allele and “–” indicates absence.

|  |  |  | **WE1** | | **468** | | **FG7** | | | | | | **XAF** | |
| --- | --- | --- | --- | --- | --- | --- | --- | --- | --- | --- | --- | --- | --- | --- |
| **Cross** | **Generation** | **Individual** | **224** | **229** | **273** | **276** | **393** | **399** | **406** | **409** | **414** | **461** | **390** | **407** |
| - | Parent | CAD | 1 | - | 1 | - | 1 | - | - | 1 | - | - | - | 1 |
| - | Parent | ALF | - | 1 | 1 | - | 1 | - | - | 1 | - | - | - | 1 |
| - | Parent | HUE | 1 | - | - | 1 | 1 | - | 1 | - | - | 1 | 1 | 1 |
| CAD x ALF | F1 | F1.01 | 1 | 1 | 1 | - | 1 | - | - | 1 | - | - | - | 1 |
| CAD x ALF | F2 | CADxALF-F2.01 | 1 | - | 1 | - | - | - | - | - | - | - | - | 1 |
| CAD x ALF | F2 | CADxALF-F2.02 | 1 | - | 1 | - | 1 | - | - | 1 | - | - | - | 1 |
| CAD x ALF | F2 | CADxALF-F2.03 | 1 | 1 | 1 | - | 1 | - | - | 1 | - | - | - | 1 |
| CAD x ALF | F2 | CADxALF-F2.04 | 1 | - | 1 | - | 1 | - | - | 1 | - | - | - | 1 |
| CAD x ALF | F2 | CADxALF-F2.05 | 1 | 1 | 1 | - | 1 | - | - | 1 | - | - | - | 1 |
| CAD x ALF | F2 | CADxALF-F2.06 | 1 | - | 1 | - | 1 | - | - | 1 | - | - | - | 1 |
| CAD x ALF | F2 | CADxALF-F2.07 | 1 | 1 | 1 | - | 1 | - | - | 1 | - | - | - | 1 |
| CAD x ALF | F2 | CADxALF-F2.08 | 1 | 1 | 1 | - | 1 | - | - | 1 | - | - | - | 1 |
| CAD x ALF | F2 | CADxALF-F2.09 | 1 | 1 | 1 | - | 1 | - | - | - | - | - | - | 1 |
| CAD x ALF | F2 | CADxALF-F2.10 | 1 | - | 1 | - | 1 | - | - | - | - | - | - | 1 |
| CAD x ALF | F2 | CADxALF-F2.11 | 1 | 1 | 1 | - | 1 | - | - | - | - | - | - | 1 |
| CAD x ALF | F2 | CADxALF-F2.12 | - | 1 | 1 | - | 1 | - | - | 1 | - | - | - | 1 |
| CAD x ALF | F2 | CADxALF-F2.13 | 1 | - | 1 | - | 1 | - | - | - | - | - | - | 1 |
| CAD x ALF | F2 | CADxALF-F2.14 | 1 | - | 1 | - | 1 | - | - | - | - | - | - | 1 |
| CAD x ALF | F2 | CADxALF-F2.15 | - | 1 | 1 | - | 1 | - | - | 1 | - | - | - | 1 |
| CAD x ALF | F2 | CADxALF-F2.16 | - | 1 | 1 | - | 1 | - | - | 1 | - | - | - | 1 |
| CAD x ALF | F2 | CADxALF-F2.17 | - | 1 | 1 | - | 1 | - | - | 1 | - | - | - | 1 |
| CAD x ALF | F2 | CADxALF-F2.18 | 1 | 1 | 1 | - | 1 | - | - | - | - | - | - | 1 |
| CAD x ALF | F2 | CADxALF-F2.19 | 1 | - | 1 | - | 1 | - | - | - | - | - | - | 1 |
| CAD x ALF | F2 | CADxALF-F2.20 | 1 | 1 | 1 | - | 1 | - | - | 1 | - | - | - | 1 |
| CAD x ALF | F2 | CADxALF-F2.21 | 1 | 1 | 1 | - | 1 | - | - | - | - | - | - | 1 |
| CAD x ALF | F2 | CADxALF-F2.22 | 1 | - | 1 | - | 1 | - | - | - | - | - | - | 1 |
| CAD x ALF | F2 | CADxALF-F2.23 | - | 1 | 1 | - | 1 | - | - | 1 | - | - | - | 1 |
| CAD x ALF | F2 | CADxALF-F2.24 | 1 | - | 1 | - | 1 | - | - | 1 | - | - | - | 1 |
| CAD x ALF | F2 | CADxALF-F2.25 | 1 | - | 1 | - | 1 | - | - | 1 | - | - | - | 1 |
| CAD x ALF | F2 | CADxALF-F2.26 | - | 1 | 1 | - | 1 | - | - | 1 | - | - | - | 1 |
| CAD x ALF | F2 | CADxALF-F2.27 | - | 1 | 1 | - | 1 | - | - | - | - | - | - | 1 |
| CAD x ALF | F2 | CADxALF-F2.28 | 1 | - | 1 | - | 1 | - | - | 1 | - | - | - | 1 |
| CAD x ALF | F2 | CADxALF-F2.29 | - | 1 | 1 | - | 1 | - | - | 1 | - | - | - | 1 |
| CAD x ALF | F2 | CADxALF-F2.30 | - | 1 | 1 | - | 1 | - | - | 1 | - | - | - | 1 |
| CAD x ALF | F2 | CADxALF-F2.31 | 1 | 1 | 1 | - | 1 | - | - | - | - | - | - | 1 |
| CAD x ALF | F2 | CADxALF-F2.32 | 1 | - | 1 | - | 1 | - | - | 1 | - | - | - | 1 |
| CAD x ALF | F2 | CADxALF-F2.33 | 1 | - | 1 | - | 1 | - | - | 1 | - | - | - | 1 |
| CAD x ALF | F2 | CADxALF-F2.34 | - | 1 | 1 | - | 1 | - | - | 1 | - | - | - | 1 |
| CAD x ALF | F2 | CADxALF-F2.35 | - | 1 | 1 | - | 1 | - | - | 1 | - | - | - | 1 |
| CAD x ALF | F2 | CADxALF-F2.36 | 1 | 1 | 1 | - | 1 | - | - | - | - | - | - | 1 |
| CAD x ALF | F2 | CADxALF-F2.37 | - | 1 | 1 | - | 1 | - | - | - | - | - | - | 1 |
| CAD x ALF | F2 | CADxALF-F2.38 | - | 1 | 1 | - | 1 | - | - | 1 | - | - | - | 1 |
| CAD x ALF | F2 | CADxALF-F2.39 | - | 1 | 1 | - | 1 | - | - | 1 | - | - | - | 1 |
| CAD x ALF | F2 | CADxALF-F2.40 | 1 | - | 1 | - | 1 | - | - | 1 | - | - | - | 1 |
| CAD x ALF | F2 | CADxALF-F2.41 | 1 | 1 | 1 | - | 1 | - | - | 1 | - | - | - | 1 |
| CAD x ALF | F2 | CADxALF-F2.42 | 1 | 1 | 1 | - | 1 | - | - | 1 | - | - | - | 1 |
| CAD x ALF | F2 | CADxALF-F2.43 | 1 | 1 | 1 | - | 1 | - | - | 1 | - | - | - | 1 |
| CAD x ALF | F2 | CADxALF-F2.44 | - | - | 1 | - | 1 | - | - | 1 | - | - | - | 1 |
| CAD x ALF | F2 | CADxALF-F2.45 | 1 | 1 | 1 | - | 1 | - | - | 1 | - | - | - | 1 |
| CAD x ALF | F2 | CADxALF-F2.46 | - | 1 | 1 | - | 1 | - | - | 1 | - | - | - | 1 |
| CAD x ALF | F2 | CADxALF-F2.47 | 1 | 1 | 1 | - | 1 | - | - | 1 | - | - | - | 1 |
| CAD x ALF | F2 | CADxALF-F2.48 | - | 1 | 1 | - | 1 | - | - | 1 | - | - | - | 1 |
| CAD x ALF | F2 | CADxALF-F2.49 | 1 | 1 | 1 | - | 1 | - | - | 1 | - | - | - | 1 |
| CAD x ALF | F2 | CADxALF-F2.50 | 1 | 1 | 1 | - | 1 | - | - | 1 | - | - | - | 1 |
| CAD x ALF | F2 | CADxALF-F2.51 | - | 1 | 1 | - | 1 | - | - | 1 | - | - | - | 1 |
| CAD x ALF | F2 | CADxALF-F2.52 | 1 | 1 | 1 | - | 1 | - | - | 1 | - | - | - | 1 |
| CAD x ALF | F2 | CADxALF-F2.53 | 1 | 1 | 1 | - | 1 | - | - | 1 | - | - | - | 1 |
| CAD x ALF | F2 | CADxALF-F2.54 | 1 | - | 1 | - | 1 | - | - | 1 | - | - | - | 1 |
| CAD x ALF | F2 | CADxALF-F2.55 | 1 | 1 | 1 | - | 1 | - | - | - | - | - | - | 1 |
| CAD x HUE | F1 | CADxHUE-F1.001 | 1 | 1 | 1 | 1 | 1 | - | 1 | 1 | - | 1 | 1 | 1 |
| CAD x HUE | F1 | CADxHUE-F1.002 | 1 | 1 | 1 | 1 | 1 | - | 1 | 1 | - | 1 | 1 | 1 |
| CAD x HUE | F1 | CADxHUE-F1.003 | 1 | 1 | 1 | 1 | 1 | - | 1 | 1 | - | 1 | 1 | 1 |
| CAD x HUE | F1 | CADxHUE-F1.004 | 1 | - | 1 | 1 | - |  | 1 | - | - | - | - | 1 |
| CAD x HUE | F2 | CADxHUE-F2.001 | 1 | - | - | 1 | 1 | - | 1 | - | - | - | - | 1 |
| CAD x HUE | F2 | CADxHUE-F2.002 | 1 | - | - | 1 | 1 | - | - | 1 | - | - | - | - |
| CAD x HUE | F2 | CADxHUE-F2.003 | 1 | - | 1 | - | - | - | - | 1 | - | - | - | 1 |
| CAD x HUE | F2 | CADxHUE-F2.004 | 1 | - | 1 | 1 | - | - | - | 1 | - | 1 | - | 1 |
| CAD x HUE | F2 | CADxHUE-F2.005 | 1 | - | 1 | 1 | - | - | 1 | - | - | 1 | - | 1 |
| CAD x HUE | F2 | CADxHUE-F2.006 | 1 | - | 1 | 1 | - | - | - | 1 | - | 1 | 1 | - |
| CAD x HUE | F2 | CADxHUE-F2.007 | 1 | - | - | 1 | - | - | - | 1 | - | - | 1 | - |
| CAD x HUE | F2 | CADxHUE-F2.008 | 1 | - | 1 | 1 | - | - | - | 1 | - | 1 | 1 | - |
| CAD x HUE | F2 | CADxHUE-F2.009 | 1 | - | 1 | 1 | - | - | - | - | - | 1 | 1 | - |
| CAD x HUE | F2 | CADxHUE-F2.010 | 1 | - | 1 | 1 | - | - | - | - | - | 1 | 1 | - |
| CAD x HUE | F2 | CADxHUE-F2.011 | 1 | - | - | 1 | - | - | - | 1 | - | 1 | 1 | 1 |
| CAD x HUE | F2 | CADxHUE-F2.012 | 1 | - | - | 1 | 1 | - | - | - | - | - | 1 | 1 |
| CAD x HUE | F2 | CADxHUE-F2.013 | 1 | - | 1 | - | - | - | - | 1 | - | 1 | - | 1 |
| CAD x HUE | F2 | CADxHUE-F2.014 | 1 | - | 1 | - | 1 | - | 1 | 1 | - | 1 | 1 | 1 |
| CAD x HUE | F2 | CADxHUE-F2.015 | 1 | - | 1 | 1 | - | - | 1 | - | - | 1 | 1 | 1 |
| CAD x HUE | F2 | CADxHUE-F2.016 | 1 | - | 1 | 1 | - | - | 1 | - | - | 1 | 1 | - |
| CAD x HUE | F2 | CADxHUE-F2.017 | 1 | - | 1 | 1 | - | - | - | - | - | 1 | 1 | - |
| CAD x HUE | F2 | CADxHUE-F2.018 | 1 | - | - | 1 | - | - | - | - | - | 1 | - | 1 |
| CAD x HUE | F2 | CADxHUE-F2.019 | 1 | - | - | 1 | - | - | - | 1 | - | - | 1 | - |
| CAD x HUE | F2 | CADxHUE-F2.020 | 1 | - | 1 | - | - | - | - | 1 | - | 1 | 1 | - |
| CAD x HUE | F2 | CADxHUE-F2.021 | 1 | - | - | 1 | 1 | - | 1 | 1 | - | - | 1 | 1 |
| CAD x HUE | F2 | CADxHUE-F2.022 | 1 | - | 1 | 1 | 1 | - | 1 | 1 | - | 1 | 1 | - |
| CAD x HUE | F2 | CADxHUE-F2.023 | 1 | - | - | 1 | 1 | - | 1 | - | - | 1 | 1 | 1 |
| CAD x HUE | F2 | CADxHUE-F2.024 | 1 | - | 1 | 1 | 1 | - | 1 | - | - | 1 | - | 1 |
| CAD x HUE | F2 | CADxHUE-F2.025 | 1 | - | 1 | 1 | 1 | - | 1 | - | - | 1 | 1 | 1 |
| CAD x HUE | F2 | CADxHUE-F2.026 | 1 | - | - | 1 | 1 | - | 1 | 1 | - | 1 | 1 | 1 |
| CAD x HUE | F2 | CADxHUE-F2.027 | 1 | - | 1 | - | 1 | - | 1 | 1 | - | 1 | - | 1 |
| CAD x HUE | F2 | CADxHUE-F2.028 | 1 | - | 1 | 1 | - | - | - | 1 | - | 1 | 1 | 1 |
| CAD x HUE | F2 | CADxHUE-F2.029 | 1 | - | 1 | 1 | 1 | - | 1 | 1 | - | 1 | 1 | - |
| CAD x HUE | F2 | CADxHUE-F2.030 | 1 | - | - | 1 | 1 | - | - | 1 | - | 1 | 1 | 1 |
| CAD x HUE | F2 | CADxHUE-F2.031 | 1 | - | 1 | - | 1 | - | - | 1 | - | 1 | - | 1 |
| CAD x HUE | F2 | CADxHUE-F2.032 | 1 | - | 1 | 1 | 1 | - | - | 1 | - | 1 | 1 | 1 |
| CAD x HUE | F2 | CADxHUE-F2.033 | 1 | - | 1 | 1 | 1 | - | 1 | 1 | - | 1 | 1 | - |
| CAD x HUE | F2 | CADxHUE-F2.034 | 1 | - | - | 1 | 1 | - | - | 1 | - | 1 | 1 | 1 |
| CAD x HUE | F2 | CADxHUE-F2.035 | 1 | - | 1 | 1 | 1 | - | - | - | - | 1 | 1 | - |
| CAD x HUE | F2 | CADxHUE-F2.036 | 1 | - | - | 1 | 1 | - | 1 | - | - | - | 1 | 1 |
| CAD x HUE | F2 | CADxHUE-F2.037 | 1 | - | - | 1 | - | - | - | - | - | 1 | - | 1 |
| CAD x HUE | F2 | CADxHUE-F2.038 | 1 | - | 1 | 1 | 1 | - | - | 1 | - | - | 1 | 1 |
| CAD x HUE | F2 | CADxHUE-F2.039 | 1 | - | 1 | 1 | 1 | - | 1 | 1 | - | 1 | 1 | 1 |
| CAD x HUE | F2 | CADxHUE-F2.040 | 1 | - | - | 1 | - | - | - | 1 | - | 1 | 1 | 1 |
| CAD x HUE | F2 | CADxHUE-F2.041 | 1 | - | 1 | 1 | 1 | - | - | 1 | - | - | - | 1 |
| CAD x HUE | F2 | CADxHUE-F2.042 | 1 | - | 1 | 1 | 1 | - | - | 1 | - | 1 | 1 | 1 |
| CAD x HUE | F2 | CADxHUE-F2.043 | 1 | - | 1 | 1 | 1 | - | - | 1 | - | 1 | 1 | - |
| CAD x HUE | F2 | CADxHUE-F2.044 | 1 | - | 1 | 1 | 1 | - | - | 1 | - | 1 | 1 | 1 |
| CAD x HUE | F2 | CADxHUE-F2.045 | 1 | - | 1 | 1 | 1 | - | - | 1 | - | 1 | 1 | - |
| CAD x HUE | F2 | CADxHUE-F2.046 | 1 | - | 1 | 1 | 1 | - | 1 | 1 | - | - | - | 1 |
| CAD x HUE | F2 | CADxHUE-F2.047 | 1 | - | - | 1 | - | - | - | 1 | - | - | 1 | - |
| CAD x HUE | F2 | CADxHUE-F2.048 | 1 | - | 1 | 1 | 1 | - | 1 | 1 | - | 1 | 1 | 1 |
| CAD x HUE | F2 | CADxHUE-F2.049 | 1 | - | 1 | 1 | 1 | - | - | 1 | - | - | 1 | 1 |
| CAD x HUE | F2 | CADxHUE-F2.050 | 1 | - | 1 | 1 | 1 | - | - | 1 | - | 1 | 1 | 1 |
| CAD x HUE | F2 | CADxHUE-F2.051 | 1 | - | 1 | 1 | 1 | - | - | 1 | - | 1 | 1 | 1 |
| CAD x HUE | F2 | CADxHUE-F2.052 | 1 | - | 1 | 1 | 1 | - | 1 | 1 | - | 1 | 1 | 1 |
| CAD x HUE | F2 | CADxHUE-F2.053 | 1 | - | 1 | 1 | 1 | - | 1 | - | - | - | 1 | - |
| CAD x HUE | F2 | CADxHUE-F2.054 | 1 | - | 1 | 1 | 1 | - | - | - | - | 1 | 1 | 1 |
| CAD x HUE | F2 | CADxHUE-F2.055 | 1 | - | - | 1 | 1 | - | - | - | - | 1 | 1 | 1 |
| CAD x HUE | F2 | CADxHUE-F2.056 | 1 | - | 1 | 1 | 1 | - | - | 1 | - | 1 | 1 | 1 |
| CAD x HUE | F2 | CADxHUE-F2.057 | 1 | - | 1 | - | 1 | - | - | 1 | - | 1 | - | 1 |
| CAD x HUE | F2 | CADxHUE-F2.058 | 1 | - | 1 | 1 | 1 | - | - | 1 | - | - | - | 1 |
| CAD x HUE | F2 | CADxHUE-F2.059 | 1 | - | 1 | 1 | 1 | - | - | 1 | - | - | - | 1 |
| CAD x HUE | F2 | CADxHUE-F2.060 | 1 | - | 1 | 1 | 1 | - | - | 1 | - | 1 | 1 | - |
| CAD x HUE | F2 | CADxHUE-F2.061 | 1 | - | 1 | - | 1 | - | 1 | 1 | - | - | - | 1 |
| CAD x HUE | F2 | CADxHUE-F2.062 | 1 | - | - | 1 | 1 | - | - | - | - | 1 | 1 | - |
| CAD x HUE | F2 | CADxHUE-F2.063 | 1 | - | 1 | 1 | 1 | - | - | 1 | - | - | 1 | 1 |
| CAD x HUE | F2 | CADxHUE-F2.064 | 1 | - | 1 | 1 | 1 | - | - | 1 | - | - | 1 | - |
| CAD x HUE | F2 | CADxHUE-F2.065 | 1 | - | 1 | 1 | 1 | - | 1 | 1 | - | - | 1 | 1 |
| CAD x HUE | F2 | CADxHUE-F2.066 | 1 | - | 1 | 1 | 1 | - | - | - | - | 1 | 1 | - |
| CAD x HUE | F2 | CADxHUE-F2.067 | 1 | - | 1 | 1 | 1 | - | - | 1 | - | 1 | 1 | 1 |
| CAD x HUE | F2 | CADxHUE-F2.068 | 1 | - | 1 | 1 | - | - | - | 1 | - | 1 | - | 1 |
| CAD x HUE | F2 | CADxHUE-F2.069 | 1 | - | 1 | - | 1 | - | - | - | - | 1 | 1 | - |
| CAD x HUE | F2 | CADxHUE-F2.070 | 1 | - | 1 | 1 | 1 | - | 1 | - | - | 1 | 1 | 1 |
| CAD x HUE | F2 | CADxHUE-F2.071 | 1 | - | 1 | 1 | 1 | - | - | 1 | - | 1 | 1 | 1 |
| CAD x HUE | F2 | CADxHUE-F2.072 | 1 | - | 1 | 1 | 1 | - | - | 1 | - | - | 1 | 1 |
| CAD x HUE | F2 | CADxHUE-F2.073 | 1 | - | 1 | 1 | 1 | - | - | 1 | - | - | 1 | 1 |
| CAD x HUE | F2 | CADxHUE-F2.074 | 1 | - | 1 | 1 | 1 | - | - | - | - | 1 | 1 | 1 |
| CAD x HUE | F2 | CADxHUE-F2.075 | 1 | - | 1 | 1 | 1 | - | 1 | 1 | - | - | 1 | 1 |
| CAD x HUE | F2 | CADxHUE-F2.076 | 1 | - | 1 | 1 | 1 | - | 1 | 1 | - | 1 | 1 | 1 |
| CAD x HUE | F2 | CADxHUE-F2.077 | 1 | - | 1 | 1 | - | - | - | 1 | - | 1 | 1 | - |
| CAD x HUE | F2 | CADxHUE-F2.078 | 1 | - | 1 | 1 | 1 | - | - | 1 | - | - | 1 | 1 |
| CAD x HUE | F2 | CADxHUE-F2.079 | 1 | - | 1 | 1 | 1 | - | - | 1 | - | 1 | 1 | 1 |
| CAD x HUE | F2 | CADxHUE-F2.080 | 1 | - | 1 | 1 | 1 | - | - | 1 | - | - | 1 | 1 |
| CAD x HUE | F2 | CADxHUE-F2.081 | 1 | - | 1 | 1 | 1 | - | - | 1 | - | 1 | 1 | 1 |
| CAD x HUE | F2 | CADxHUE-F2.082 | 1 | - | 1 | 1 | 1 | - | - | 1 | - | 1 | 1 | - |
| CAD x HUE | F2 | CADxHUE-F2.083 | 1 | - | 1 | - | 1 | - | 1 | - | - | - | 1 | - |
| CAD x HUE | F2 | CADxHUE-F2.084 | 1 | - | 1 | 1 | 1 | - | 1 | 1 | - | - | 1 | - |
| CAD x HUE | F2 | CADxHUE-F2.085 | 1 | - | 1 | 1 | 1 | - | - | 1 | - | - | 1 | - |
| CAD x HUE | F2 | CADxHUE-F2.086 | 1 | - | 1 | 1 | 1 | - | 1 | 1 | - | 1 | 1 | - |
| CAD x HUE | F2 | CADxHUE-F2.087 | 1 | - | 1 | 1 | 1 | - | - | 1 | - | 1 | 1 | 1 |
| CAD x HUE | F2 | CADxHUE-F2.088 | 1 | - | 1 | 1 | 1 | - | - | 1 | - | 1 | 1 | - |
| CAD x HUE | F2 | CADxHUE-F2.089 | 1 | - | 1 | 1 | 1 | - | - | 1 | - | 1 | 1 | - |
| CAD x HUE | F2 | CADxHUE-F2.090 | 1 | - | 1 | - | 1 | - | - | 1 | - | 1 | 1 | 1 |
| CAD x HUE | F2 | CADxHUE-F2.091 | 1 | - | 1 | 1 | 1 | - | - | - | - | 1 | 1 | 1 |
| CAD x HUE | F2 | CADxHUE-F2.092 | 1 | - | - | 1 | 1 | - | - | 1 | - | 1 | 1 | 1 |
| CAD x HUE | F2 | CADxHUE-F2.093 | 1 | - | 1 | 1 | 1 | - | - | 1 | - | - | 1 | - |
| CAD x HUE | F2 | CADxHUE-F2.094 | 1 | - | 1 | 1 | 1 | - | - | - | - | 1 | 1 | 1 |
| CAD x HUE | F2 | CADxHUE-F2.095 | 1 | - | 1 | 1 | 1 | - | - | 1 | - | - | 1 | 1 |
| CAD x HUE | F2 | CADxHUE-F2.096 | 1 | - | 1 | 1 | 1 | - | - | - | - | 1 | 1 | 1 |
| CAD x HUE | F2 | CADxHUE-F2.097 | 1 | - | 1 | - | 1 | - | - | - | - | 1 | - | 1 |
| CAD x HUE | F2 | CADxHUE-F2.098 | 1 | - | 1 | 1 | 1 | - | - | - | - | 1 | 1 | - |
| CAD x HUE | F2 | CADxHUE-F2.099 | 1 | - | - | 1 | 1 | - | - | - | - | 1 | 1 | 1 |
| CAD x HUE | F2 | CADxHUE-F2.100 | 1 | - | 1 | 1 | 1 | - | - | 1 | - | 1 | 1 | 1 |
| CAD x HUE | F2 | CADxHUE-F2.101 | 1 | - | 1 | - | 1 | - | - | 1 | - | 1 | 1 | - |
| CAD x HUE | F2 | CADxHUE-F2.102 | 1 | - | 1 | 1 | 1 | - | - | 1 | - | 1 | 1 | 1 |
| CAD x HUE | F2 | CADxHUE-F2.103 | 1 | - | - | 1 | 1 | - | - | 1 | - | - | 1 | - |
| CAD x HUE | F2 | CADxHUE-F2.104 | 1 | - | 1 | 1 | 1 | - | - | 1 | - | - | - | 1 |
| CAD x HUE | F2 | CADxHUE-F2.105 | 1 | - | 1 | 1 | 1 | - | - | 1 | - | 1 | 1 | - |
| CAD x HUE | F2 | CADxHUE-F2.106 | 1 | - | - | 1 | 1 | - | - | 1 | - | - | 1 | - |
| CAD x HUE | F2 | CADxHUE-F2.107 | 1 | - | 1 | 1 | 1 | - | - | 1 | - | 1 | - | 1 |
| CAD x HUE | F2 | CADxHUE-F2.108 | 1 | - | 1 | 1 | 1 | - | - | 1 | - | 1 | 1 | 1 |
| CAD x HUE | F2 | CADxHUE-F2.109 | 1 | - | 1 | 1 | 1 | - | - | 1 | - | 1 | 1 | 1 |
| CAD x HUE | F2 | CADxHUE-F2.110 | 1 | - | 1 | - | - | - | - | 1 | - | - | 1 | 1 |
| CAD x HUE | F2 | CADxHUE-F2.111 | 1 | - | 1 | 1 | - | - | - | 1 | - | 1 | - | 1 |
| CAD x HUE | F2 | CADxHUE-F2.112 | 1 | - | - | 1 | 1 | - | - | 1 | - | 1 | 1 | 1 |
| CAD x HUE | F2 | CADxHUE-F2.113 | 1 | - | 1 | 1 | 1 | - | - | - | - | 1 | 1 | 1 |
| CAD x HUE | F2 | CADxHUE-F2.114 | 1 | - | 1 | - | 1 | - | - | 1 | - | - | - | 1 |
| CAD x HUE | F2 | CADxHUE-F2.115 | 1 | - | 1 | 1 | 1 | - | - | 1 | - | 1 | 1 | - |
| CAD x HUE | F2 | CADxHUE-F2.116 | 1 | - | 1 | 1 | 1 | - | - | 1 | - | - | 1 | 1 |
| CAD x HUE | F2 | CADxHUE-F2.117 | 1 | - | 1 | 1 | - | - | - | - | - | 1 | 1 | 1 |
| CAD x HUE | F2 | CADxHUE-F2.118 | 1 | - | 1 | 1 | 1 | - | - | 1 | - | 1 | 1 | 1 |
| CAD x HUE | F2 | CADxHUE-F2.119 | 1 | - | 1 | 1 | 1 | - | - | 1 | - | 1 | 1 | 1 |
| CAD x HUE | F2 | CADxHUE-F2.120 | 1 | - | 1 | 1 | 1 | - | 1 | - | - | 1 | 1 | 1 |
| CAD x HUE | F2 | CADxHUE-F2.121 | 1 | - | 1 | - | 1 | - | - | 1 | - | 1 | - | 1 |
| CAD x HUE | F2 | CADxHUE-F2.122 | 1 | - | 1 | - | 1 | - | 1 | 1 | - | - | 1 | 1 |
| CAD x HUE | F2 | CADxHUE-F2.123 | 1 | - | 1 | 1 | 1 | - | 1 | - | - | 1 | 1 | - |
| CAD x HUE | F2 | CADxHUE-F2.124 | 1 | - | 1 | 1 | 1 | - | - | - | - | 1 | - | 1 |
| CAD x HUE | F2 | CADxHUE-F2.125 | 1 | - | 1 | 1 | 1 | - | - | - | - | 1 | 1 | 1 |
| CAD x HUE | F2 | CADxHUE-F2.126 | 1 | - | 1 | - | 1 | - | - | - | - | - | 1 | - |
| CAD x HUE | F2 | CADxHUE-F2.127 | 1 | - | 1 | - | 1 | - | - | 1 | - | - | - | 1 |
| CAD x HUE | F2 | CADxHUE-F2.128 | 1 | - | 1 | - | 1 | - | - | 1 | - | - | - | 1 |
| CAD x HUE | F2 | CADxHUE-F2.129 | 1 | - | 1 | 1 | 1 | - | - | - | - | 1 | 1 | - |
| CAD x HUE | F2 | CADxHUE-F2.130 | 1 | - | 1 | - | 1 | - | - | 1 | - | - | 1 | 1 |
| CAD x HUE | F2 | CADxHUE-F2.131 | 1 | - | 1 | - | 1 | - | - | 1 | - | - | - | 1 |
| CAD x HUE | F2 | CADxHUE-F2.132 | 1 | - | 1 | - | 1 | - | - | 1 | - | - | 1 | 1 |
| CAD x HUE | F2 | CADxHUE-F2.133 | 1 | - | 1 | - | 1 | - | 1 | 1 | - | - | - | 1 |
| CAD x HUE | F2 | CADxHUE-F2.134 | 1 | - | 1 | 1 | 1 | - | 1 | 1 | - | 1 | 1 | 1 |
| CAD x HUE | F2 | CADxHUE-F2.135 | 1 | - | 1 | 1 | 1 | - | - | 1 | - | 1 | 1 | 1 |
| CAD x HUE | F2 | CADxHUE-F2.136 | 1 | - | 1 | 1 | 1 | - | 1 | 1 | - | 1 | - | 1 |
| CAD x HUE | F2 | CADxHUE-F2.137 | 1 | - | 1 | - | 1 | - | 1 | 1 | - | 1 | - | 1 |
| CAD x HUE | F2 | CADxHUE-F2.138 | 1 | - | 1 | 1 | 1 | - | 1 | 1 | - | 1 | 1 | 1 |
| CAD x HUE | F2 | CADxHUE-F2.139 | 1 | - | 1 | - | 1 | - | - | 1 | - | - | - | 1 |
| CAD x HUE | F2 | CADxHUE-F2.140 | 1 | - | 1 | - | 1 | - | - | 1 | - | - | 1 | 1 |
| CAD x HUE | F2 | CADxHUE-F2.141 | 1 | - | 1 | - | 1 | - | 1 | - | - | 1 | 1 | 1 |
| CAD x HUE | F2 | CADxHUE-F2.142 | 1 | - | 1 | 1 | 1 | - | - | 1 | - | 1 | 1 | 1 |
| CAD x HUE | F2 | CADxHUE-F2.143 | 1 | - | 1 | 1 | 1 | - | - | 1 | - | 1 | 1 | 1 |
| CAD x HUE | F2 | CADxHUE-F2.144 | 1 | - | 1 | 1 | 1 | - | - | 1 | - | 1 | 1 | 1 |

**Supplementary Table 7. Summary statistics of linkage maps for the CAD × ALF and CAD × HUE F₂ populations.** The table reports the number of markers retained after filtering, the number of linkage groups (LGs), and key descriptors of map length and LG size for each cross and reference genome. Parenthesis in column *distorted markers* represent the number of severely distorted markers (*p* < 0.001).

| **Cross** | **Reference** | **Markers** | **Number of LGs** | **Total length (cM)** | **Mean LG (cM)** | **Smallest LG (cM)** | **Largest LG (cM)** | **Distorted markers** |
| --- | --- | --- | --- | --- | --- | --- | --- | --- |
| CAD x ALF | CAD | 3,463 | 40 | 2,547 | 56.6 | 42.4 | 98.6 | 407 (56) |
| CAD x ALF | ALF | 3,414 | 38 | 2,541 | 62.0 | 38.5 | 112.5 | 410 (57) |
| CAD x HUE | CAD | 3,200 | 40 | 2,672 | 59.4 | 44.8 | 103.3 | 476 (171) |

**Supplementary Table 8. Genetic lengths of subdivided linkage groups affected by segregation distortion.** This table details the subdivision of linkage groups (LGs) that exhibited inflated genetic distances due to segregation distortion in the CAD × ALF and CAD × HUE crosses. For each cross and reference genome, affected LGs were manually split into multiple subdivisions (a–d), and the genetic length (cM) of each resulting segment is reported. The total genetic length after subdivision is shown in the *Total* column, with the corresponding genetic length prior to subdivision indicated in parentheses.

| **Cross** | **Reference** | **LG** | **subdivision** | **cM** | **Total** |
| --- | --- | --- | --- | --- | --- |
| CAD x ALF | CAD | 22 | a | 32.3 |  |
| CAD x ALF | CAD | 22 | b | 32.25 | 64.57 (84.28) |
| CAD x ALF | CAD | 29 | a | 53.98 | |
| CAD x ALF | CAD | 29 | b | 0.97 |  |
| CAD x ALF | CAD | 29 | c | 0 |  |
| CAD x ALF | CAD | 29 | d | 0 | 54.97 (126.67) |
| CAD x ALF | CAD | 33 | a | 5.97 |  |
| CAD x ALF | CAD | 33 | b | 53.60 | 59.57 (73.79) |
| CAD x ALF | ALF | 15 | a | 47.80 | |
| CAD x ALF | ALF | 15 | b | 0.99 | 48.79 (69.90) |
| CAD x ALF | ALF | 22 | a | 32.59 | |
| CAD x ALF | ALF | 22 | b | 53.51 | 86.01 (91.16) |
| CAD x ALF | ALF | 32 | a | 32.57 | |
| CAD x ALF | ALF | 32 | b | 5.97 | 38.54 (50.31) |
| CAD x HUE | CAD | 22 | a | 34.80 | |
| CAD x HUE | CAD | 22 | b | 45.78 | 80.58 (91.76) |
| CAD x HUE | CAD | 29 | a | 43.64 | |
| CAD x HUE | CAD | 29 | b | 0 |  |
| CAD x HUE | CAD | 29 | c | 0.87 |  |
| CAD x HUE | CAD | 29 | d | 0.27 | 44.78 (109.67) |
| CAD x HUE | CAD | 33 | a | 4.66 |  |
| CAD x HUE | CAD | 33 | b | 48.81 | 53.47 (67.54) |

**Supplementary Table 9. Chromosome-level recombination statistics for the CAD × ALF and CAD × HUE crosses.** For each chromosome, genome size (Mb), total genetic map length (cM), and recombination rate (cM/Mb) are reported. The *readjusted* column indicates chromosomes with chromosomal rearrangement based on the synteny analysis.

| **Cross** | **Genome** | **Chromosome** | **Size Mb** | **total cM** | **recombination rate (cM/Mb)** | **reajusted** |
| --- | --- | --- | --- | --- | --- | --- |
| CAD x ALF | ALF | 1 | 16.36 | 88.23 | 5.39 | no |
| CAD x ALF | ALF | 2 | 14.11 | 98.78 | 7 | no |
| CAD x ALF | ALF | 3 | 13.87 | 88.72 | 6.4 | no |
| CAD x ALF | ALF | 4 | 13.75 | 112.45 | 8.18 | yes |
| CAD x ALF | ALF | 5 | 12.99 | 73.44 | 5.66 | no |
| CAD x ALF | ALF | 6 | 12.93 | 82.68 | 6.39 | no |
| CAD x ALF | ALF | 7 | 12.91 | 68.56 | 5.31 | no |
| CAD x ALF | ALF | 8 | 12.49 | 63.56 | 5.09 | no |
| CAD x ALF | ALF | 9 | 12.38 | 75.46 | 6.1 | no |
| CAD x ALF | ALF | 10 | 11.81 | 86.74 | 7.34 | no |
| CAD x ALF | ALF | 11 | 11.78 | 64.24 | 5.45 | no |
| CAD x ALF | ALF | 12 | 11.43 | 77.01 | 6.74 | no |
| CAD x ALF | ALF | 13 | 10.94 | 74.65 | 6.82 | no |
| CAD x ALF | ALF | 14 | 10.78 | 76.53 | 7.1 | no |
| CAD x ALF | ALF | 15 | 10.49 | 47.8 | 4.56 | yes |
| CAD x ALF | ALF | 16 | 10.48 | 111.25 | 10.61 | yes |
| CAD x ALF | ALF | 17 | 10.19 | 61.55 | 6.04 | no |
| CAD x ALF | ALF | 18 | 10.16 | 72.73 | 7.16 | no |
| CAD x ALF | ALF | 19 | 9.77 | 57.49 | 5.88 | no |
| CAD x ALF | ALF | 20 | 9.6 | 52.67 | 5.49 | no |
| CAD x ALF | ALF | 21 | 9.25 | 69.58 | 7.52 | no |
| CAD x ALF | ALF | 22 | 9.09 | 53.51 | 5.89 | yes |
| CAD x ALF | ALF | 23 | 8.99 | 66.82 | 7.43 | no |
| CAD x ALF | ALF | 24 | 8.45 | 61.57 | 7.28 | no |
| CAD x ALF | ALF | 25 | 8.45 | 58.84 | 6.96 | no |
| CAD x ALF | ALF | 26 | 8.29 | 57.68 | 6.96 | no |
| CAD x ALF | ALF | 27 | 8.19 | 48.12 | 5.88 | no |
| CAD x ALF | ALF | 28 | 8.14 | 53.49 | 6.57 | no |
| CAD x ALF | ALF | 29 | 8.06 | 56.87 | 7.06 | no |
| CAD x ALF | ALF | 30 | 7.93 | 46.82 | 5.91 | no |
| CAD x ALF | ALF | 31 | 7.56 | 55.61 | 7.36 | no |
| CAD x ALF | ALF | 32 | 7.54 | 32.57 | 4.32 | yes |
| CAD x ALF | ALF | 33 | 7.45 | 50.66 | 6.8 | no |
| CAD x ALF | ALF | 34 | 7.15 | 47.76 | 6.67 | no |
| CAD x ALF | ALF | 35 | 6.82 | 56.68 | 8.31 | no |
| CAD x ALF | ALF | 36 | 6.14 | 62.15 | 10.12 | no |
| CAD x ALF | ALF | 37 | 5.69 | 42.86 | 7.53 | no |
| CAD x ALF | ALF | 38 | 5.05 | 45.69 | 9.05 | no |
| CAD x ALF | CAD | 1 | 16.53 | 88.19 | 5.33 | no |
| CAD x ALF | CAD | 2 | 14.84 | 98.64 | 6.65 | no |
| CAD x ALF | CAD | 3 | 14.01 | 88.7 | 6.33 | no |
| CAD x ALF | CAD | 4 | 12.82 | 73.44 | 5.73 | no |
| CAD x ALF | CAD | 5 | 12.82 | 63.53 | 4.96 | no |
| CAD x ALF | CAD | 6 | 12.81 | 82.64 | 6.45 | no |
| CAD x ALF | CAD | 7 | 12.76 | 72.29 | 5.66 | no |
| CAD x ALF | CAD | 8 | 12.51 | 75.44 | 06.03 | no |
| CAD x ALF | CAD | 9 | 12.17 | 86.75 | 7.13 | no |
| CAD x ALF | CAD | 10 | 12.05 | 63.86 | 5.3 | no |
| CAD x ALF | CAD | 11 | 11.54 | 77.45 | 6.71 | no |
| CAD x ALF | CAD | 12 | 10.99 | 74.7 | 6.8 | no |
| CAD x ALF | CAD | 13 | 10.69 | 76.51 | 7.16 | no |
| CAD x ALF | CAD | 14 | 10.27 | 61.52 | 5.99 | no |
| CAD x ALF | CAD | 15 | 10.01 | 72.86 | 7.28 | no |
| CAD x ALF | CAD | 16 | 9.48 | 53.72 | 5.67 | no |
| CAD x ALF | CAD | 17 | 9.4 | 52.66 | 5.6 | no |
| CAD x ALF | CAD | 18 | 9.38 | 62.64 | 6.68 | no |
| CAD x ALF | CAD | 19 | 9.34 | 54.33 | 5.82 | yes |
| CAD x ALF | CAD | 20 | 9.06 | 67.04 | 7.4 | no |
| CAD x ALF | CAD | 21 | 9.01 | 69.65 | 7.73 | no |
| CAD x ALF | CAD | 22 | 8.67 | 32.32 | 3.73 | yes |
| CAD x ALF | CAD | 23 | 8.58 | 57.53 | 6.71 | no |
| CAD x ALF | CAD | 24 | 8.45 | 47.69 | 5.64 | yes |
| CAD x ALF | CAD | 25 | 8.44 | 48.3 | 5.72 | no |
| CAD x ALF | CAD | 26 | 8.42 | 56.53 | 6.72 | no |
| CAD x ALF | CAD | 27 | 8.39 | 58.94 | 07.02 | no |
| CAD x ALF | CAD | 28 | 8.38 | 56.48 | 6.74 | no |
| CAD x ALF | CAD | 29 | 8.34 | 53.98 | 6.47 | yes |
| CAD x ALF | CAD | 30 | 8.21 | 53.5 | 6.52 | no |
| CAD x ALF | CAD | 31 | 8.1 | 55.65 | 6.87 | no |
| CAD x ALF | CAD | 32 | 7.91 | 46.51 | 5.88 | no |
| CAD x ALF | CAD | 33 | 7.79 | 53.6 | 6.88 | yes |
| CAD x ALF | CAD | 34 | 7.64 | 50.6 | 6.63 | no |
| CAD x ALF | CAD | 35 | 7.22 | 47.79 | 6.62 | no |
| CAD x ALF | CAD | 36 | 7.04 | 56.84 | 08.08 | no |
| CAD x ALF | CAD | 37 | 5.99 | 62.27 | 10.4 | no |
| CAD x ALF | CAD | 38 | 5.34 | 42.85 | 08.02 | no |
| CAD x ALF | CAD | 39 | 4.44 | 54.03 | 12.17 | yes |
| CAD x ALF | CAD | 40 | 4.08 | 56.08 | 13.74 | yes |
| CAD x HUE | CAD | 1 | 16.53 | 94.77 | 5.73 | no |
| CAD x HUE | CAD | 2 | 14.84 | 103.28 | 6.96 | no |
| CAD x HUE | CAD | 3 | 14.01 | 95.54 | 6.82 | no |
| CAD x HUE | CAD | 4 | 12.82 | 77.58 | 6.05 | no |
| CAD x HUE | CAD | 5 | 12.82 | 66.86 | 5.21 | no |
| CAD x HUE | CAD | 6 | 12.81 | 89.5 | 6.99 | no |
| CAD x HUE | CAD | 7 | 12.76 | 83.68 | 6.56 | no |
| CAD x HUE | CAD | 8 | 12.51 | 77.19 | 6.17 | no |
| CAD x HUE | CAD | 9 | 12.17 | 94.9 | 7.8 | no |
| CAD x HUE | CAD | 10 | 12.05 | 84.98 | 7.05 | no |
| CAD x HUE | CAD | 11 | 11.54 | 83.53 | 7.24 | no |
| CAD x HUE | CAD | 12 | 10.99 | 80.45 | 7.32 | no |
| CAD x HUE | CAD | 13 | 10.69 | 81.4 | 7.62 | no |
| CAD x HUE | CAD | 14 | 10.27 | 70.51 | 6.87 | no |
| CAD x HUE | CAD | 15 | 10.01 | 75.53 | 7.55 | no |
| CAD x HUE | CAD | 16 | 9.48 | 55.77 | 5.88 | no |
| CAD x HUE | CAD | 17 | 9.4 | 64.19 | 6.83 | no |
| CAD x HUE | CAD | 18 | 9.38 | 61.06 | 6.51 | no |
| CAD x HUE | CAD | 19 | 9.34 | 53.36 | 5.71 | yes |
| CAD x HUE | CAD | 20 | 9.06 | 63.14 | 6.97 | no |
| CAD x HUE | CAD | 21 | 09.01 | 67.41 | 7.48 | no |
| CAD x HUE | CAD | 22 | 8.67 | 45.78 | 5.28 | yes |
| CAD x HUE | CAD | 23 | 8.58 | 55.89 | 6.51 | no |
| CAD x HUE | CAD | 24 | 8.45 | 45.6 | 5.4 | yes |
| CAD x HUE | CAD | 25 | 8.44 | 59.9 | 7.09 | no |
| CAD x HUE | CAD | 26 | 8.42 | 59.99 | 7.13 | no |
| CAD x HUE | CAD | 27 | 8.39 | 63 | 7.51 | no |
| CAD x HUE | CAD | 28 | 8.38 | 57.52 | 6.87 | no |
| CAD x HUE | CAD | 29 | 8.34 | 43.64 | 5.23 | yes |
| CAD x HUE | CAD | 30 | 8.21 | 62.46 | 7.61 | no |
| CAD x HUE | CAD | 31 | 8.1 | 53.33 | 6.58 | no |
| CAD x HUE | CAD | 32 | 7.91 | 55.49 | 7.01 | no |
| CAD x HUE | CAD | 33 | 7.79 | 48.81 | 6.27 | yes |
| CAD x HUE | CAD | 34 | 7.64 | 55.23 | 7.23 | no |
| CAD x HUE | CAD | 35 | 7.22 | 54.65 | 7.57 | no |
| CAD x HUE | CAD | 36 | 7.04 | 51.92 | 7.38 | no |
| CAD x HUE | CAD | 37 | 5.99 | 49.23 | 8.22 | no |
| CAD x HUE | CAD | 38 | 5.34 | 47.05 | 8.81 | no |
| CAD x HUE | CAD | 39 | 4.44 | 49.6 | 11.17 | yes |
| CAD x HUE | CAD | 40 | 4.08 | 47.77 | 11.7 | yes |

**Supplementary Table 10. Multi-scale comparison of recombination rates (cM/Mb) between rearranged and non-rearranged chromosomes across crosses and parental backgrounds.** For each cross–genome combination (CAD × ALF with ALF parental map, CAD × ALF with CAD parental map, and CAD × HUE with CAD parental map), the table reports the number of markers, global weighted recombination rates (total cM / total Mb), mean marker-level recombination rates, and effect sizes estimated from LMM and GAMLSS analyses. Effect sizes are presented as estimates ± standard error (SE) with corresponding *p*-values. Significance levels: *p* < 0.05 (**), p < 0.01 (**), p < 0.001 (****).

| **Cross** | **Parent** | **n markers in rearranged / not rearranged chromosomes** | **Global recombination rate in rearranged / not rearranged chromosomes (cM/Mb)** | **Markers mean recombination rate in rearranged / not rearranged chromosomes (cM/Mb)** | **LMM Effect Size ± SE (p-value)** | **GAMLSS μ Effect (Mean) ± SE (p-value)** | **GAMLSS σ Effect (Variance) ± SE (p-value)** |
| --- | --- | --- | --- | --- | --- | --- | --- |
| CAD x ALF | ALF | 425 / 2919 | 7.73 / 6.58 | 8.46 / 6.93 | 1.485 ± 0.643 (0.027*) | 1.346 ± 1.102 (0.231) | 0.884 ± 0.341 (0.014*) |
| CAD x ALF | CAD | 432 / 2974 | 7.65 / 6.48 | 8.07 / 6.91 | 1.391 ± 0.673 (0.047*) | 1.264 ± 1.409 (0.376) | 1.155 ± 0.362 (0.003**) |
| CAD x HUE | CAD | 405 / 2735 | 7.34 / 6.90 | 7.84 / 7.23 | 0.609 ± 0.514 (0.236) | 1.637 ± 1.428 (0.260) | 1.895 ± 0.383 (<0.001***) |

**Supplementary Table 11.** **Descriptive statistics for reproductive, vegetative, and biomass-related traits measured in the CAD × ALF and CAD × HUE populations**. For each trait, the category (reproductive and/or vegetative), the number of individuals (N), mean, standard deviation (SD), minimum (Min), maximum (Max), and coefficient of variation (CV%) are presented.

| **Cross** | **Category** | **Trait** | **N** | **Mean** | **SD** | **Min** | **Max** | **CV (%)** |
| --- | --- | --- | --- | --- | --- | --- | --- | --- |
| CAD x ALF | Reproductive | Number of male spikes | 48 | 03.08 | 4.47 | 0 | 19 | 145.02 |
| CAD x ALF | Reproductive | Number of female spikes | 48 | 3.85 | 6.90 | 0 | 35 | 179.04 |
| CAD x ALF | Reproductive | Number of androgynous spikes | 48 | 0.12 | 0.33 | 0 | 1 | 267.37 |
| CAD x ALF | Reproductive | Number of floral stems | 48 | 03.02 | 4.39 | 0 | 19 | 145.42 |
| CAD x ALF | Reproductive | Number of utricles | 48 | 0.81 | 4.35 | 0 | 30 | 535.97 |
| CAD x ALF | Reproductive | Male spike weight | 48 | 0.06 | 0.09 | 0 | 0.34 | 154.56 |
| CAD x ALF | Reproductive | Female spike weight | 48 | 0.02 | 0.05 | 0 | 0.26 | 230.38 |
| CAD x ALF | Reproductive | Stem weight | 48 | 1.60 | 2.21 | 0 | 10.06 | 138.27 |
| CAD x ALF | Reproductive | Utricle weight | 48 | 0.00 | 0.01 | 0 | 0.04 | 547.54 |
| CAD x ALF | Vegetative | Maximum leaf width | 48 | 8.25 | 1.72 | 6.00 | 10.25 | 20.85 |
| CAD x ALF | Vegetative | Maximum leaf length | 48 | 53.66 | 13.19 | 32.30 | 78.10 | 24.58 |
| CAD x ALF | Vegetative | Leaf weight | 48 | 16.03 | 8.77 | 1.93 | 38.75 | 54.71 |
| CAD x ALF | Vegetative and Reproductive | Aerial part weight | 48 | 17.71 | 9.35 | 4.61 | 43.21 | 52.82 |
| CAD x ALF | Vegetative | Rhizome weight | 48 | 3.18 | 3.97 | 0.27 | 27.80 | 124.81 |
| CAD x ALF | Reproductive and vegetative | Plant weight | 48 | 20.89 | 10.86 | 5.01 | 48.82 | 51.98 |
| CAD x HUE | Reproductive | Number of male spikes | 107 | 3.18 | 4.64 | 0 | 27 | 145.88 |
| CAD x HUE | Reproductive | Number of female spikes | 107 | 3.7 | 5.97 | 0 | 36 | 161.41 |
| CAD x HUE | Reproductive | Number of androgynous spikes | 107 | 0.5 | 1.15 | 0 | 6 | 228.33 |
| CAD x HUE | Reproductive | Number of floral stems | 107 | 3.13 | 4.54 | 0 | 27 | 144.99 |
| CAD x HUE | Reproductive | Number of utricles | 107 | 1.12 | 7.29 | 0 | 72 | 649.92 |
| CAD x HUE | Reproductive | Male spike weight | 107 | 0.07 | 0.1 | 0 | 0.48 | 144.98 |
| CAD x HUE | Reproductive | Female spike weight | 107 | 0.03 | 0.07 | 0 | 0.39 | 208.05 |
| CAD x HUE | Reproductive | Stem weight | 107 | 1.81 | 2.6 | 0 | 13.62 | 143.97 |
| CAD x HUE | Reproductive | Utricle weight | 107 | 0 | 0.01 | 0 | 0.06 | 613.37 |
| CAD x HUE | Vegetative | Maximum leaf width | 107 | 10.08 | 2.12 | 5 | 15 | 21.08 |
| CAD x HUE | Vegetative | Maximum leaf length | 107 | 50.79 | 8.76 | 29.5 | 75 | 17.24 |
| CAD x HUE | Vegetative | Leaf weight | 107 | 18.18 | 8.63 | 1.31 | 44.2 | 47.49 |
| CAD x HUE | Vegetative and Reproductive | Aerial part weight | 107 | 20.09 | 9.24 | 1.31 | 50.25 | 45.98 |
| CAD x HUE | Vegetative | Rhizome weight | 107 | 3.5 | 1.77 | 0.22 | 8.22 | 50.48 |
| CAD x HUE | Reproductive and vegetative | Plant weight | 107 | 23.59 | 10.8 | 1.53 | 57.76 | 45.73 |

**Supplementary Table 12. Summary of significant QTLs for reproductive traits.** Significant peaks were identified using a two-part model based on 1,000 genome-wide permutations (α = 0.05). Physical positions correspond to the ALF and CAD reference genomes.

| **Cross** | **Genome** | **Trait** | **Chr** | **Physical position (bp)** | **Segregation distortion** | **Genetic position (cM)** | **LOD** | **Permutation threshold** | **95% Bayes interval (cM)** | | **Variance explained (%)** |
| --- | --- | --- | --- | --- | --- | --- | --- | --- | --- | --- | --- |
| CAD × ALF | ALF | Number of male spikes | 16 | 8,354,929 | moderate | 63.35 | 8.80 | 8.01 | 60–83 | 12.30 | |
| CAD × ALF | ALF | Number of floral stems | 16 | 8,354,929 | moderate | 63.35 | 8.86 | 7.88 | 60–82 | 12.43 | |
| CAD × ALF | CAD | Number of male spikes | 40 | 1,925,252 | moderate | 8.24 | 8.80 | 8.09 | 5–28 | 12.33 | |
| CAD × ALF | CAD | Number of floral stems | 40 | 1,925,252 | moderate | 8.24 | 8.86 | 8.03 | 5–27 | 12.43 | |
| CAD × HUE | CAD | Weight of floral stems | 4 | 9,749,532 | none | 54 | 8.98 | 8.00 | 54–54 | 8.24 | |
| CAD × HUE | CAD | Weight of reproductive tissue | 4 | 9,749,532 | none | 54 | 8.98 | 8.05 | 54–54 | 8.20 | |

**Supplementary Figures**

**Supplementary Figure 1. Germination rate by population and cross type.** Boxplots showing germination rates across populations for different cross types. Colors indicate the generation: blue for selfed parents (P), yellow for F₁ hybrids, and red for F₂ progeny.

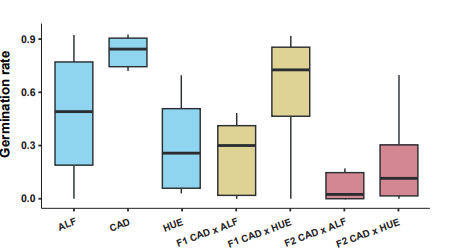

**Supplementary Figure 2. HiC and telomeres plot from the two parental genomes sequenced. a)** Whole-genome Hi-C interaction heatmaps for ALF (top) and CAD (bottom) genomes. The heatmaps display the density of pairwise chromatin interactions across all established pseudomolecules. **b)** Telomeric repeat identification (TIDK) of all preliminary pseudomolecules of ALF (top) and CAD (bottom) genomes. Line plots are aligned with the corresponding chromosomal coordinates shown above and indicate localized enrichment of the canonical plant telomeric repeat motif (TTTAGGG), visible as distinct density peaks.

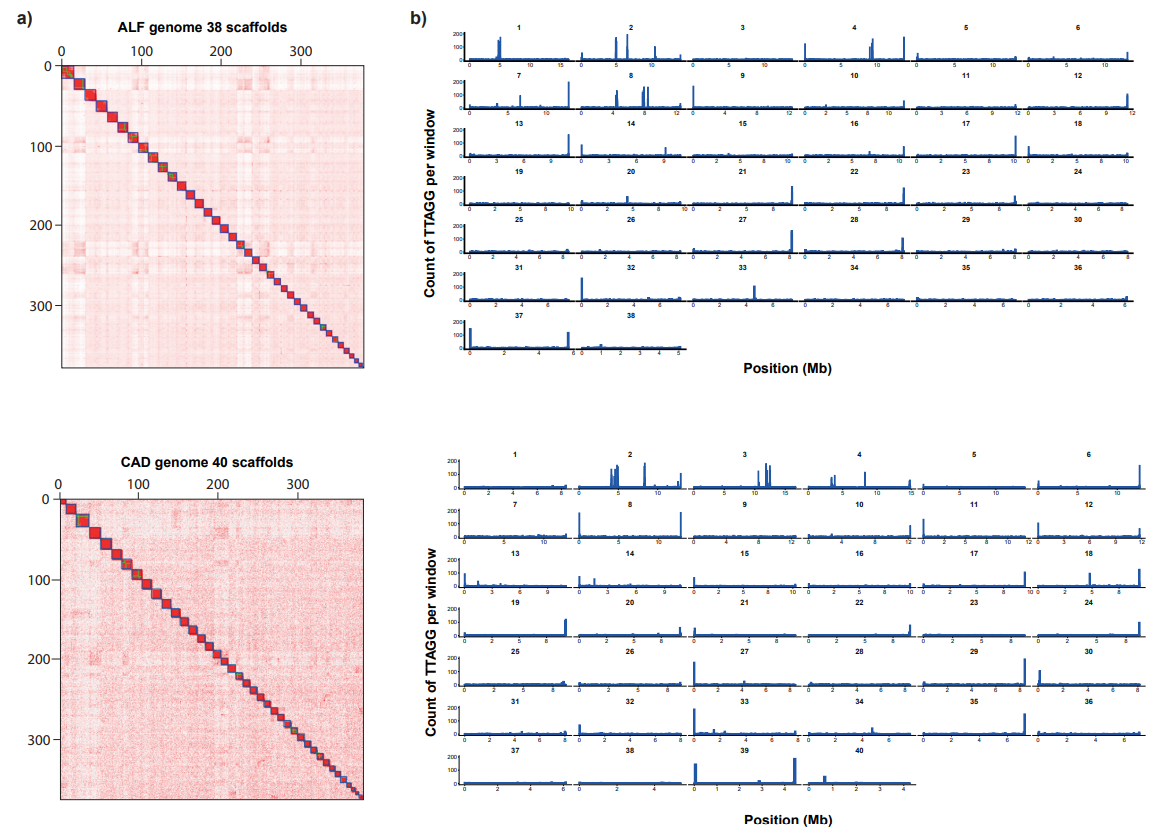

**Supplementary Figure 3.** **Synteny plot of *C. laevigata* genomes and haplotype assemblies.** Each genome and haplotype is shown as a single horizontal line divided into chromosomes. Syntenic regions are depicted as braids connecting homologous regions across genomes. Colored braids highlight chromosomes that show structural rearrangements (CRs) relative to the other genomes, while grey braids indicate non-rearranged chromosomes.

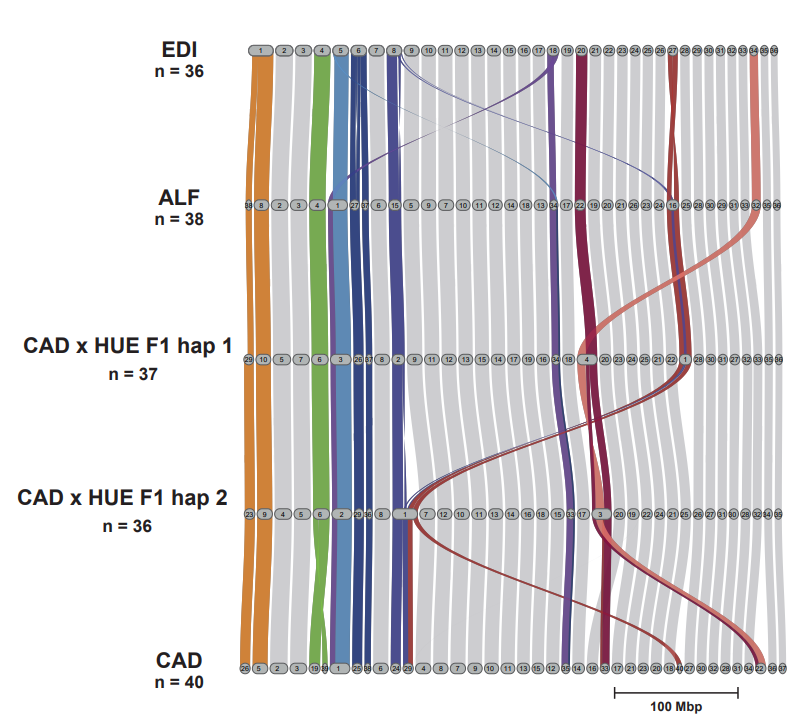

**Supplementary Figure 4. Genomic features along the EDI genome.** Proportion of coding sequences (CDS) and repeat elements, and GC content calculated in 50Kb sliding windows across the genome. Light blue vertical lines indicate breakpoint regions.

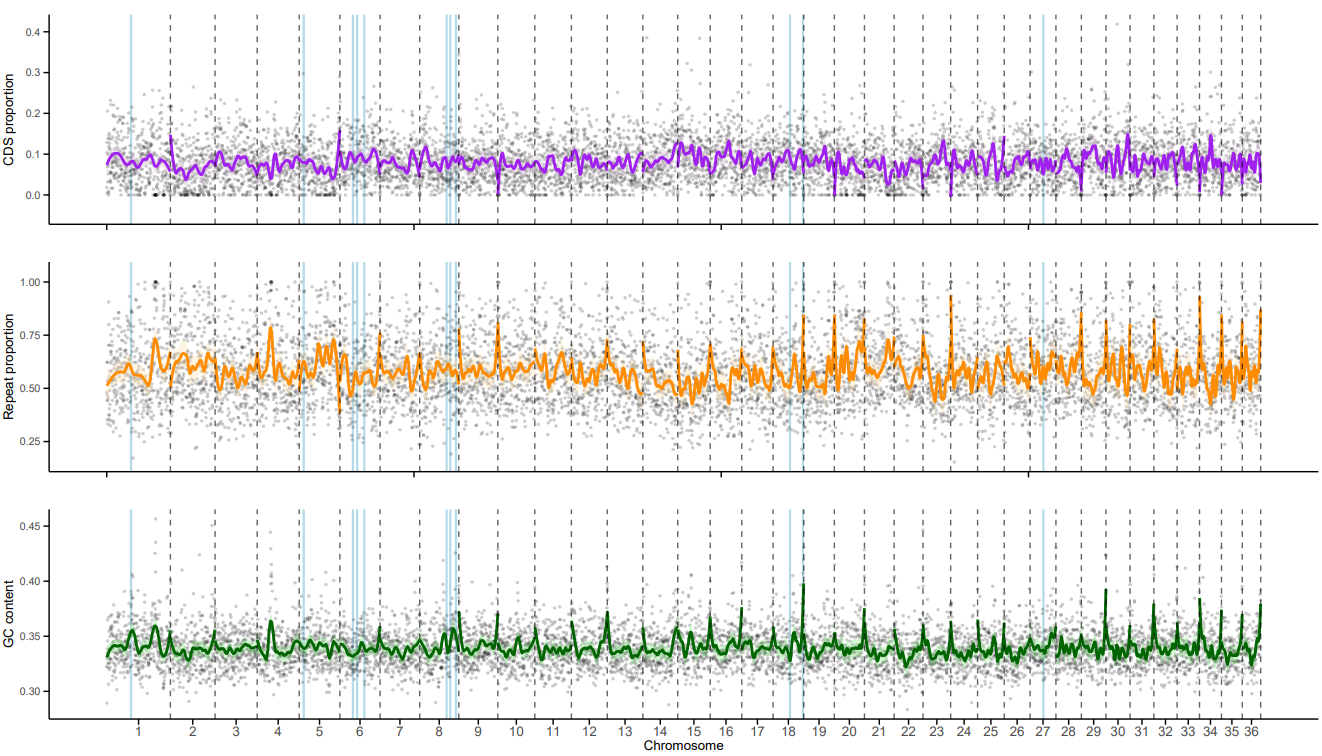

**Supplementary Figure 5. Genomic features along the ALF genome.** Proportion of coding sequences (CDS) and repeat elements, and GC content calculated in 50Kb sliding windows across the genome. Light blue vertical lines indicate breakpoint regions.

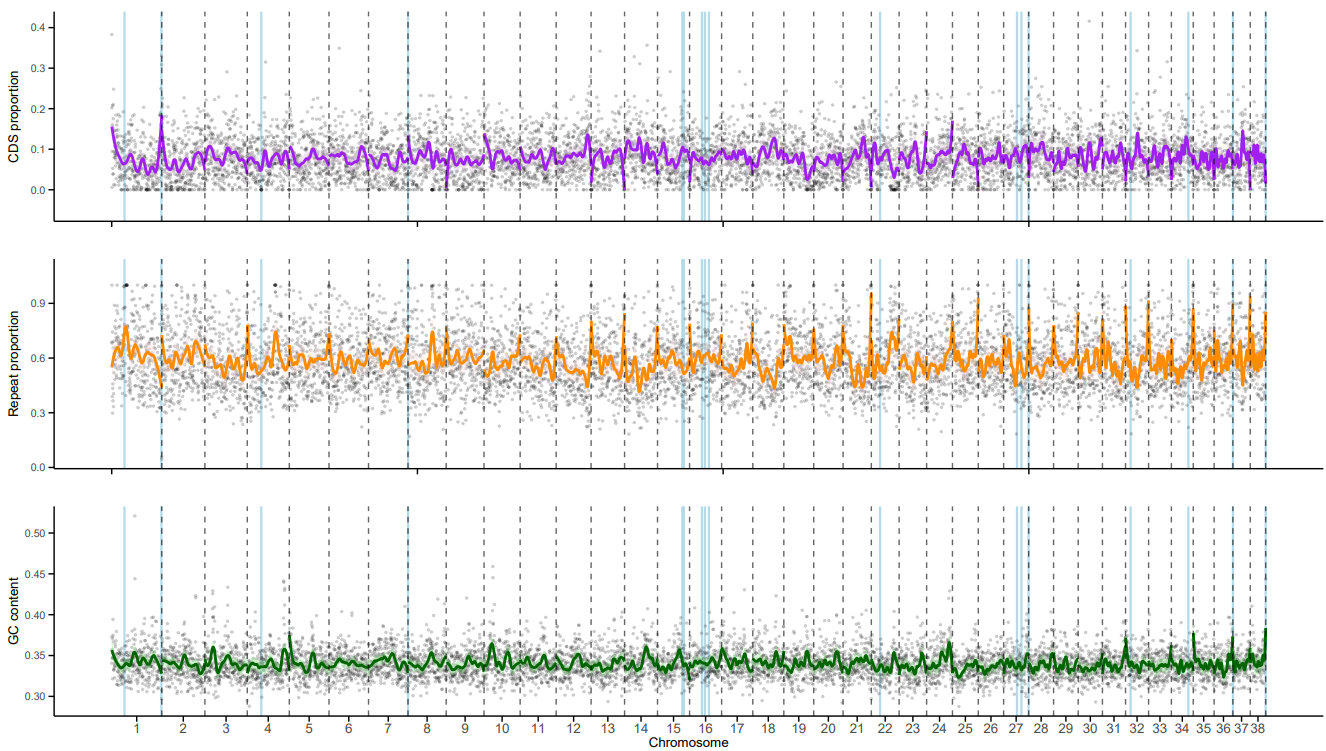

**Supplementary Figure 6. Genomic features along the CAD genome.** Proportion of coding sequences (CDS) and repeat elements, and GC content calculated in 50Kb sliding windows across the genome. Light blue vertical lines indicate breakpoint regions.

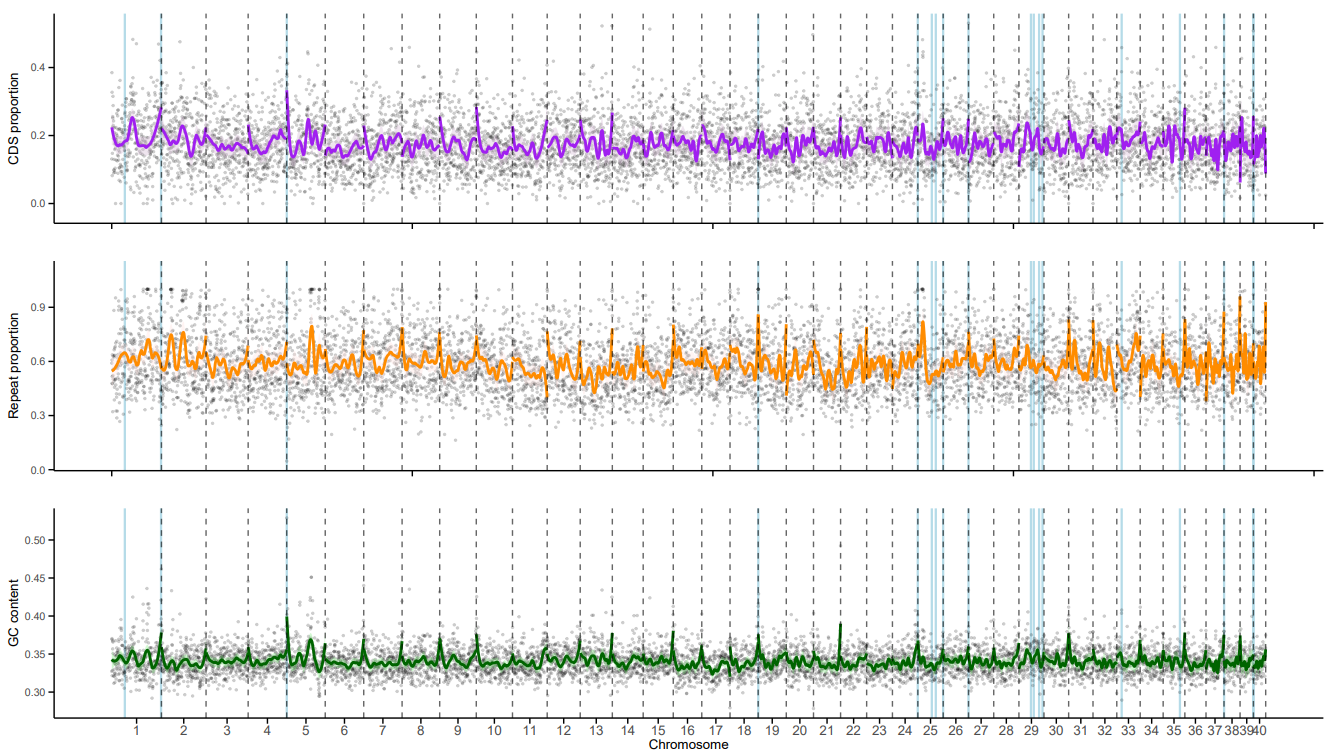

**Supplementary Figure 7. Correlation matrices of genomic features across EDI, ALF and CAD genomes.** The panels display pairwise correlations between GC content, Coding Sequences (CDS), total Repeats, and LTR gypsy elements. The diagonal shows the density distribution for each feature. The lower triangle contains scatter plots with linear regression lines representing the relationship between variables. The upper triangle provides the Pearson correlation coefficients (Corr), where asterisks (***) indicate statistical significance at *p* < 0.001.

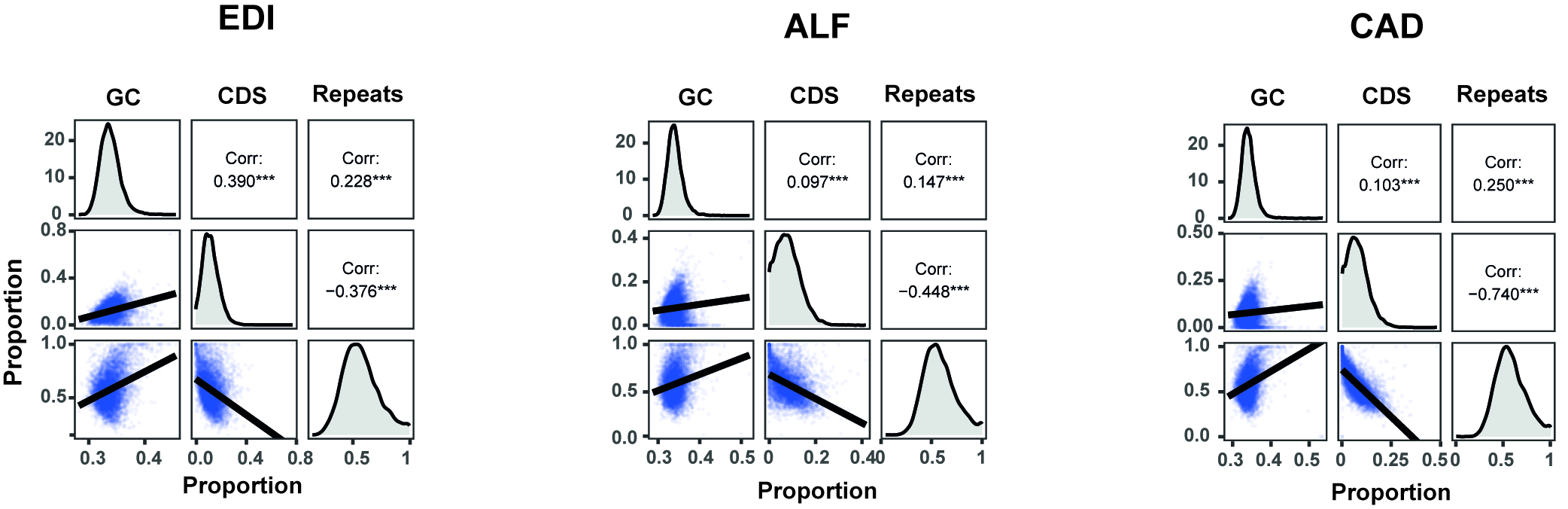

**Supplementary Figure 8. Genetic structure analyses with diemR of different crosses and datasets.** The first two rows correspond to markers from the CAD × ALF F₂ population, aligned to the ALF genome (top row) and the CAD genome (second row), respectively. The last row corresponds to markers from the CAD × HUE F₂ population aligned to the CAD genome. **a)** Polarization plots showing patterns of genetic polarization among samples/populations based on allele frequency estimates. **(b)** De Finetti triangular plots showing genotype distributions and relative allele frequencies.

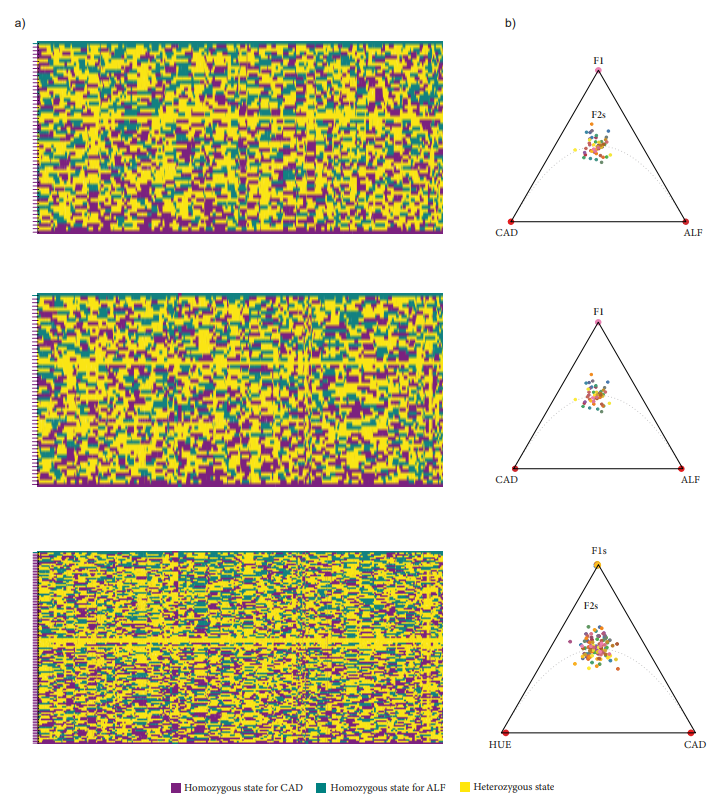

**Supplementary Figure 9. Recombination landscapes of the 38 chromosomes of the ALF genome based on CAD × ALF F₂ RAD-seq data.** For each chromosome, three panels are shown. The upper panel depicts the cumulative genetic distance (cM) along the chromosome, the middle panel shows the recombination rate expressed as cM/Mb, and the lower panel displays the recombination fraction (rf) matrix. In the upper and middle panels, black dots indicate markers without segregation distortion, whereas red and purple dots represent markers showing moderate (0.001 < *p* ≤ 0.05) and severe (*p* < 0.001) segregation distortion, respectively. Colours in the recombination fraction matrix range from red (rf = 0.5) to dark blue (rf = 0). Dashed lines in chromosomes 15, 22, and 32 indicate the separation into distinct linkage groups.

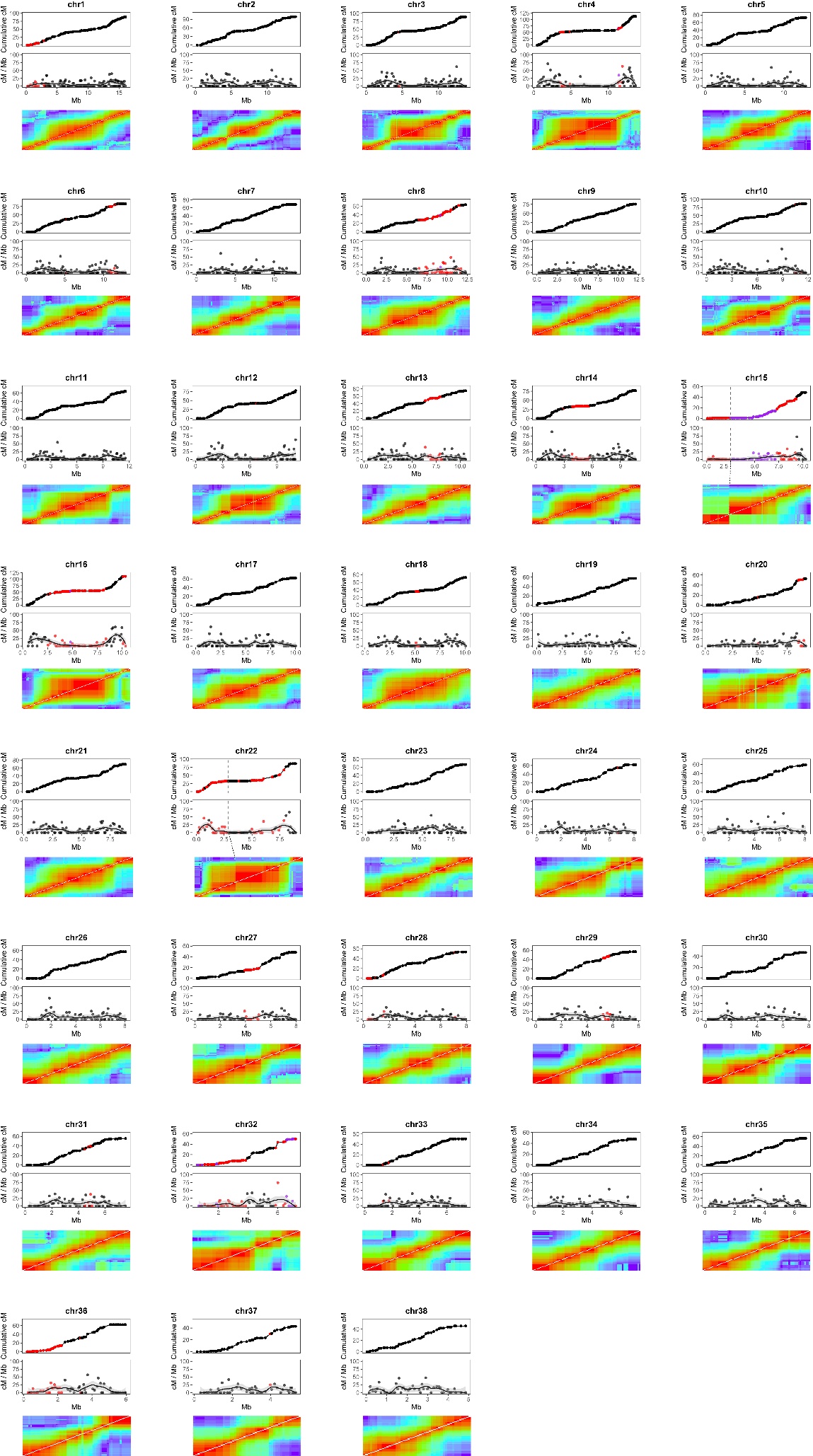

**Supplementary Figure 10. Recombination landscapes of the 40 chromosomes of the CAD genome based on CAD × ALF F₂ RAD-seq data.** For each chromosome, three panels are shown. The upper panel depicts the cumulative genetic distance (cM) along the chromosome, the middle panel shows the recombination rate expressed as cM/Mb, and the lower panel displays the recombination fraction (rf) matrix. In the upper and middle panels, black dots indicate markers without segregation distortion, whereas red and purple dots represent markers showing moderate (0.001 < *p* ≤ 0.05) and severe (*p* < 0.001) segregation distortion, respectively. Colours in the recombination fraction matrix range from red (rf = 0.5) to dark blue (rf = 0). Dashed lines in chromosomes 22, 29, and 33 indicate the separation into distinct linkage groups.

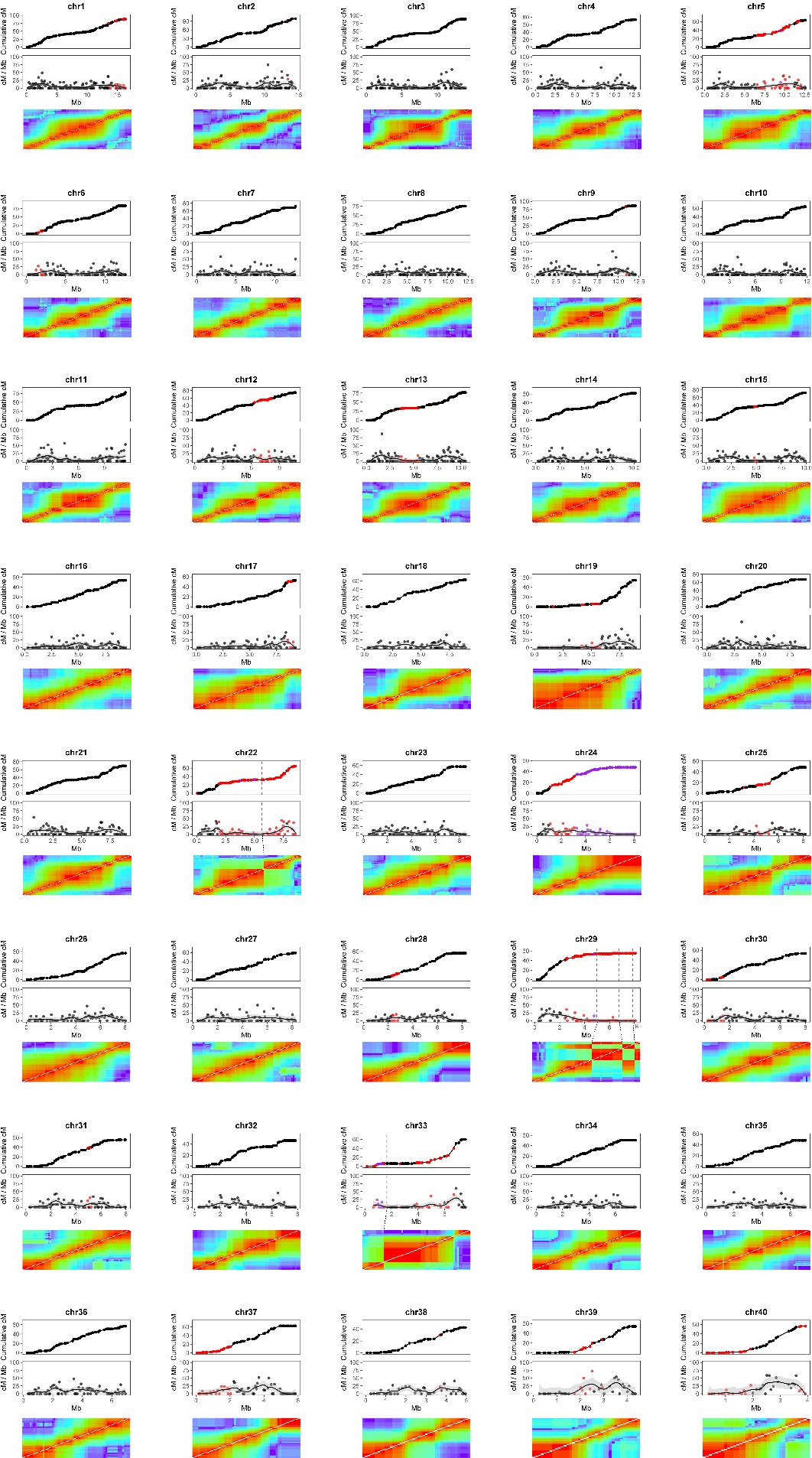

**Supplementary Figure 11. Recombination landscapes of the 40 chromosomes of the CAD genome based on CAD × HUE F₂ RAD-seq data.** For each chromosome, three panels are shown. The upper panel depicts the cumulative genetic distance (cM) along the chromosome, the middle panel shows the recombination rate expressed as cM/Mb, and the lower panel displays the recombination fraction (rf) matrix. In the upper and middle panels, black dots indicate markers without segregation distortion, whereas red and purple dots represent markers showing moderate (0.001 < *p* ≤ 0.05) and severe (*p* < 0.001) segregation distortion, respectively. Colours in the recombination fraction matrix range from red (rf = 0.5) to dark blue (rf = 0). Dashed lines in chromosomes 22, 29, and 33 indicate the separation into distinct linkage groups.

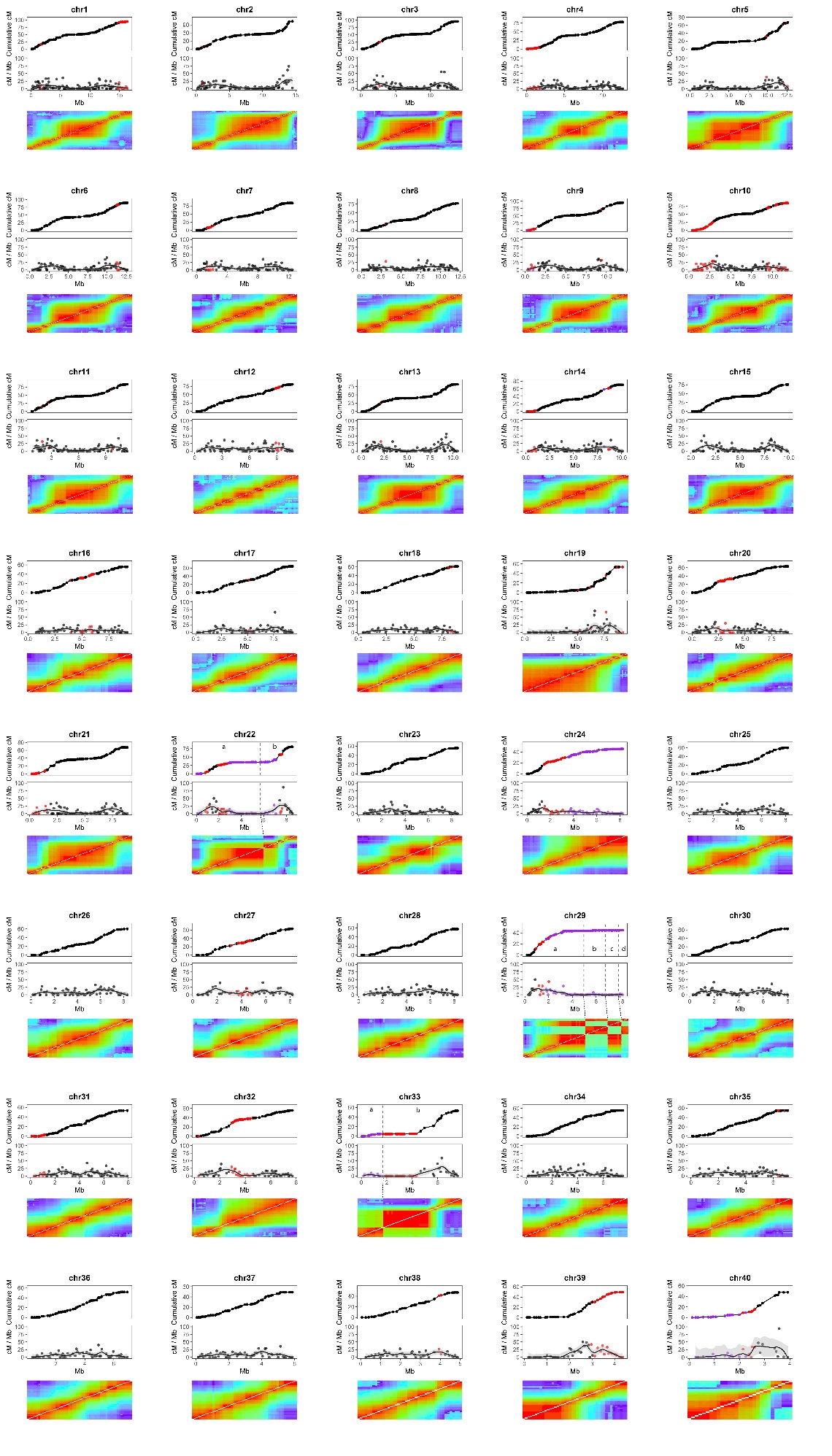

**Supplementary Figure 12. Linkage groups of the 38 chromosomes of the ALF genome based on CAD × ALF F₂ RAD-seq data, visualized with LinkageMapView.** Each linkage group is represented as a chromosome-like ideogram. Horizontal lines within each linkage group correspond to individual markers. The colour palette indicates marker density per centimorgan (cM).

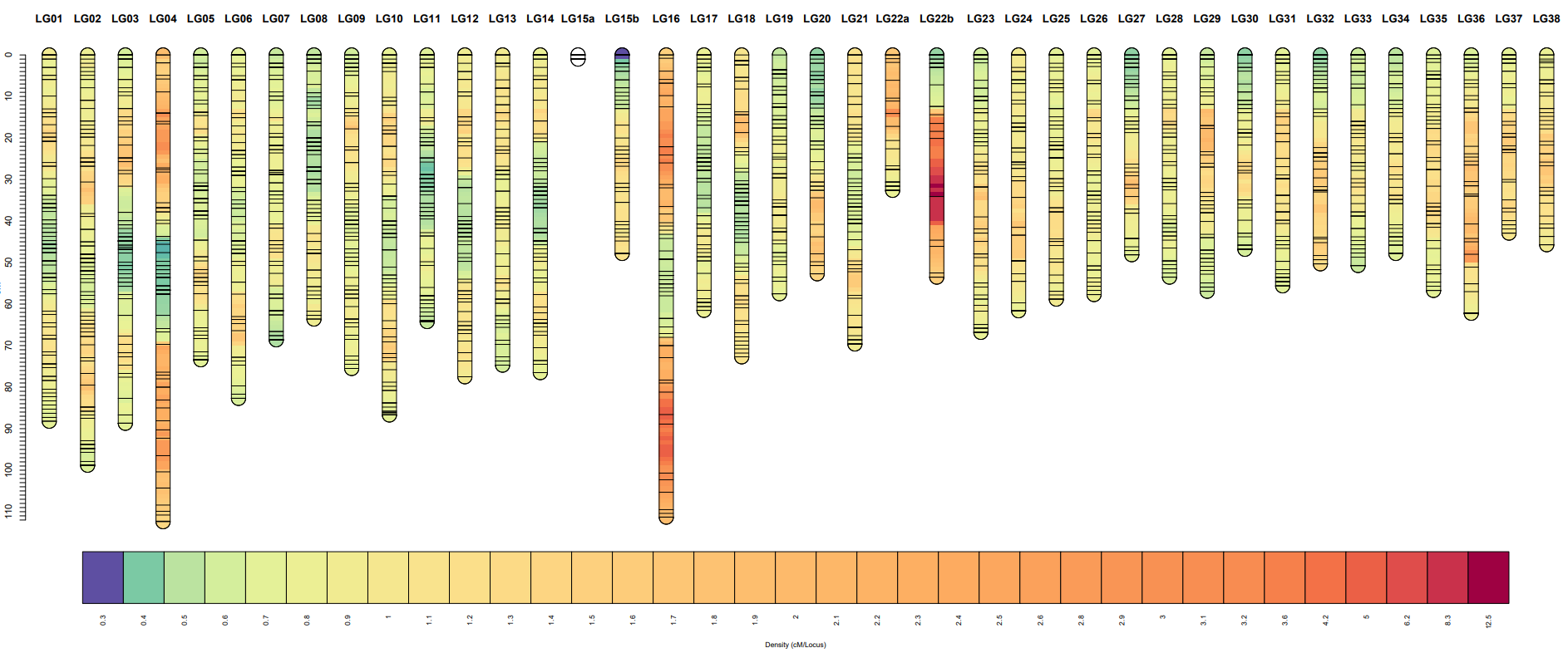

**Supplementary Figure 13. Linkage groups of the 40 chromosomes of the CAD genome based on CAD × ALF F₂ RAD-seq data, visualized with LinkageMapView.** Each linkage group is represented as a chromosome-like ideogram. Horizontal lines within each linkage group correspond to individual markers. The colour palette indicates marker density per centimorgan (cM).

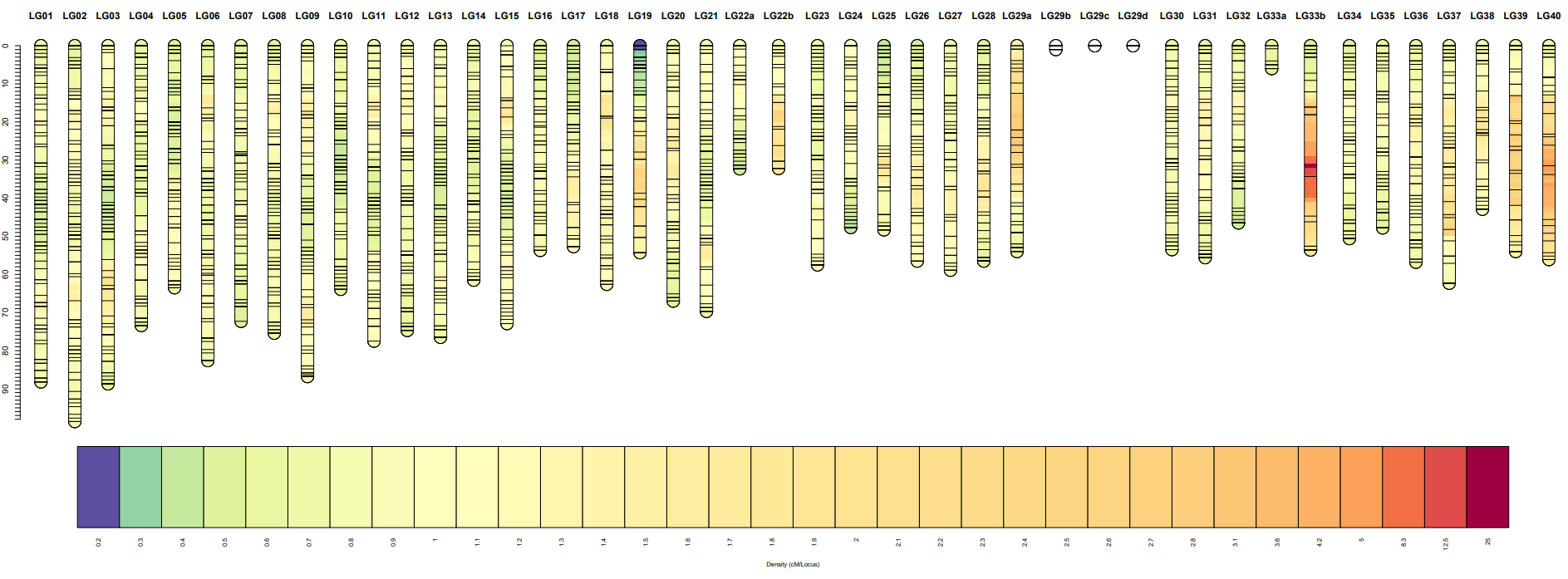

**Supplementary Figure 14.** **Linkage groups of the 40 chromosomes of the CAD genome based on CAD × HUE F₂ RAD-seq data, visualized with LinkageMapView.** Each linkage group is represented as a chromosome-like ideogram. Horizontal lines within each linkage group correspond to individual markers. The colour palette indicates marker density per centimorgan (cM).

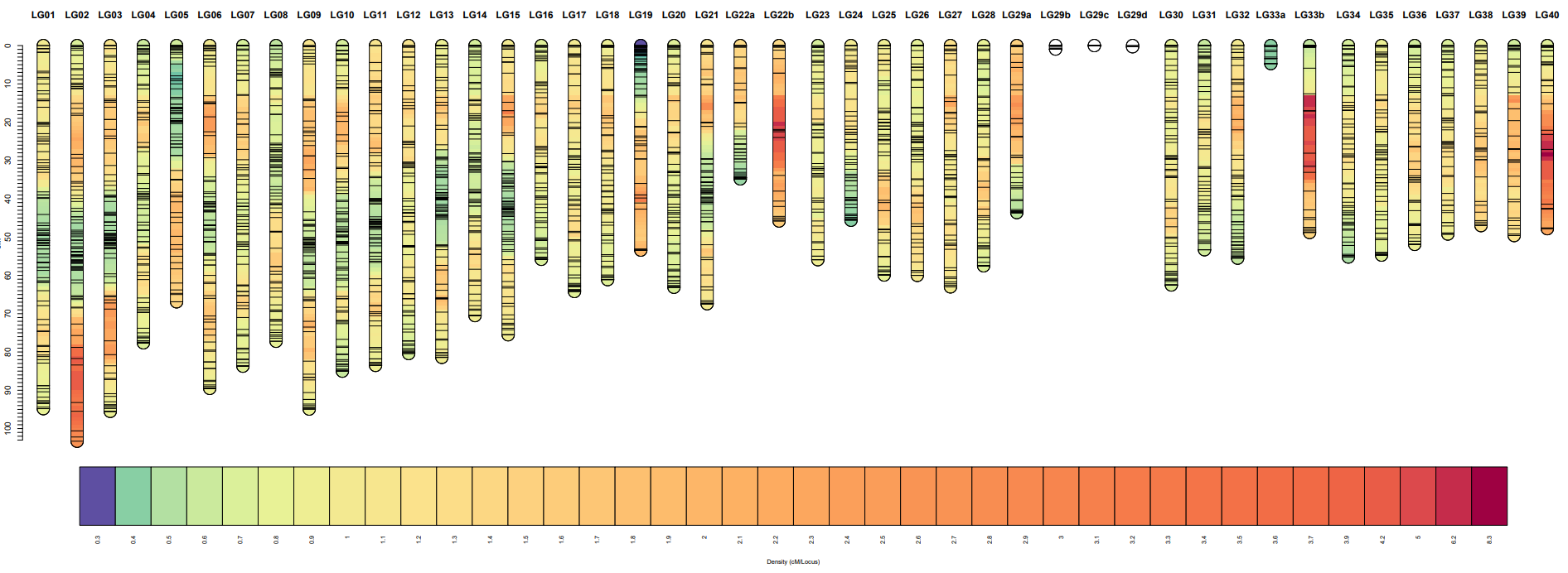

**Supplementary Figure 15. Relationship between chromosome size and recombination in ALF genome based on CAD × ALF F₂ RAD-seq data.** Recombination values for each chromosome are log-trans formed, and the line represents the fitted linear model.

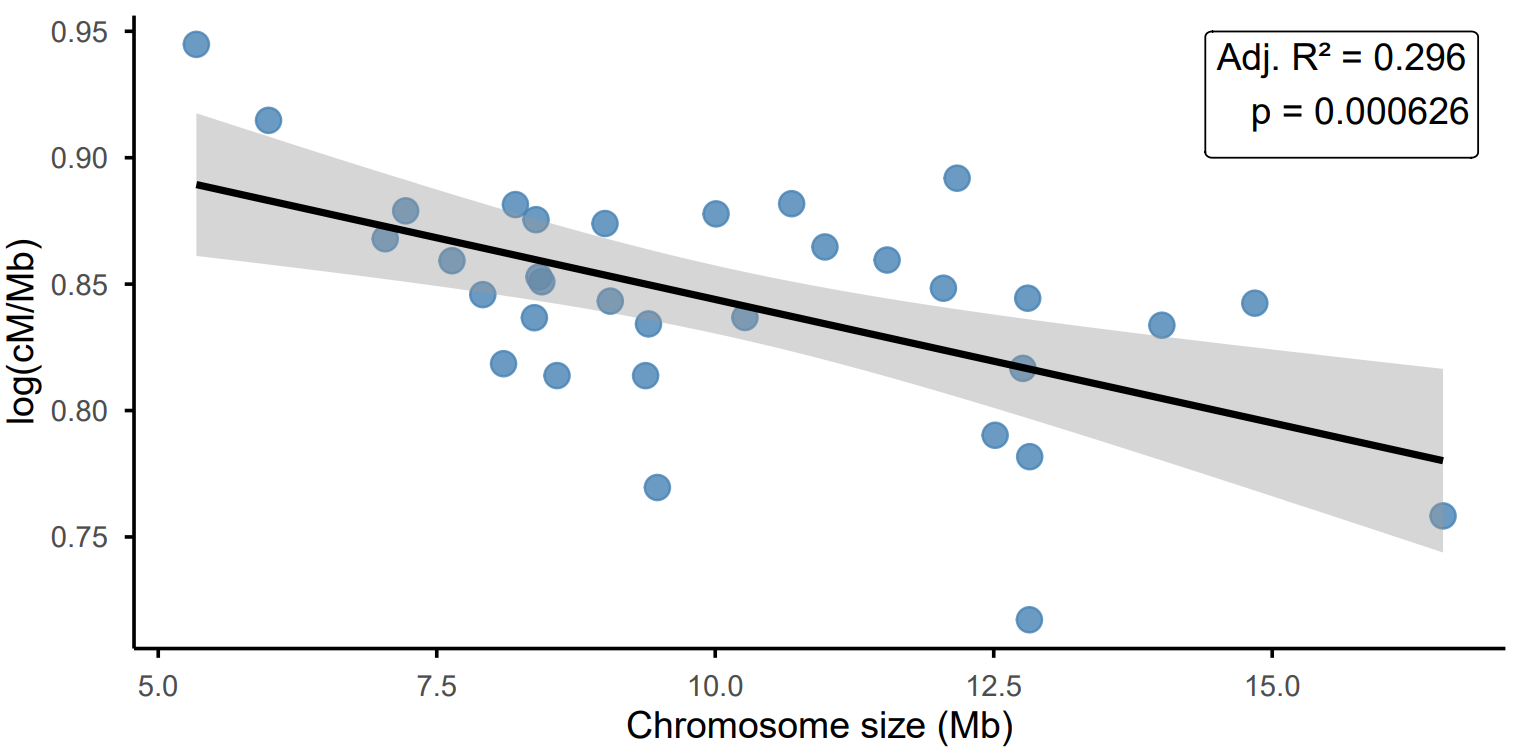

**Supplementary Figure 16. Relationship between chromosome size and recombination in CAD genome based on CAD × ALF F₂ RAD-seq data.** Recombination values for each chromosome are log-transformed, and the line represents the fitted linear model.

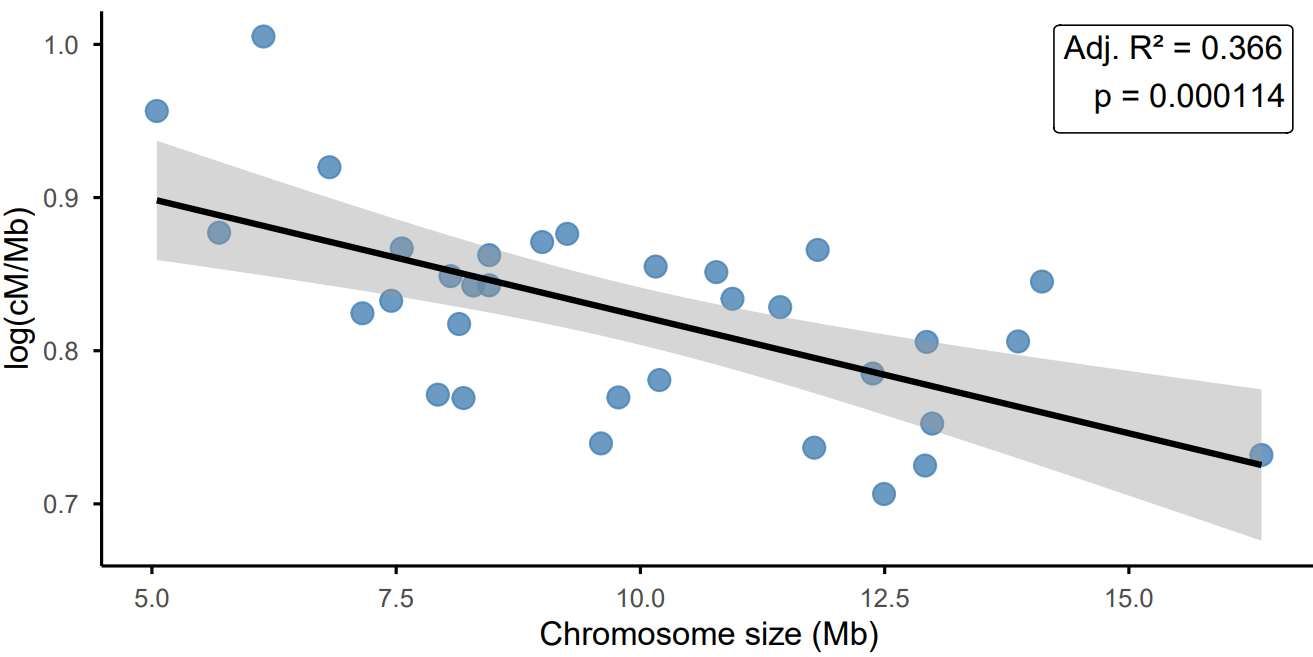

**Supplementary Figure 17. Relationship between chromosome size and recombination in CAD genome based on CAD × HUE F₂ RAD-seq data.** Recombination values for each chromosome are log-transformed, and the line represents the fitted linear model.

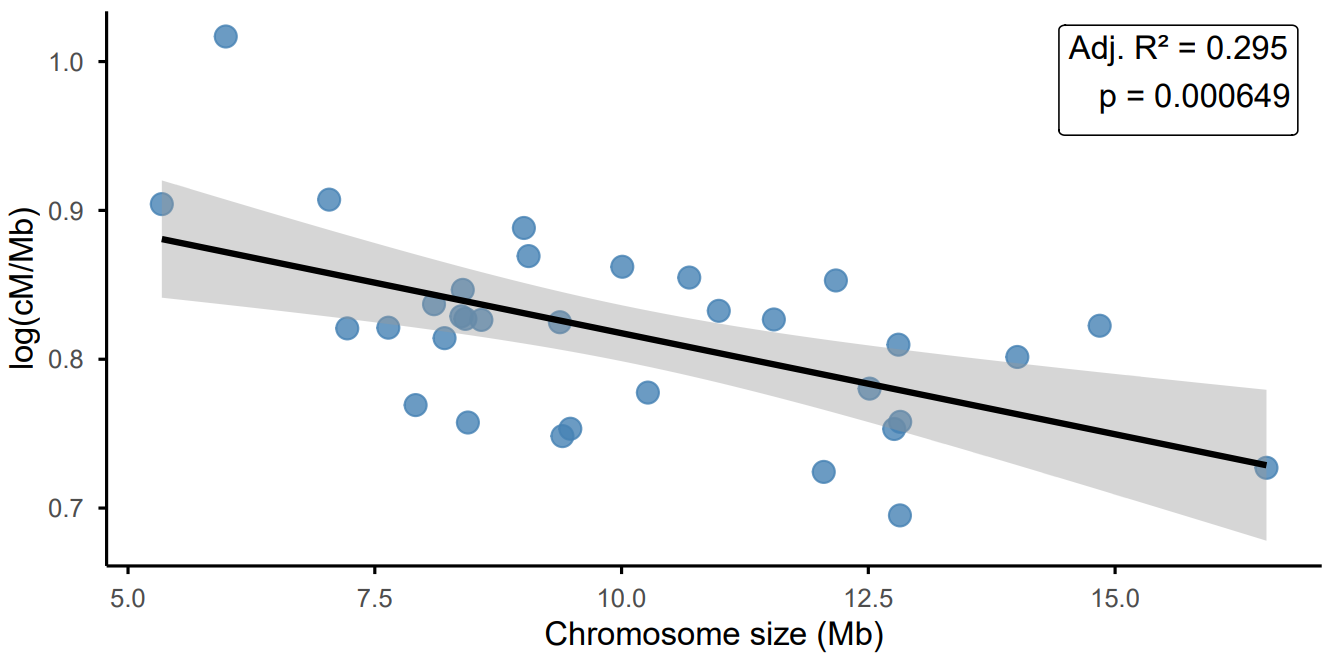

**Supplementary Figure 18. Spearman’s correlation matrix of phenotypic traits.** The matrix shows pairwise correlations among 15 morphological traits for the CAD x ALF (a) and CAD x HUE (b) F₂ population, with numerical values representing Spearman’s rank correlation coefficients (*r*).

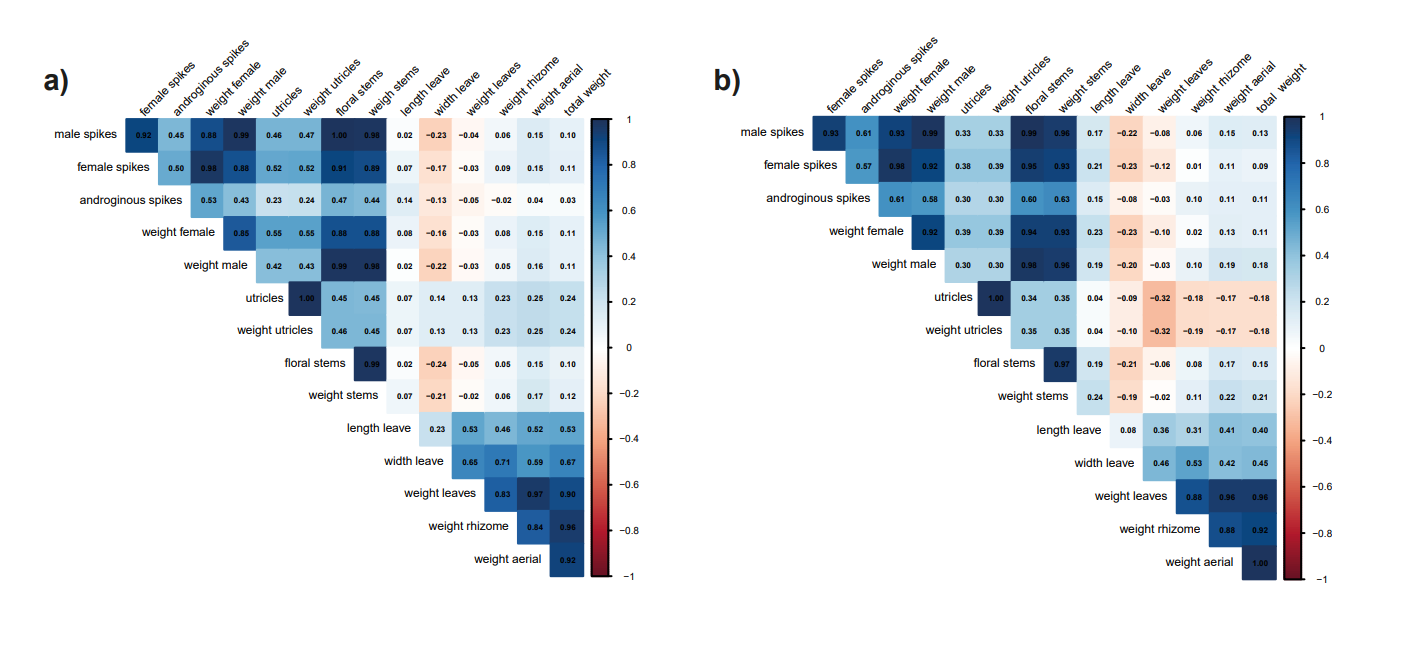

**Supplementary Figure 19. QTL mapping in the ALF genome when using the CAD x ALF F₂ individuals.** LOD score profiles for number of floral stems (green) and number of male spikes (orange) on ALF genome. Dashed lines represent the LOD threshold.

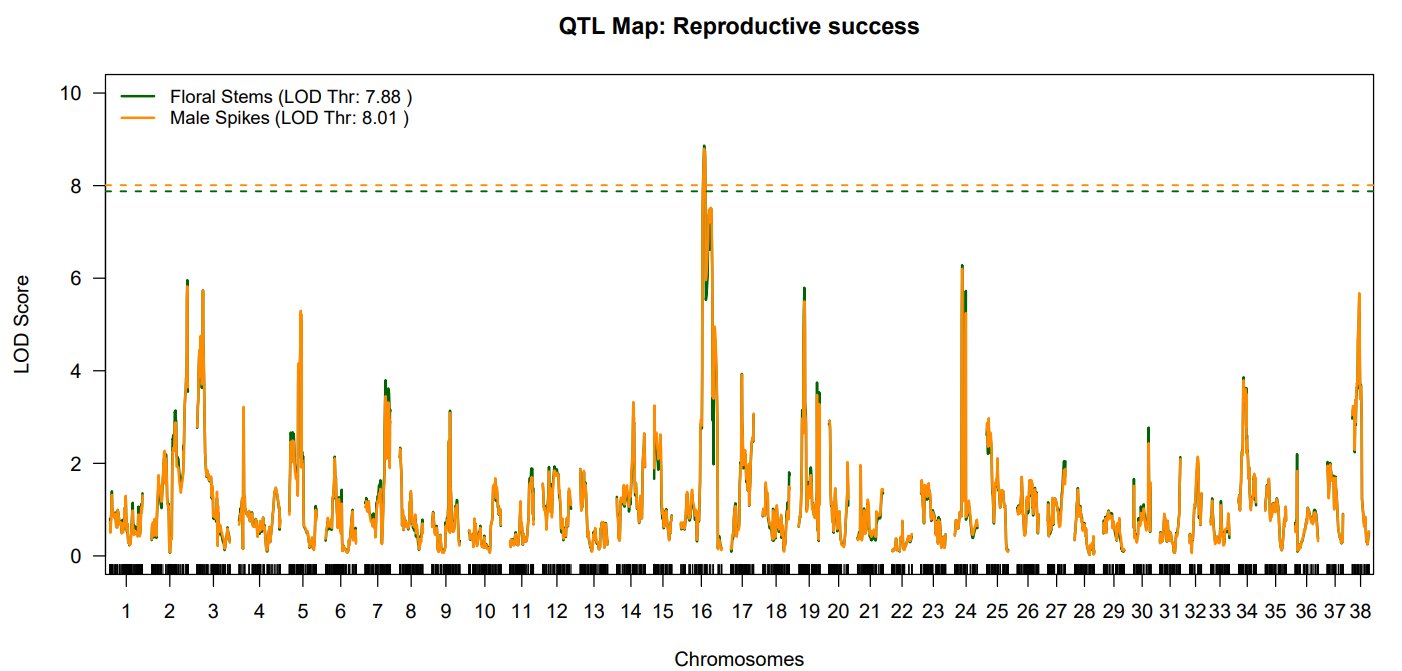

**Supplementary Figure 20. QTL mapping in CAD genome when using the CAD x ALF F₂ individuals.** LOD score profiles for number of floral stems (green) and number of male spikes (orange) on CAD genome. Dashed lines represent the LOD threshold.

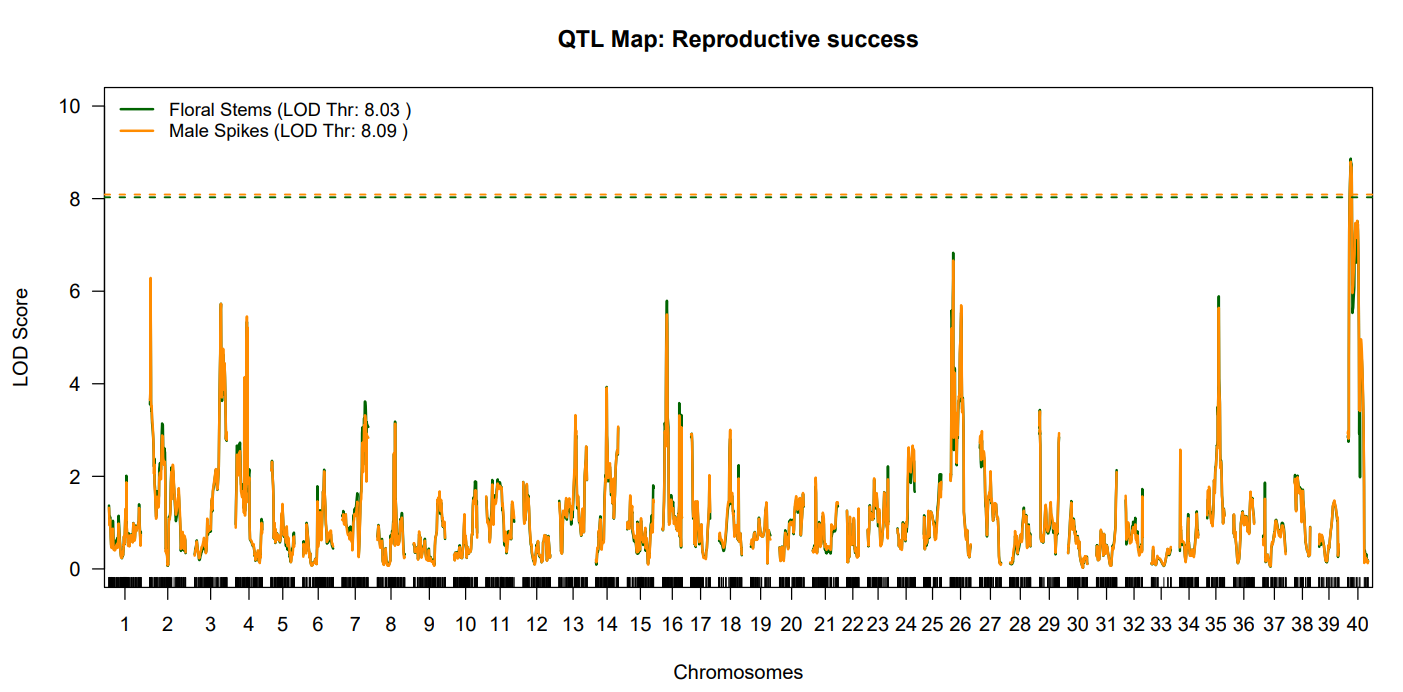

**Supplementary Figure 21. QTL mapping in CAD genome when using the CAD x HUE F₂ individuals.** LOD score profiles for floral stem weight (orange) and total reproductive biomass (green) are shown across the CAD genome. Dashed lines represent the LOD threshold.

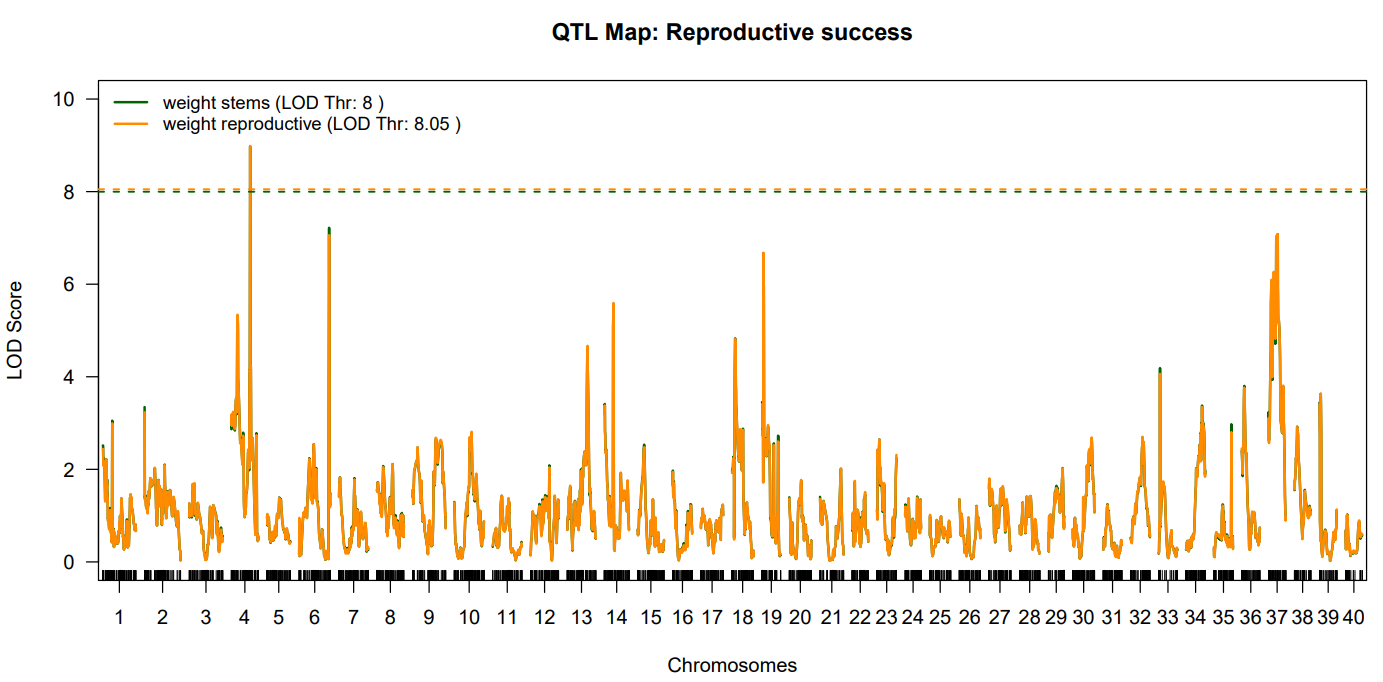
